# Schisandrin B for treatment of male infertility

**DOI:** 10.1101/2020.01.20.912147

**Authors:** Di-Xin Zou, Xue-Dan Meng, Ying Xie, Rui Liu, Jia-Lun Duan, Chun-Jie Bao, Yi-Xuan Liu, Ya-Fei Du, Jia-Rui Xu, Qian Luo, Zhan Zhang, Shuang Ma, Wei-Peng Yang, Rui-Chao Lin, Wan-Liang Lu

## Abstract

The decline of male fertility and its consequences on human populations are important public-health issues. However, there are limited choices for treatment of male infertility. In an attempt to identify a compound that could promote male fertility, we identified and characterized a library of small molecules from an ancient formulation Wuzi Yanzong-Pill, which was used as a folk medicine since the Tang dynasty of China. We found that SB enabled evident repairs in oligoasthenospermia-associated testicular tissue abnormality and in spermatogenesis disruption, resulting in significant improvements of sperm count, mobility, and reproductive ability in oligoasthenospermia mice. Furthermore, SB could alter substantial testicular genes (2033), among which, upregulation of *Fst* while downregulation of *Inhba* involved in reproductive signaling pathway could explain its role in enhancing spermatogenesis. The encouraging preclinical data with pharmacokinetics warranted a rapid development of this new class of therapeutic agent. Our finding provides a strong potent drug for treatment of male infertility.

## Introduction

Infertility is a failure to conceive despite 1 year of regular unprotected intercourse. Based on assessments by the World Health Organization (WHO) and European healthcare bodies over recent years, approximately 8–15% of couples experience infertility. Of these, in 20% of cases the man will be solely responsible; men contribute in an additional 30% of infertility cases [1, 2]. The causes of male infertility vary widely, but oligoasthenospermia is a common cause [3]. Treatment choices for male infertility are limited [4–6]. The ancient formulation Wuzi Yanzong-Pill (WP) has been documented as a medication of male infertility in the book *Xuanjielu* since the Tang dynasty of China. However, its active component(s) are not known. In addition, scientific evaluation of its efficacy and mechanism of action are still lacking. Here, we find that schisandrin B (SB) from WP can be used to treat male infertility, and we further uncover its underlying mechanism of action.

## Results and Discussion

### Identification of schisandrin B for treatment of oligoasthenospermia

To find active components, we had detected >120 types of compounds in WP [7]. We further identified 106 major compounds, and their druggability was evaluated using MedChem Studio (SimulationsPlus, Lancaster, CA, USA) (Supplementary Dataset S1-2; Fig. 1A). Twenty-two compounds had high scores for druggability: 14 phenylpropanoids (including SB), 3 alkaloids, 3 flavonoids and 3 organic acids.

**Fig. 1.**
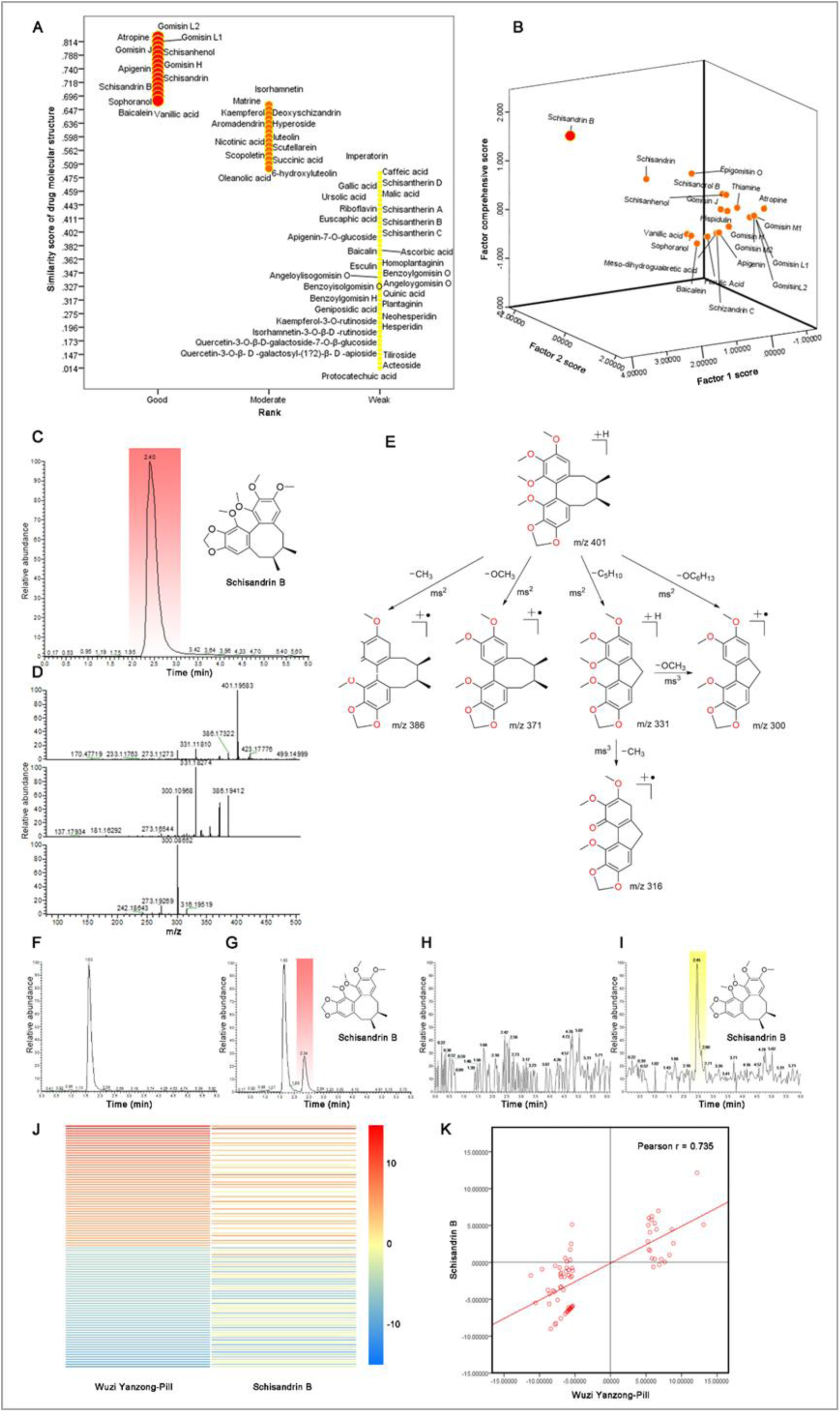
Schisandrin B is identified as a potent agent for treatment of male fertility. ***Notes:*** The studies (A-B) were performed by simulation and statistical analyses in accordance with measurements on the methanol extract of Wuzi Yanzong-Pill (WP) by UPLC-ESI-LTQ-Orbitrap-MS. **A.** Similarity scores of drug molecular structures for 106 major compounds extracted from WP. The study was performed for evaluating the druggability for each component by software of Medchem Studio v3.0 (Simulations Plus, Inc., Lancaster, CA). The result reveals that schisandrin B (SB) along with 21 other components has been listed in the higher score in evaluating the druggability. **B.** Factor comprehensive score of SB among 22 compounds which have higher similarity scores of drug molecular structures. The study was performed for further screening the drug candidate with the Factor Analysis with software of SPSS v 20 (IBM, Armonk, NY). Factor 1, the similarity scores of drug molecular structures; Factor 2, the relative abundances of a compound among 22 compounds extracted from WP. The result indicates that SB has the highest druggability among them in evaluating the factor comprehensive score. The studies (C-I) were analyzed by UPLC-ESI-LTQ-Orbitrap-MS: **C.** Typical total ion chromatogram (TIC) of pure SB; **D.** Triple fragment spectra of pure SB; **E.** Fragmentation pathways of SB; **F.** Typical TIC chromatogram of blank mouse plasma; **G.** Typical TIC chromatogram of mouse plasma after oral administration of SB (20mg/kg) at 3 h; **H.** Typical TIC chromatogram of blank mouse testis; **I.** Typical TIC chromatograms of mouse testis after oral administration of SB (20mg/kg) at 3 h. The studies (J-K) were performed by gene sequence profiling on the testicular samples of oligoasthenospermia mice (OM) after oral administration of SB (20mg/kg/d for 2 weeks; n = 3) or WP (1.56g/kg/d for 2 weeks; n = 3): **J.** Gene heatmaps for the most significant up- and downregulated genes (each 50 genes) in the testicular samples from OM after oral treatment with WP (1.56g/kg/d for 2 weeks; n = 3) or SB (20mg/kg/d for 2 weeks; n = 3). Red color indicates the upregulated genes; Blue color indicates the downregulated genes. **K.** Pearson correlation of the regulated gene log-folds between WP and SB. r represents correlation coefficient.

To screen a drug candidate, the relative content of these components was assessed by factor analysis using SPSS v20 (IBM, Armonk, NY, USA) (Supplementary Dataset S3; Fig. 1B). SB had the highest druggability and was then selected as a candidate.

To determine its oral availability according to site of action, the SB levels in the plasma and in testicular tissues of normal male mice were measured by ultra-performance liquid chromatography-tandem mass spectrometry (UPLC-MS/MS). For the purpose of identification, pure SB was used as the standard reference. The [M+H]^+^ of pure SB had a mass/charge (*m/z*) of 401.19434 (C_23_H_29_O_6_) and eluted at 2.40 min. The representative fragment ions of multistage MS were displayed at *m/z* 401 [M+H]^+^, 386 [M+H-CH_3_]^+**·**^, 370 [M+H-OCH_3_]^+**·**^, 331 [M+H-C_5_H_10_]^+^, 316 [M+H-C_5_H_10_-CH_3_]^+**·**^, and 300 [M+H-OC_6_H_13_]^+**·**^ (Fig. 1C–E).

Three hours after oral administration, SB was identified in the plasma (Fig. 1F, G) and testicular samples (Fig. 1H, I) of mice. SB structure in plasma and testicular samples was identified further by multistage MS as compared with that of pure SB (Supplementary Fig. S1). These results demonstrated the SB availability in plasma and testicular tissue of mice upon oral administration.

Inspired by gene-profiling studies used to screen new chemical chaperones [8–11], we investigated SB involvement in regulation of testicular gene (TG) expression by comparing it with that of WP in an established model of OM [12, 13]. In mice, expression of 100 of the most upregulated and downregulated TGs (50:50) by WP was compared with the corresponding TGs regulated by SB. Heatmap analyses revealed that SB had a similar role in regulation of TG expression as that of WP (Fig. 1J; Supplementary Dataset S4).

Subsequently, Pearson’s correlation analysis was used to assess this similarity quantitatively. The roles of SB and WP were highly correlated: both were involved in regulation of TG expression (*r* = 0.735) (Fig. 1K). Among the multiple active components of WP, SB had a major role, suggesting that SB could be used to treat male infertility.

### Schisandrin B enhances male fertility

To observe the effect of SB on the target tissue, testicular tissues from OM were sampled after oral administration, and histology slices investigated by microscopy. As a pathologic control, the seminiferous tubules of OM were distributed disorderly, and their spermatogenic cells shed severely. In contrast, the seminiferous tubules were distributed uniformly, and injured spermatogenic cells (spermatogonia, spermatocytes, spermatids) were repaired, by treatment with SB or WP (Fig. 2A). As a positive control (testosterone propionate (TP) treatment), the repairing effect on seminiferous tubules and spermatogenic cells was also observed, but to a moderate degree. These results demonstrated that SB and WP could repair damaged seminiferous tubules and spermatogenic cells.

**Fig. 2.**
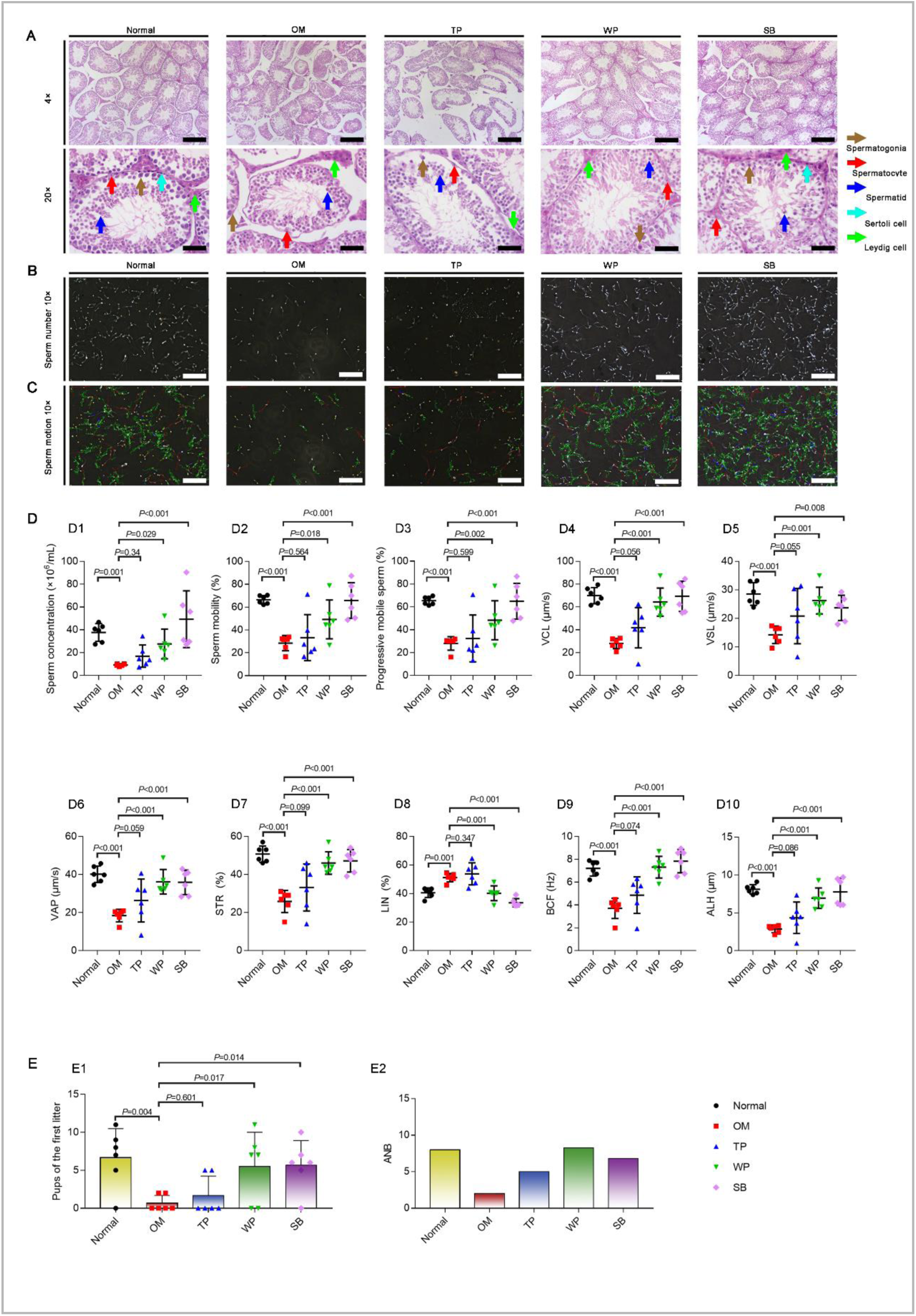
Schisandrin B enables to enhance male fertility in oligoasthenospermia mice. ***Notes:*** The studies (**A-C**) were performed to verify the efficacy of SB in treatment of male fertility, including spermatogenesis, sperm number, and sperm activity in Balb/c mice. **A.** Hematoxylin and eosin staining images of mouse testicular samples. The samples were obtained from normal mice (n=6), OM (n=6), and TP-treated OM (n=6; *i.p.*TP 0.2mg/kg /twice a week for 2 weeks), WP-treated OM (n=6; *i.g.* WP 1.56g/kg/d for 2 weeks) or SB-treated OM (n=6; *i.g.* SB 20mg/kg/d for 2 weeks). *i.g.*, intragastric administration; OM, oligoasthenospermia mice; SB, schisandrin B; TP, testosterone propionate; WP, Wuzi Yanzong-Pill. Scale bar, 200 μm. Brown arrow indicates spermatogonia; red arrow indicates spermatocyte; blue arrow indicates spermatid; cyan arrow indicates sertoli cells; and green arrow indicates leydig cells. The results demonstrate that SB enables to repair the disrupted spermatogenesis of OM. **B.** Sperm number images of mouse cauda epididymidis samples under Suiplus Semen Analysis Automatic Detection System (Suiplus, BeiJing, China). The samples were obtained from normal mice (n=6), OM (n=6), and TP-treated OM (n=6; *i.p.* TP 0.2mg/kg /twice a week for 2 weeks), WP-treated OM (n=6; *i.g.* WP 1.56g/kg/d for 2 weeks) or SB-treated OM (n = 6; *i.g.* SB 20mg/kg/d for 2 weeks). The results directly demonstrate that SB enable to increase the sperm number of OM. The dynamic videos of this study are available in the **Supplementary Video 1-Video 5**. **C.** Sperm motion track images of mouse cauda epididymidis samples under Suiplus Semen Analysis Automatic Detection System (Suiplus). The samples were obtained from the same as above (Fig. 2B). The observation displays that SB increases the sperm mobile activity of OM. The analyses were performed for evaluating the quality of sperms in OM after oral treatment with SB. **D.** Quality of spermatogenesis. **D1,** sperm concentrations; **D2,** sperm mobility; **D3,** progressive mobile sperms; **D4,** curvilinear velocity (VCL); **D5,** straight-line velocity (VSL); **D6,** average path velocity (VAP); **D7,** straightness (STR); **D8,** linearity (LIN); **D9,** beat cross frequency (BCF); **D10**, amplitude of lateral head displacement (ALH). The studies (**E-F**) were performed for evaluating the male reproductive ability by comparing the number of pups in the first litter of female mice, and the average number of births (ANB; = total number of births/birth females). Each male mouse was placed in one cage, and mated with two females. **E.** Efficacy in enhancing reproductive ability (n=3). **E1,** pups in the first litter of female mice; **E2,** average number of births (ANB; = total number of births/birth females). These data demonstrate that SB significantly increase male reproductive ability, leading to an enhanced ability of male mice to make female mice pregnant and the mean number of offspring.

To observe the effect of SB on sperm, sperm samples were collected from OM after oral administration, and the number and movement of sperm investigated by a computer-aided sperm-analysis system. As a pathologic control, the sperm number of OM was decreased significantly, and sperm movement was inactive or less motile. In contrast, sperm number was increased significantly and sperm movement was very active in OM after oral treatment with SB or WP, indicating a similar number and motility of sperm to those of normal mice (Fig. 2B, C; Supplementary Video 1–5). As a positive control after TP treatment, the repairing effects on the number and movement of sperm were exhibited to a moderate degree. These results revealed that SB and WP could increase the number and motility of sperm.

Subsequently, five visual fields from each video were recorded for quantitative evaluation of sperm parameters. The established OM model met the diagnostic criteria set by the WHO [14] for oligozoospermia (sperm concentration <20×10^6^/mL) and asthenospermia (sperm mobility <40%; progressive mobile sperm <32%) (Fig. 2D1–3). Furthermore, the sperm-activity parameters of OM were also decreased significantly, *i.e.*, sperm-motion velocities (curvilinear velocity (VCL) (Fig. 2D4); straight-line velocity (VSL) (Fig. 2D5); average path velocity (VAP) (Fig. 2D6), sperm-motion locus (straightness (STR) (Fig. 2D7)), and dynamic parameters (beat cross frequency (BCF) (Fig. 2D9); amplitude of lateral head displacement (ALH) (Fig. 2D10)), but not an increase in linearity (LIN) (Fig. 2D8). After treatment with SB or WP, all sperm parameters were increased significantly to the levels observed in normal mice (Fig. 2D1–10; Supplementary Dataset S5). As a positive control (TP treatment), the repairing effect on OM was moderate. These results provided robust evidence that SB and WP could increase the number and quality of sperm in OM.

To investigate the reproductive ability, male mice were mated with female mice at a 1:2 ratio (Supplementary Dataset S6). For verification purposes, various experimental and control groups were designed. Normal female mice were included in all groups. Male mice were normal mice, OM, OM after treatment with SB, WP or TP for 2 weeks, respectively. The number of pups in the first litter of OM was diminished significantly as compared with that of normal male mice. In contrast, the number of pups in the first litter was increased markedly in the group of OM after treatment with SB or WP (Fig. 2E1). Furthermore, the number of pups in the first litter of OM after treatment with SB or WP was very close to that of normal mice. In addition, TP exhibited only slight efficacy upon treatment in OM. Treatment of OM with SB or WP increased the average number of births (ANB) significantly, showing fertility close to that of normal mice, respectively (Fig. 2E2). Hence, SB could be used to treat to treat infertility in OM.

### Schisandrin B regulates testicular genes in the reproductive signaling pathway

We wished to reveal the mechanism of action of SB. Hence, RNA sequencing was done on the testicular tissues of OM and after oral treatment with SB for 2 weeks. Volcano plots showed that SB treatment resulted in significantly changed expression of 2033 genes (836 upregulated; 1197 downregulated) (Fig. 3A).

**Fig. 3.**
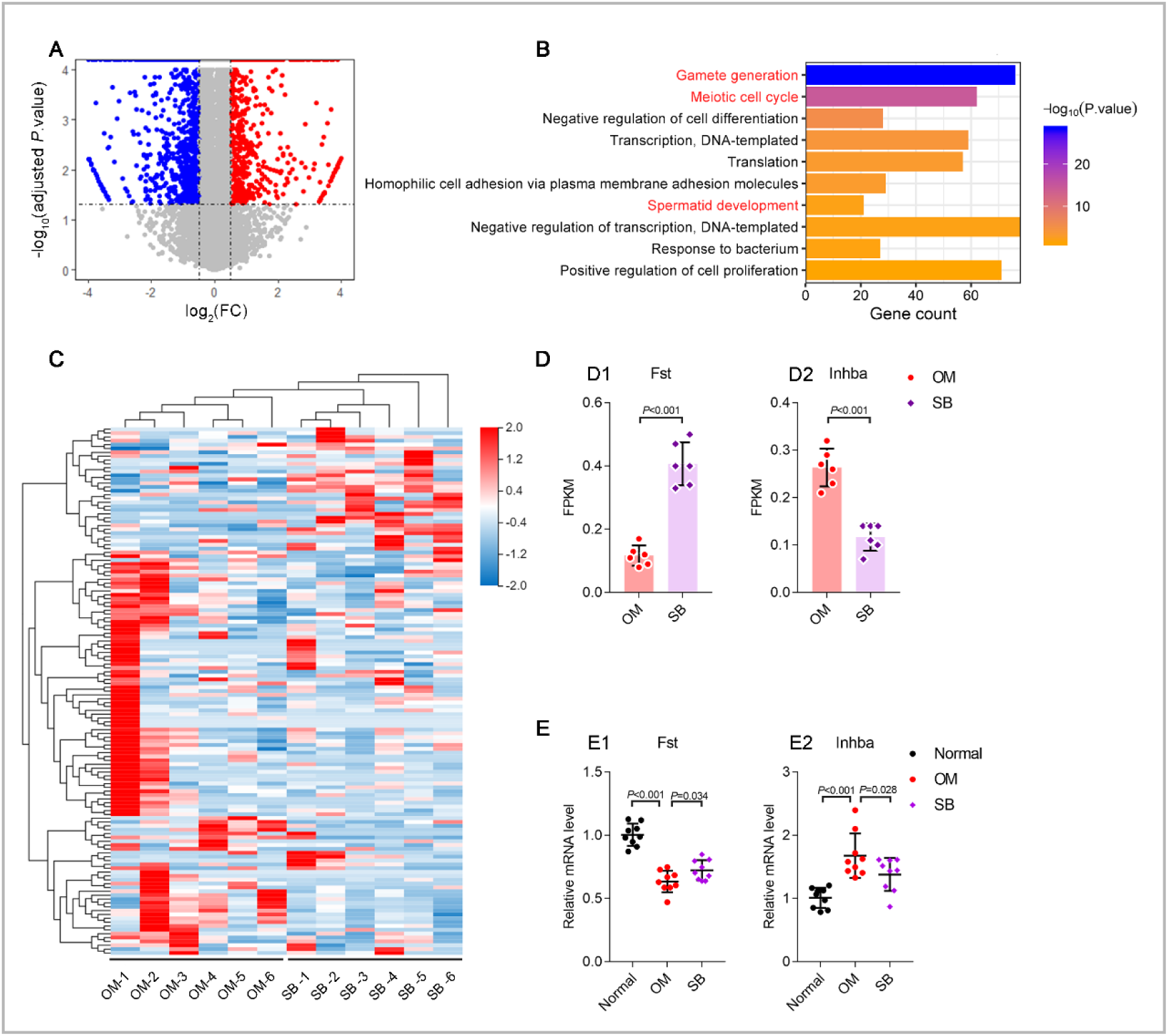
Schisandrin B regulates testicular gene expressions of *Fst* and *Inhba* in the reproductive pathway. ***Notes:*** The studies (**A-D**) were performed to reveal the regulated functional genes by schisandrin B (SB) by using gene sequencing on testicular samples of oligoasthenospermia mice (OM) and on those of SB-treated OM (*i.g.* SB 20mg/kg/d for 2 weeks; n = 6): **A.** Volcano plot of SB-mediated changes of testicular genes by software of Dr Tom (v2.0, Beijing Genomics Institute, BGI Shenzhen, China).The samples were obtained from OM (n=6), and SB-treated OM (*i.g.* SB 20mg/kg/d for 2 weeks; n = 6). SB-changed genes were identified with two threshold criteria: fold up- or down regulation in SB -treated mice of |log ^(FC)^| > 0.58, and adjusted *P* value of less than 0.05. The results reveal that, after oral administration of SB in OM, it significantly up-regulates 836 genes, while down-regulates 1197 genes. **B.** Top ten GO pathways involved in above changed genes by GO analysis. GO enrichment was performed on above regulated-genes (totally 2033 genes) by using software of Dr Tom. The results indicate that, among top ten GO pathways, three reproductive pathways, including gamete generation, meiotic cell cycle, and spermatid development, are involved in the gene regulations by SB. Besides, a number of 137 genes are included in the reproductive pathways. **C.** Gene heatmap for SB-regulated testicular genes (n=137 genes) by using software of Dr Tom. The results indicate that oral administration of SB significantly alters testicular gene signature in OM. Furthermore, it reveals that *Fst* gene is the mostly regulated functional gene in viewing the absolute fold change or adjusted *P* value. ***D.*** *Fst* and *Inhba* gene expressions in testicular samples of OM after oral treatment of SB. **D1,** *Fst* gene expression level; **D2,** *Inhba* gene expression level. FPKM represents the fragments per kilobase per million mapped fragments. The results reveal that SB significantly up-regulates *Fst* gene while down-regulates *Inhba* gene in OM after oral treatment of SB. The studies were performed for verifying the regulated mRNA levels of *Fst* and *Inhba* gene expressions by RT-qPCR in testicular samples of OM after oral treatment of SB: **E.** mRNA levels of *Fst* and *Inhba* in testicular samples of OM after oral treatment of SB. **E1,** *Fst* mRNA expression; **E2,** *Inhba* mRNA expression. The samples were obtained from normal mice (n=3), OM (n=3), and SB treated-OM (*i.g.* SB 20mg/kg/d for 2 weeks; n = 3).The results exhibit that oral treatment of SB significantly increases *Fst* mRNA expression while decreases *Inhba* mRNA expression in testicular tissue of OM, indicating that SB could treat oligoasthenospermia by regulating expressions of *Fst* and *Inhba* genes.

To further identify the relevant biologic pathways, the Gene Ontology (GO) database was applied. Three reproductive pathways were involved in the top-10 pathways: gamete generation, meiotic cell cycle, and spermatid development (Fig. 3B; Supplementary Dataset S7).

In these three pathways, 137 genes whose expression was regulated significantly were plotted into heatmaps. We found an obvious difference in the TG signature between OM and SB-treated OM (Fig. 3C; Supplementary Dataset S8).

Of the enriched 137 genes in the three reproductive pathways, the predicted genes and pseudogenes were assigned “low priority” by searching gene databases in the public domain (National Center for Biotechnology Information, Ensembl, Gene Cards) and, finally, 19 candidate genes were obtained.

Oligoasthenospermia can manifest as upregulation or downregulation of TG expression, which correspond to positive or negative fold-changes. Hence, TGs with the most significant absolute fold-changes were studied. Accordingly, the absolute fold-changes of the 19 TGs mentioned above were calculated by comparing SB-treated mice with untreated mice, and processed by statistical analyses with adjusted *P*-values. *Fst* was the most regulated TG, showing remarkable upregulation of expression upon oral administration of SB (Supplementary Dataset S8).

We further investigated expression of *Inhba* in testicular tissues in OM treated and un-treated by SB. As a pathologic control, mRNA sequencing showed that *Fst* was expressed at a low level whereas *Inhba* was expressed at a high level in the testicular tissues of OM. Conversely, *Fst* expression was upregulated significantly whereas *Inhba* expression was downregulated markedly in the testicular tissue of OM after SB treatment (Fig. 3D, Supplementary Dataset S9). Furthermore, upregulation of *Fst* expression and downregulation of *Inhba* expression were verified by quantitative reverse transcription-polymerase chain reaction (RT-qPCR) (Fig. 4E, Supplementary Dataset S10).

**Fig. 4.**
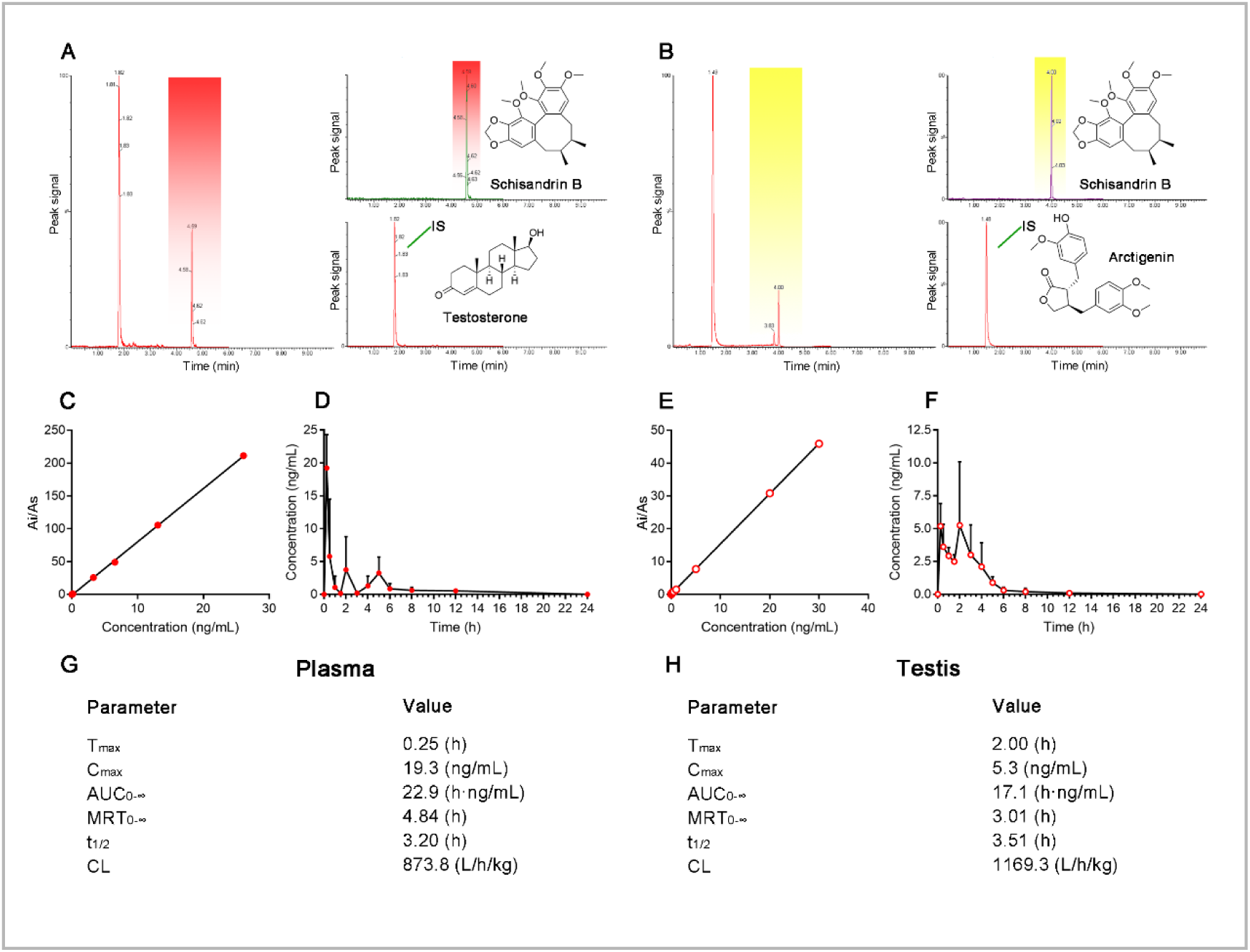
Pharmacokinetics of schisandrin B in plasma and in testicular tissue in normal mice after oral administration. ***Notes:*** _The studies (**A-F**) were performed to establish analysis method for the separation and determination of schisandrin B (SB) in plasma and testicular tissue of normal mice._ **A.** Typical multiple reaction monitoring (MRM) chromatogram of mouse plasma after oral administration of SB (20mg/kg) at 3 h. Internal standard (IS) added in plasma, testosterone. The results display that the chromatogram peaks of SB and IS in plasma appear at 4.59 min and 1.82 min, respectively, and there are no interfering peaks in the chromatogram, demonstrating a good specificity of analysis method for plasma samples. **B.** Typical MRM chromatogram of mouse testes after oral administration of SB (20mg/kg) at 3 h. IS added in testicular tissue, arctigenin. The results display that the chromatogram peaks of SB and IS in testicular tissue appear at 4.00 min and 1.48 min, and there are no interfering peaks in the chromatogram, demonstrating a good specificity of analysis method for testicular samples. **C.** Calibration curve of SB in mice plasma. Ai indicates peak area of SB; and As indicates peak area of IS (testosterone) in plasma samples. The results show that the peaks and concentration of SB are linearly correlated in the range of 0.07 to 26.09 ng/mL in plasma. **D.** Mean plasma concentration-time curve of SB. The sampling was performed at 0 min, 15 min, 30 min, 1 h, 1.5 h, 2 h, 3 h, 4 h, 5 h, 6 h, 8 h, 12 h and 24h before and oral administration of SB (20mg/kg) (n= 5). The results show that SB can be rapidly absorbed into blood, and has triple absorption peaks, suggesting a hepato-intestinal circulation pathway during absorption and metabolism. **E.** Calibration curve of SB in testicular tissue. Ai indicates peak area of SB; and As indicates peak area of IS in testicular samples. The results show that the peaks and concentration of SB are linearly correlated in the range of 0.10 to 30.00 ng/mL in testicular tissue. **F.** Mean testicular concentration-time curve of SB. The sampling was performed at 0 min, 15 min, 30 min, 1 h, 1.5 h, 2 h, 3 h, 4 h, 5 h, 6 h, 8 h, 12 h and 24h before and oral administration of SB (20mg/kg) (n= 5). The results show that SB reaches the testicular tissue rapidly after absorption, demonstrating that SB is able to reach the action site. The studies (**G-H**) were performed to calculate the pharmacokinetic parameters in mice plasma and in testicular tissues by software of DAS v3.2 (China State Drug Administration, Shanghai, China). **G.** Major pharmacokinetic parameters in mice plasma after oral administration of SB. **H.** Major pharmacokinetic parameters in mice testicular tissue after oral administration of SB.

These results indicated that SB could treat oligoasthenospermia by regulating expression of *Fst and Inhba*. We postulated a mechanism. Briefly, overexpression of activin-A protein (encoded by *Inhba*) can inhibit the growth of testicular spermatogenic cells and induces apoptosis in spermatogenic cells. Conversely, follistatin protein (encoded by *Fst*) promotes the growth and development of spermatogenic cells by blocking the action of activin-A protein [15–19]. There is serious disorder in the testicular spermatogenic cells of patients with oligoasthenozoospermia, and the major reason is activin-A overexpression, which leads to spermatogenic blockage [20–22]. Therefore, upregulated *Fst* expression would increase follistatin expression, which enables blockade of the action of overexpressed activin-A, thereby repairing spermatogenic blockage [23–25]. Moreover, downregulation of *Inhba* expression could contribute directly to the decrease in activin A-expression, thereby attenuating spermatogenic blockage. Therefore, we revealed that upregulation of *Fst* expression and downregulation of *Inhba* expression could promote spermatogenesis by inhibiting apoptosis of spermatogenic cells.

### Preclinical pharmacokinetic evaluation on schisandrin B

With regard to potential clinical use [26–28], we investigated the pharmacokinetics of SB in normal mice after oral administration. Plasma and testicular concentrations of SB were measured by LC-MS/MS. SB and the internal standard (IS) in plasma were eluted at 4.59 min and 1.82 min (Fig. 4A), and those in testicular samples were eluted at 4.00 min and 1.48 min (Fig. 4B), respectively. There were no interfering peaks in the chromatograms of plasma and testicular samples, indicating suitable specificity for measurement. The calibration curves of SB in plasma and testicular samples exhibited appropriate linearity in a wide concentration range, respectively (Fig. 4C, E). Besides, the measurement was validated [29–30] and consisted of: correlation coefficients, linear ranges, and lower limit of quantifications (LLoQs)(Supplementary Dataset S11); intra-/inter-day precisions and accuracies (Supplementary Dataset S12); recovery stability; measurement stability (Supplementary Dataset S13 and S14).

After oral administration, plasma and testicular concentration–time profiles for SB were plotted (Fig. 4D, F; Supplementary Dataset S15, S16), and the corresponding pharmacokinetic parameters calculated (Supplementary Dataset S17 and S18), respectively. The major pharmacokinetic parameters of SB were evaluated: time to reach maximum concentration (T_max_), maximum concentration (C_max_), area under the concentration–time curve (AUC_0–∞_), mean residence time (MRT_0–∞_), half-life of elimination (t_1/2_), and clearance (CL).

Accordingly, SB parameters in plasma were: T_max_= 15 min; C_max_= 19.3 ng/mL; AUC_0–∞_= 22.9 h.ng/mL; MRT_0–∞_= 4.84 h; CL = 873.8 L/h/kg; t_1/2_ = 3.2 h (Fig. 4G). Plasma concentration–time profiles revealed re-absorption in the gastrointestinal tract due to three concentration peaks, suggesting a hepatointestinal circulation. Assessment of plasma parameters demonstrated that oral administration led to rapid absorption and an effective exposure of SB in blood. Furthermore, SB could be eliminated from blood within 1 day (7-fold half-life washing-out period about 21 h).

SB parameters in testicular tissue were: T_max_= 2.0 h; C_max_ = 5.3 ng/mL; AUC_0–∞_ = 17.1 h.ng/mL; MRT_0–∞_ = 3.0 h; CL = 1169.3 L/h/kg; t_1/2_ = 3.5 h (Fig. 4H). Concentration–time profiles from testicular tissue showed two concentration peaks (minor peak at 15 min and maximum peak at 3 h) suggesting that, after absorption, SB was distributed effectively into testicular tissue but with a delay. Nonetheless, there was effective testicular exposure of SB as well. Hence, SB in testicular tissue had comparable pharmacokinetic behavior to that in blood.

### Conclusion

In summary, SB could be used to treat infertility in male mice. The present study involved four main stages. First, SB was detected from 106 compounds of an ancient formulation (WP) to treat male infertility by study of: similarity in drug structure; SB availability after oral administration; regulation of TG expression by comparing SB with WP. Second, the efficacy of SB was assessed to treat male infertility by studying: repair of damaged seminiferous tubules and spermatogenic cells by pathologic staining; enhancement of sperm-number and sperm-motility parameters using a computer-aided sperm-analysis system; and improvement of reproductive ability. Third, 2033 differentially expressed genes induced by SB were revealed by RNA sequencing. Use of the GO database showed that three reproductive pathways were enriched in gene regulation: gamete generation, meiotic cell cycle and spermatid development. We found that upregulation of *Fst* expression and downregulation of *inhba* expression interacted to repair spermatogenesis. This phenomenon could be explained by the fact that follistatin (encoded by *Fst*) promotes the growth and development of spermatogenic cells by blocking the induced apoptosis of spermatogenic cells by activin-A (encoded by *Inhba*). Fourth, pharmacokinetic studies demonstrated that SB could be absorbed rapidly after oral administration, and became fully available at the intended action site, indicating a remarkable potential for clinical application.

In conclusion, SB enables the repairs of spermatogenesis arrest and male infertility. The action mechanism could be explained by the repaired spermatogenesis via upregulation of *Fst* while downregulation of *Inhba* genes involved in the reproductive signaling pathway. Our study provides a promising drug for treatment of male infertility and a novel strategy for discovery of new small-molecule drugs from vast plant-based medicinal resources.

## Materials and Methods

### Ethical approval of the study protocol

All procedures involving the care and handing of animals were carried out with approval of the Authorities for Laboratory Animal Care of Peking University (Beijing, China; LA2018330).

### Reagents

Busulfan was purchased from Macklin Biochemicals (Shanghai, China). Pure SB was obtained from the National Institutes for Food and Drug Control (Beijing, China; purity >98%; HPLC grade). TP (injection) was purchased from Shanghai General Pharmaceuticals (Shanghai, China). *Fructus lych, semen cuscutae* (fried), *fructus rubi*, *fructus Schisandrae chinensis* (steamed), and *semen plantaginis* (fried with salt) were from Beijing Tong Ren Tang Group (Beijing, China). Acetonitrile, methanol, ethanol, and formic acid were purchased from Fisher Scientific (Fair Lawn, NJ, USA; LC-MS grade). Water was purified by a Milli-Q™ ultraviolet purification system (Millipore, Bedford, MA, USA). Dimethyl sulfoxide was obtained from Sigma–Aldrich (Saint Louis, MO, USA). Polyethylene glycol (PEG)_400_ was purchased from Harveybio Gene Technology (Beijing, China). Sperm Culture Medium 199 (M199) was obtained from Thermo Scientific (Waltham, MA, USA). Bovine serum albumin was purchased from Solarbio (Beijing, China). PCR primers were obtained from Tsingke Biological Technology (Beijing, China). All other chemicals were from commercial sources.

### Animals

Male and female Balb/c mice (10 weeks; 20.0 ± 2 g) were obtained from the Department of Laboratory Animal Science, Peking University Health Science Center (order ID: SCXK (jing) 2016-0010). Each animal was housed in an individual cage at controlled temperature (25±1°C) and humidity (55±5%) and exposed to a 12-h light–dark cycle (7 pm to 7am). Animals had free access to food (regular chow comprising 5% fat, 53% carbohydrate and 23% protein) and water unless indicated otherwise.

### Ancient formulation for treatment of male infertility

The ancient formulation consisted of *Fructus lych*, *semen cuscutae* (fried), *fructus rubi*, *fructus Schisandrae chinensis* (steamed), and *semen slantaginis* (fried with salt). The medicinal materials were weighed, mixed (8:8:4:2:1, *w/w*) and crushed to powder (mesh size = 40). Then, they were immersed in a 10-fold volume of water for 1 h at 100°C. After boiling, heating was continued until the volume was reduced to fivefold volume as compared with the original one. The mixture was filtered immediately through gauze, concentrated to 1 g of crude drug per mL, and freeze-dried to become powders. Finally, refined honey (85 g) was mixed with freeze-dried powders (100 g) to make pellets of WP for experimental use.

### Extraction of compounds and prediction of druggability

WP (250 mg) were extracted with 50 mL of methanol with the aid of ultrasound for 60 min. Extracts were centrifuged at 10,000 revolution per minute (rpm) for 15 min at 4°C. The supernatant was collected and passed through a filter (0.22 μm). The filtrate was collected for UPLC coupled with electrospray ionization-linear ion trap-Orbitrap tandem mass spectrometry (UPLC-ESI-LTQ-Orbitrap-MS) measurement. Based on chromatographic data, 106 major compounds were identified in the WP extract, including organic acids, flavonoids, phenylpropanoids, alkaloids and terpenoids. To predict the most promising drug candidate, the druggability was evaluated on the 106 compounds extracted from WP using MedChem Studio v3.0 (Simulations Plus). Compounds with a good drug-similarity score (i.e., druggability) were selected for further consideration by combination with drug contents in the extract. Based on the druggability and drug content (which was indicated in the corresponding peak relative abundance in the chromatogram of UPLC-ESI-LTQ-Orbitrap-MS), drug candidates were selected preliminarily in accordance with the highest factor comprehensive score using factor analysis employing SPSS v20.0 (IBM).

### Availability of SB by action site

#### Measurement

To ascertain if the drug candidate (SB) could be absorbed in blood or reach the action site, SB in plasma and testicular tissue (action site of drugs for male infertility) was measured by UPLC-MS/MS after oral administration of SB in male mice. Analyses were undertaken on a UPLC system (Acquity™ UPLC Ӏ-Class system; Waters, Milford, MA, USA) consisting of an auto-sampler, quaternary pump, and column oven. A C18 reverse-phase column (Acquity UPLC BEH, 100 × 2.1 mm, 1.7 μm, 130 Å) was used to separate samples. The mobile phase was 0.1% formic acid in water (A) and acetonitrile (B). The gradient elution was: 0 min 78% B, 1 min 78% B, 4 min 60% B, and 6 min 78% B. Samples were kept in the autosampler at 4°C until measurement. The column was maintained at 40°C, the flow rate was 0.3 mL/min, and injection volume was 5 μL. The UPLC was connected to a mass spectrometer (LTQ/Orbitrap; Thermo Scientific) *via* an ESI interface. The effluent was split at a ratio of approximately 3:1 (*v/v*) before entering the ESI source. Positive-ion mode was used, and operation parameters were: capillary voltage, 25 V; electrospray voltage, 4.0 kV; capillary temperature, 350°C; sheath gas, 30 (arbitrary units); auxiliary gas, 5 (arbitrary units); tube lens, 110 V. High-resolution full scan was used to scan samples with a resolution of 30,000 and a scanning mass range of 100 to 500 amu. Data-dependent scan was used to scan secondary and tertiary mass spectra, and the three peaks with the highest abundance in the upper MS level were selected for collision-induced fragmentation scanning. The normalized collision energy was set to 35%. To avoid many repeated data acquisitions on the same sample, dynamic exclusion was used for data collection with an exclusion duration of 60 s and the repeat count was set at 5 with a dynamic repeat time at 30 s. An external calibration for mass accuracy was carried out before the analysis. The measured masses were within 5 ppm of the theoretical masses. Data analyses were processed using a Xcaliber 2.1 workstation (Thermo Fisher Scientific). Meanwhile, pure SB (5 mg) was dissolved in 10 mL of methanol, passed through a filter (0.22 μm), and used as the reference for analyses.

#### Dosing

Pure SB (1 mg/mL) was dissolved in a mixture of ethanol, PEG_400_ and 0.5% sodium carboxymethyl cellulose (CMC-Na) (1:1:1, *v/v/v*) for oral administration. Male mice were divided randomly into two groups of three. In the treatment group, each mouse was administered SB (20 mg/kg, i.g.). In the blank control group, each mouse was given physiologic saline (PS).

#### Sampling

Three hours after dosing, venous blood (0.75 mL) was sampled and centrifuged at 5000 rpm for 10 min at 4°C. Then,+ 200 μL plasma was transferred, added to 600 μL of acetonitrile, vortex-mixed (120 s), and centrifuged (13,000 rpm, 10 min, 4°C) to remove proteins. The supernatant was evaporated at 25°C by a CentriVap™ centrifugal thickener (Labconco, Kansas City, MO, USA). The residues were dissolved in 200 μL of methanol, and centrifuged (13,000 rpm, 10 min, 4°C). The resultant supernatant was injected into the UPLC-MS/MS system.

The animals were sacrificed. The testicular tissues were collected on an ice plate at the same time as blood sampling. Next, they were washed with PS, drained with filter paper and weighed. PS (1:4, *w/v*) was added and the testicular tissue homogenized. One milliliter of testicular-tissue homogenate was centrifuged at 5,000 rpm for 10 min at 4°C. The supernatant (200 μL) was collected, and 400 μL of acetonitrile added, followed by vortex-mixing (120 s) and centrifugation at 13,000 rpm for 15 min at 4°C. Finally, the resultant supernatant was injected into the UPLC-MS/MS system.

### Involvement of SB in regulation of TG expression

To investigate SB involvement in regulating TG expression by comparing it with that of WPs, OM models were induced by intraperitoneal injection of busulfan (20 mg/kg dissolved in sterile dimethyl sulfoxide). OM were divided into three groups of three, and treated once daily with PS, SB (20 mg/kg/d, i.g.) or WP (1.56 g/kg/d, dissolved in 0.5% CMC-Na). After 2-week treatment, the testes of all animals were dissected, frozen immediately in liquid nitrogen, and stored at −80°C for gene sequencing.

To extract total RNA, 200 mg of the testicular sample was processed using TRIzol by following manufacturer (Invitrogen, Carlsbad, CA, USA) protocols and its expression determined using a 2100 Bioanalyzer (Agilent Technologies, Santa Clara, VCA, USA). Only qualified RNAs from testicular samples were used for construction of cDNA libraries. Preparation and sequencing of cDNA libraries were undertaken by the BGI Genomics Co., Ltd. (BGI, Shenzhen, China) using the BGISEQ-500 platform.

To analyze RNA-sequencing data, initially raw reads were excluded if they contained >10% nitrogen, or were adapter or low-quality reads, using SOAPnuke v1.5.2 by BGI. High-quality reads were aligned to the reference genome (mouse) using HISAT v2.0.4 and gene expression was normalized to fragments per kilobase of exon model per million mapped reads (FPKM) using RSEM v1.2.12 by BGI. Normalized FPKM expression was analyzed using Dr Tom v2.0 by BGI to identify differentially expressed genes. The 100 most-regulated genes in the testes of WP-treated OM, and their corresponding gene expression fold-changes in the testes of SB-treated OM, were selected as typical gene signatures to compare the gene profile. The 100 most-regulated TGs consisted of 50 upregulated genes and 50 downregulated TGs. The comparison of gene heatmaps between WP and SB was made by Dr Tom v2.0 by BGI. Besides, Pearson’s correlation analysis was applied to quantitatively analyze the similarity in gene expressions in the testes of OM after oral treatment with SB or WP.

### Spermatogenesis repair by SB

#### Dosing

OM were divided into four groups of six and treated with SB (20 mg/kg/d, once daily, i.g.), WP (1.56 g/kg/d, once daily, i.g.), TP (0.2 mg/kg/twice a week, i.p.) or PS (14 mL/kg/d, i.g.), respectively. Normal mice (n = 6) were given PS (14 mL/kg/d, i.g.). All animals were given these agents consecutively for 2 weeks and the observations shown below made.

#### Sampling of testicular tissue

After 2-week treatment, each mouse was anesthetized with diethyl ether. Tissue from the left testes was harvested, stored in 10% formalin, and paraffin-embedded for staining (hematoxylin and eosin).

#### Sperm sampling

After 2-week treatment, the limbs of each mouse (under anesthesia) was fixed on a thermostatic hot plate (37°C). The left epididymis was dissected promptly, cleaned with PS (37°C) and transferred immediately to 0.5 g of bovine serum albumin per μL of medium 199 (1 mL, 37°C). Tissue was cut into pieces by scissors. Sperm was allowed to flow out of the tissue, and then placed in an incubator in an atmosphere of 5% CO_2_ for 3 min at 37°C. After incubation, the suspension was mixed homogeneously by a pipette, then 10 μL of sperm suspension was placed on a semen-counting slide (Yulu Optics, Nanjing, China). This slide had a depth of 0.01 mm, and enabled unimpeded movement of sperm.

#### Microscopic observation of sperm

The sperm-counting slide was placed under a phase-contrast microscope (E200; Nikon, Tokyo, Japan). A video was recorded by a semen analysis automatic detection system (Suiplus; Beijing, China). Five visual fields were taken from each counting slide for observation. The movement track, morphology, concentration and number of sperm were observed, and recorded for qualitative evaluations and parameter evaluations.

#### Quality parameters of sperm

IVOS software (Hamilton Thorne Biosciences, Beverly, MA, USA) in the semen analysis automatic detection system (Suiplus) was used to evaluate the quality parameters of sperm. The parameters were sperm concentration, sperm mobility, progressive mobile sperm, sperm motion velocity (VCL, VSL, VAP), sperm-motion locus (STR, LIN) and dynamic parameters of sperm movement (BCF, ALH).

### Efficacy of SB in enhancing male reproductive ability

#### Dosing

OM were divided into four groups of three and treated with SB (20 mg/kg/d, once daily, i.g), WP (1.56 g/kg/d, once daily, i.g.), TP (0.2 mg/kg, twice a week, i.p.) and PS (14 mL/kg/d, once daily, i.g.), respectively. Normal mice (n = 3) were given PS (14 mL/kg/d, i.g.). All animals were given these agent consecutively for 2 weeks.

#### Reproductive ability

After 2-week treatment, each male mouse was mated with females at a 1:2 ratio. Mating mice were placed in one cage for 10 days (two sex cycles of females). Females were examined for pudendal embolus each morning at 8:30. The plugged female was removed from the cage immediately. If there was no sign of intercourse, the female(s) and male mice were placed in the same cage continuously until the end of the tenth day. After 10 days, female mice were separated from the male mouse, and observed for 40 days. The total number of pups in the first litter for a pregnant female, and the number of non-pregnant females, was recorded. The ANB was calculated using the formula:

ANB = total number of births/number of females who gave birth

### Gene profiling and biologic pathways regulated by SB

#### Dosing and sampling

OM were divided into two groups of six and treated once daily with SB (20 mg/kg/d, i.g.) or PS (14 mL/kg/d, i.g.), respectively. After 2-week treatment, each mouse was anesthetized with diethyl ether. Testicular tissue was frozen immediately in liquid nitrogen and stored at −80°C for further analyses. Normal male mice were included as a blank control (n = 6). After experimentation, mice were sacrificed by cervical dislocation.

#### Gene profiling and GO analyses

Frozen testicular samples (n = 6) from SB-treated or non-SB-treated OM were used for RNA sequencing. Extraction of total RNA and data analyses were done as described above. Furthermore, functional annotation of differentially expressed genes in the GO database was applied using Dr Tom v2.0 by BGI.

### RT-qPCR verification

Frozen testicular samples (n = 3) from normal mice, OM, or SB-treated OM were used for RT-qPCR. Total RNA was extracted using a TRIzol Plus RNA Purification kit (Invitrogen), and analyzed (excitation wavelength =260 nm, emission wavelength = 280 nm) using a spectrophotometer (Nano300; Allsheng, Hangzhou, China).

cDNA was reverse-transcribed from 1 μg of total RNA using PrimeScript RT reagent (TaKaRa Biotechnology, Shiga, Japan), and 10 ng of cDNA was analyzed using SYBR Premix Ex Taq II (TaKaRa Biotechnology) on a CFX Connect TM Real-Time PCR Detection System (Bio-Rad Laboratories, Hercules, CA, USA). Each sample was tested in triplicate. Glyceraldehyde 3-phosphate dehydrogenase (GAPDH) was used as an internal control. Relative quantification of genes of interest was done using the 2^−ΔΔct^ method. Primer sequences used for RT-qPCR (forward and reverse, respectively) were 5′-TGCTCTTCTGGCGTGCTTCTTG-3′ and 5′-TGTAGTCCTGGTCTTCCTCCTCCT-3′ for the *Fst* primer; 5′-GTCCTCGCTCTCCTTCCACTCAA-3′ and 5′-AGCAGCCACACTCCTCCACAAT-3′ for the *Inhba primer*; 5′-AGAAGGTGGTGAAGCAGGCATCT-3′ and 5′-CGGCATCGAAGGTGGAAGAGTG-3′ for the *GAPDH* primer.

### Pharmacokinetics

#### Working solutions

Pure SB was weighed accurately and dissolved in methanol to prepare working standard solutions (0.05–30.0 ng/mL). The IS solution of testosterone (25.0 ng/mL) and arctigenin (25.0 ng/mL) was prepared similarly for SB measurements in plasma and testicular tissue, respectively. All solutions were stored at 4°C before use.

#### Sampling of blank plasma and testicular tissue

Normal mice (n = 15) were anesthetized with diethyl ether. Aliquots of venous blood (0.75 mL) were sampled, centrifuged at 5000 rpm for 10 min at 4°C to obtain plasma, and stored at −80°C until use. Animals were sacrificed, testicular tissues were collected on an ice plate at the same time of blood sampling, and frozen immediately at −80°C for use.

#### Calibration curves and quality control (QC)

Calibration curves and QC samples for SB in plasma and testicular tissue were prepared in duplicate to evaluate the precision, accuracy, stability and recovery of our analytical method. The handling procedures are described below.

#### Plasma handling

Plasma was thawed at 4°C for ∼30 min and vortex-mixed for 30 s. Plasma (200 μL) was vortex-mixed with 60 μL of a working solution of SB for 30 s, added to 60 μL of IS solution (testosterone) and vortex-mixed for 30 s. Then, 600 μL of acetonitrile was added, followed by vortex-mixing for 120 s, and centrifugation at 13,000 rpm for 10 min at 4°C. The resultant supernatant was injected into the UPLC-MS/MS system. SB concentrations for calibration curves were prepared at 0.05, 0.10, 3.0, 6.0, 12.0 and 25.0 ng/mL in plasma, whereas those for QC analyses were prepared at 1.0, 10.0 and 20.0 ng/mL in plasma. In these samples, the IS concentration was 25.0 ng/mL.

#### Handling of testicular tissue

Testicular tissue was thawed at 4°C for ∼30 min, washed with PS, drained with filter paper and weighed accurately. Then, PS (1:4, *w/v*) was added, and the tissue homogenized. The homogenate (200 μL) was added to 60 μL of SB, and 60 μL of IS (arctigenin) working solution. The mixture was vortex-mixed for 30 s, followed by addition of 400 μL of acetonitrile, vortex-mixing for 120 s, and centrifugation at 13,000 rpm for 15 min at 4°C. The resultant supernatant was injected into the UPLC-MS/MS system. SB concentrations for calibration curves were 0.10, 0.20, 0.50, 1.0, 5.0, 20.0 and 30.0 ng/mL in testicular tissue, whereas those for QC analyses were 1.0, 10.0 and 20.0 ng/mL in testicular tissue. In these samples, the IS concentration was 25.0 ng/mL.

#### Analytical conditions

After oral administration of SB in male mice, concentrations of SB in plasma and testicular tissues were detected by UPLC-MS/MS. Analyses were undertaken on a UPLC system (Acquity UPLC Ӏ-Class system; Waters) consisting of an auto-sampler, quaternary pump, and a column oven. A C18 reverse-phase column (Acquity UPLC BEH, 100 × 2.1 mm, 1.7 μm, 130 Å) was used to separate samples. The mobile phase comprised 0.1% formic acid in water (A) and acetonitrile (B). Gradient elutions were: 0 min 50% B, 0.5 min 50% B, 1.5 min 80% B, and 6 min 50% B. Samples were kept in the autosampler at 4°C until measurement. The column was maintained at 40°C, the flow rate was 0.3 mL/min, and injection volume was 2 μL. Detection was carried out on a Xevo triple quadrupole mass spectrometer (Waters). High-purity nitrogen served as the nebulizing gas and drying gas. Optimal MS conditions were: positive ion mode, source temperature = 110°C, desolvation-gas temperature = 450°C, cone gas flow = 50 h, desolvation gas flow = 600 L/h, capillary voltage = 3.0 kV, sampling cone voltage = 25 V, and extraction cone voltage = 3.0 V. Multiple-reaction monitoring data were acquired in centroid mode between *m* 50 and *m/z* 1000 using MassLynx v4.1 (Waters), and the scan time and interscan time were set at 0.4 s and 0.1 s, respectively. Leucine-enkephalin (*m/z* 556.2771) was used as the external reference of LockSpray infused at a constant flow of 5 μL/min. The mass spectrometer was calibrated over a range of 50–1000 Da with sodium formate. The following precursors to product ions were monitored: *m/z* 401.2843→300.3354 for SB (collision energy, 24 eV; dwell time, 25 ms); *m/z* 289.4323→253.3991 for testosterone (14 eV; 25 ms); *m/z* 373.3807→355.3415 for arctigenin (48 eV; 25 ms).

#### Specificity

Blank plasma, blank plasma with addition of working solutions of SB and IS, and plasma samples after oral administration of SB were analyzed by UPLC/MS/MS for exclusion of interference at the peak concentration of SB or IS. Similarly, specificity for measurement of SB in testicular tissue was also validated.

#### LoQ

The LLoQ was determined as the lowest concentration that the instrument could quantify accurately (i.e., the lowest concentration point on the standard curve).

#### Precision and accuracy

The precision and accuracy were validated by measuring QC samples at 1.0, 10.0 and 20.0 ng/mL of SB in plasma (n = 3) or in testicular tissue (n = 3), respectively. During measurements in 3 consecutive days, the intra- and inter-day variations were calculated. Precision was expressed as the relative standard deviation (RSD)% and accuracy was expressed as the relative error (RE)% by comparing the SB concentration measured with the SB concentration added. The criterion for acceptability was: precision, <15%, accuracy, 85%–115%; LLoQ ±20% accuracy.

#### Extraction recovery

SB recovery from plasma or testicular tissue was calculated by comparing the SB concentration measured with the SB concentration added.

#### Sample stability

SB stability was assessed on the QC samples mentioned above at three concentrations after three freeze–thaw cycles (−20°C to 25°C) on 3 consecutive days, storage at 25°C for 24 h, and storage at −80°C for 1 month, respectively. Sample stability was expressed as the RSD for the SB concentration measured.

#### Dosing

Normal male mice (n = 65) were fasted 12 h but had free access to water. Then, each mouse was administered (p.o.) a single dose of SB (20 mg/kg, i.g.) for subsequent experiments.

#### Sampling

Blood sampling was done at 0 min (before dosing), 15 min, 30 min, as well as 1, 1.5, 2, 3, 4, 5, 6, 8, 12 and 24 h (five mice at each time point) under anesthesia. Aliquots of venous blood (0.75) were sampled, centrifuged at 5,000 rpm for 10 min at 4°C to obtain plasma, and stored at −80°C until use. After each blood sampling, animals were sacrificed, testicular tissues were collected on an ice plate, and frozen immediately at −80°C for use.

#### Plasma handling

Plasma was thawed at 4°C for ∼30 min and vortex-mixed for 30 s. Plasma (200 μL) was vortex-mixed with 60 μL of IS solution (testosterone) for 30 s, followed by addition of 60 μL of methanol. After vortex-mixing for 30 s, 600 μL of acetonitrile was added, followed by vortex-mixing for 120 s, and centrifugation at 13,000 rpm for 10 min at 4°C. The resultant supernatant was injected into the UPLC-MS/MS system.

#### Handling of testicular tissue

Testicular tissue was thawed at 4°C for ∼30 min, washed with PS, drained with filter paper, and weighed accurately. Then, PS (1:4, *w/v*) was added and the tissue homogenized. The homogenate (200 μL) was added to 60 μL of methanol and 60 μL of IS (arctigenin) working solution. The mixture was vortex-mixed for 30 s, followed by addition of 400 μL of acetonitrile, vortex-mixing for 120 s, and centrifugation at 13,000 rpm for 15 min at 4°C. The resultant supernatant was injected into the UPLC-MS/MS system.

#### Pharmacokinetic analyses

Pharmacokinetic parameters in plasma and testicular tissue were calculated using a non-compartmental approach employing DAS v3.2 (China State Drug Administration, Shanghai, China).

### Statistical analyses

Statistical analyses were conducted by Prism v7.0 (GraphPad, La Jolla, CA, USA) and SPSS v20.0 (IBM). No data were excluded from analyses. The Student’s *t*-test (two-tailed) or one-way analysis of variance was used for statistical analyses. p < 0.05 was considered significant. Data are the mean ± standard deviation.

## Supplementary information

All data needed to understand and assess the conclusions of this research are available in the main text and supplementary materials. Raw datasets supporting the findings of this study are available online or from the corresponding author.

## Acknowledgements

We are grateful to the biological expertise provided by Chong Tang at the BGI Genomics Co., Ltd.

## Author contributions

Lu W.-L., Lin R.-C. and Yang W.-P. designed the study and supervised the analyses.

Zou D.-X., and Meng X.-D. completed the major research work. Xie Y., Liu R., Duan J.-L., Bao C.-J., and Liu Y.-X. undertook experiments under the direction of Lu W.-L., Lin R.-C. and Yang W.-P.

Du Y.-F., Xu J.-R., Luo Q., Zhang Z. and Ma S. helped with data analyses.

Zou D.-X., Meng X.-D., Yang W.-P., Lin R.-C. and Lu W.-L. wrote the manuscript with input from all authors.

All authors approved the final version for submission.

---

**Figure S1.**
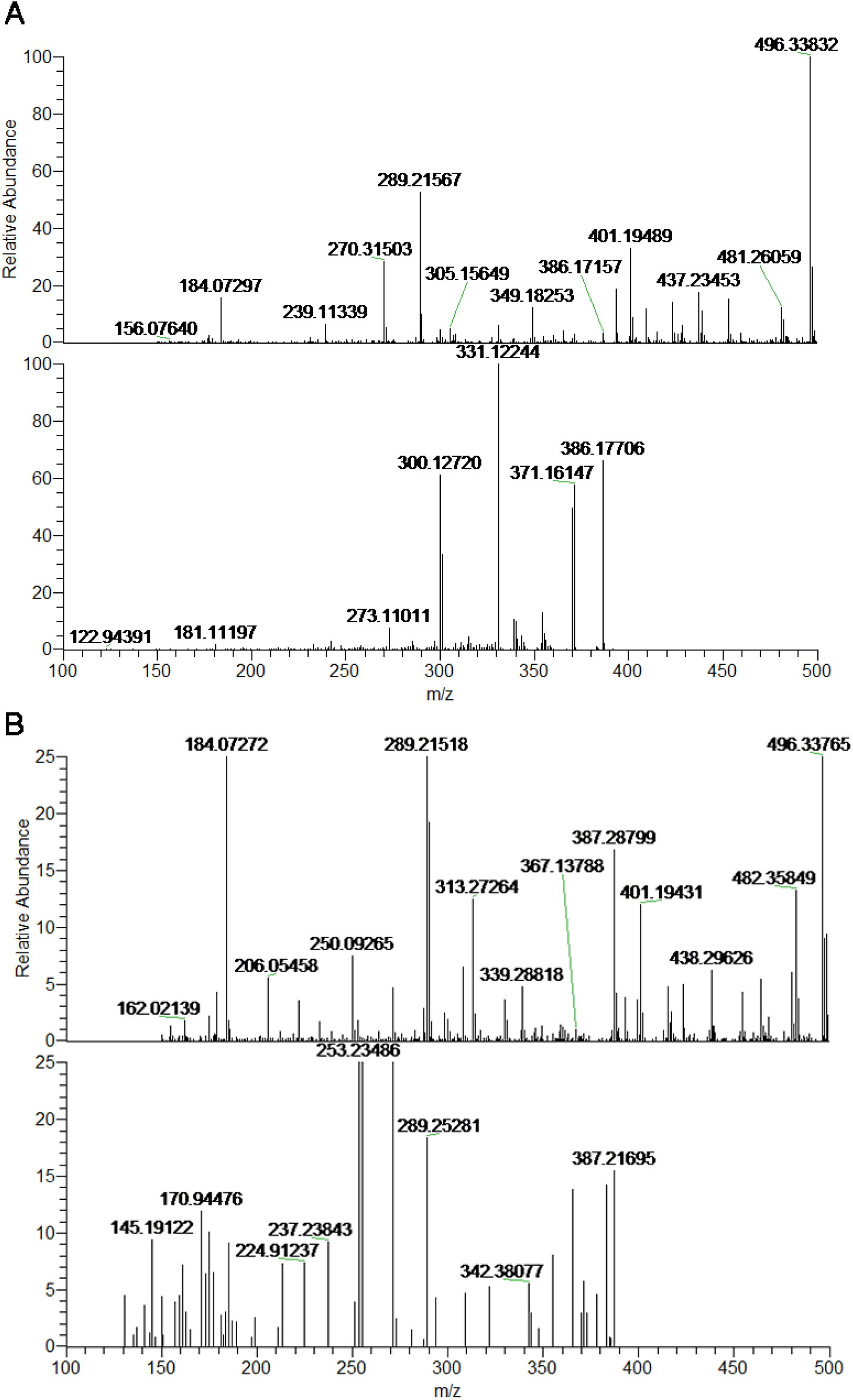
Identification of schisandrin B in mouse plasma and testicular tissue. Notes: **L.** Typical mass spectra of mouse plasma after oral administration of schisandrin B (20mg/kg) at 3 h in **Fig.1G**, which was used as identifying schisandrin B structure in plasma. **M.** Typical mass spectra of mouse testicular tissue after oral administration of schisandrin B (20mg/kg) at 3 h in **Fig.1I**, which was used for identifying schisandrin B structure in testicular tissue.

**Supply Dataset S1.**
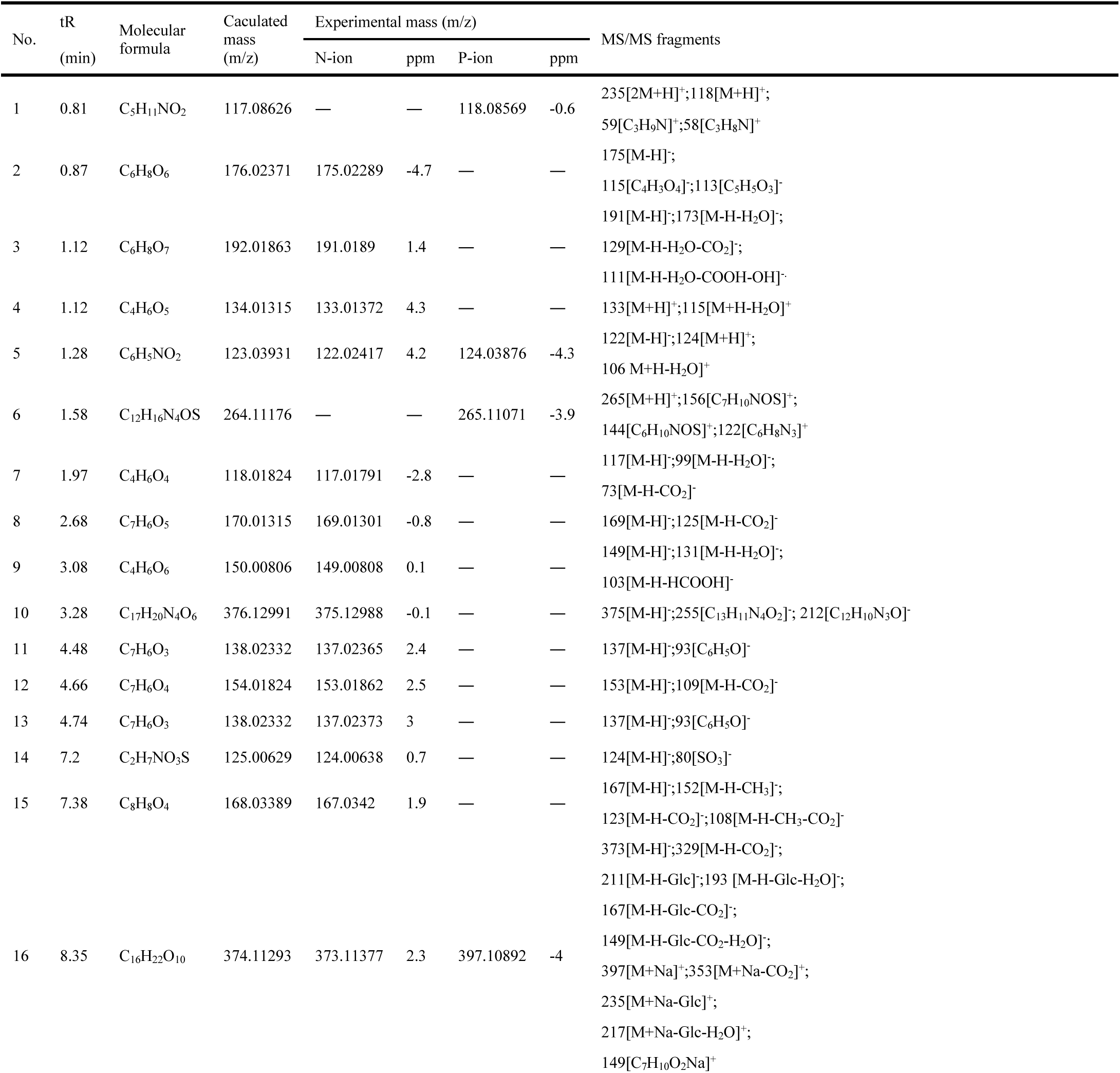
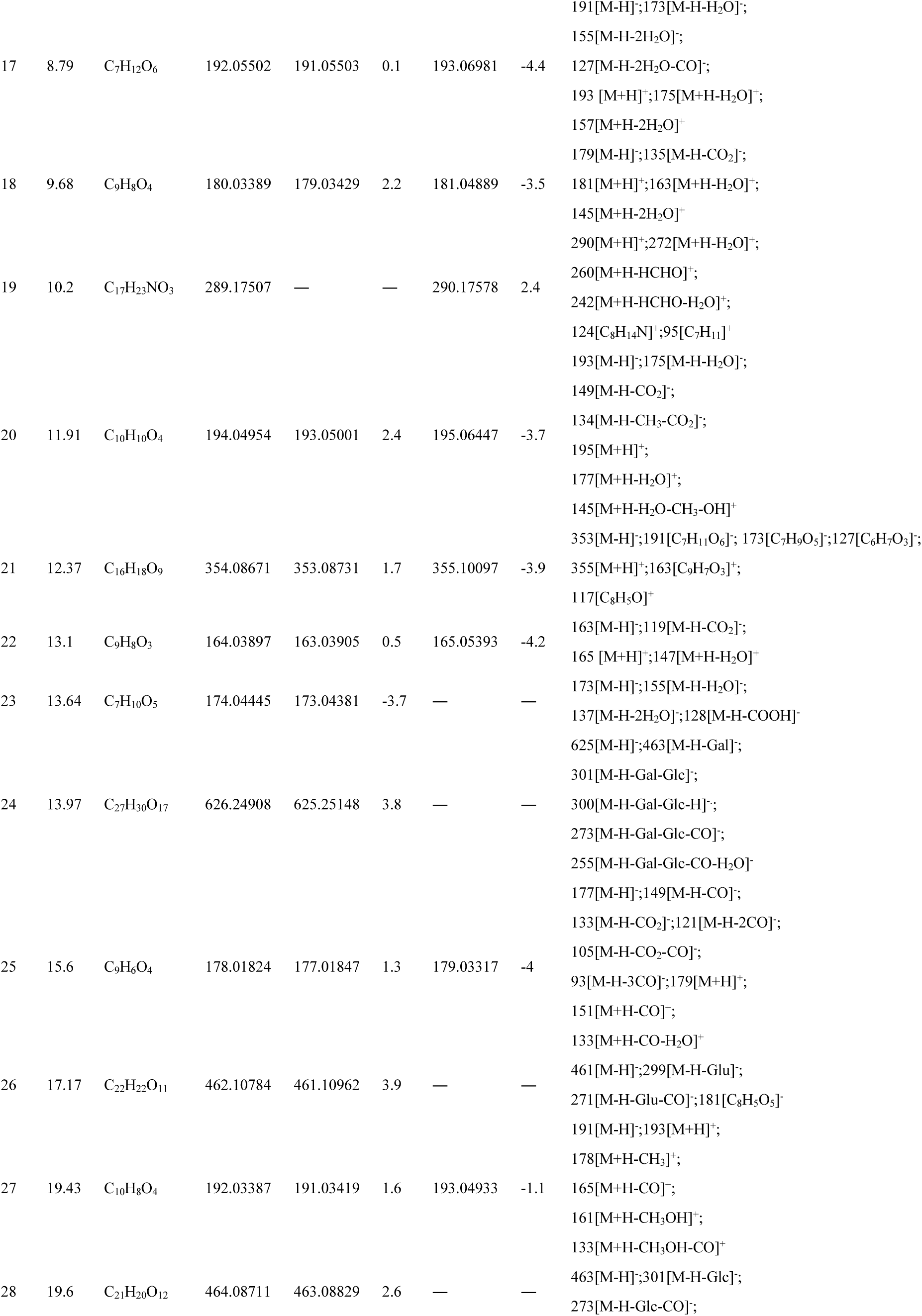
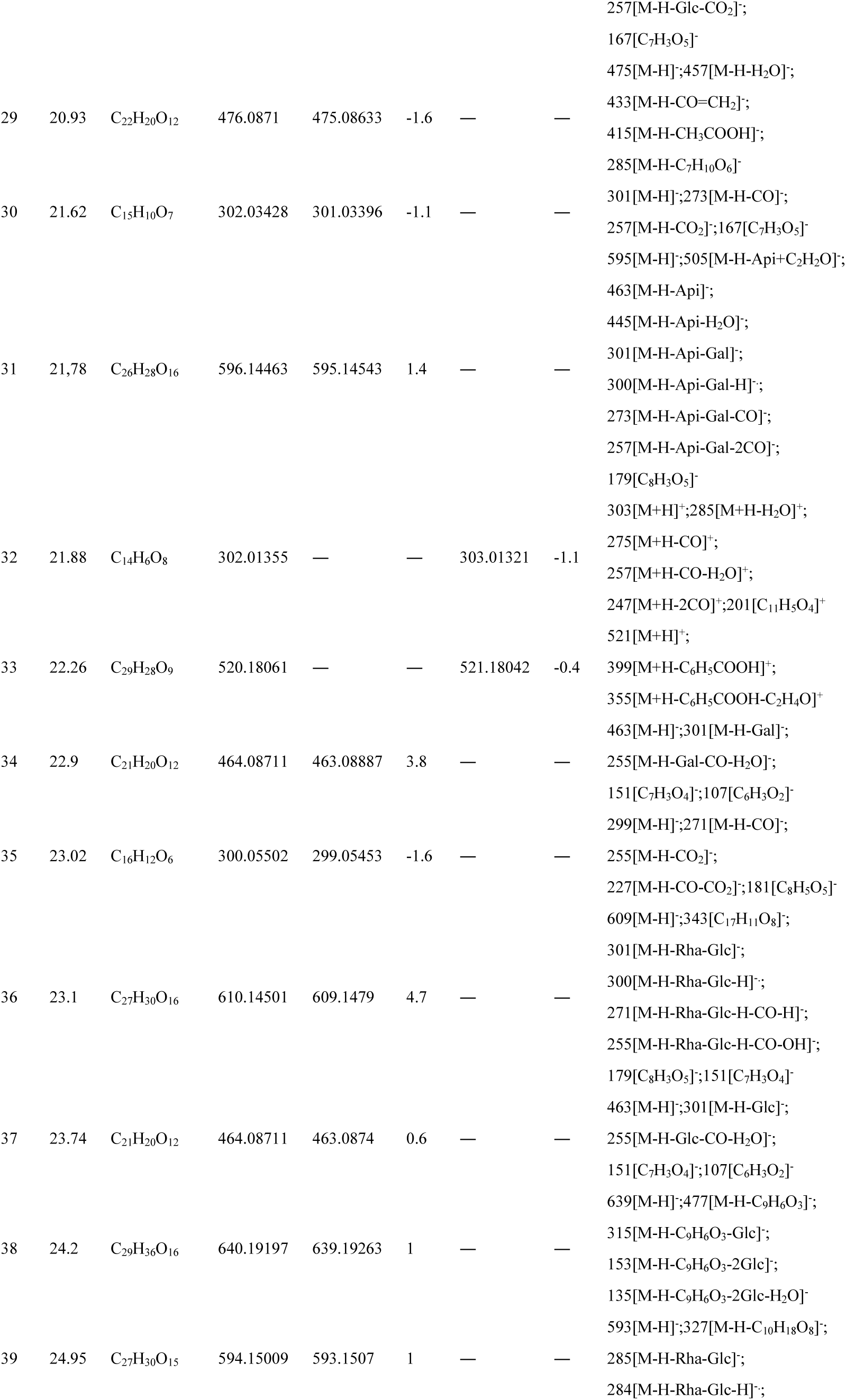
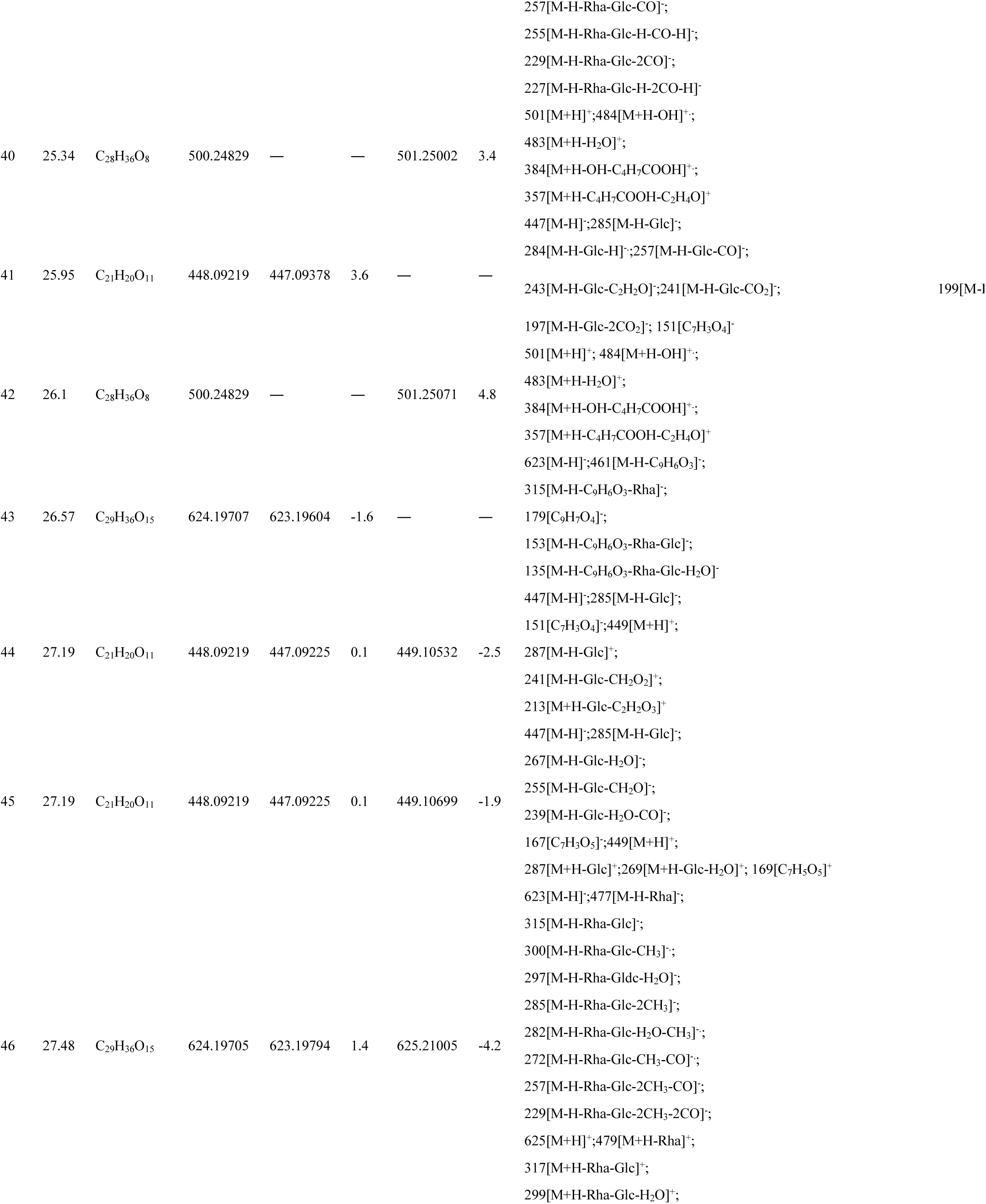
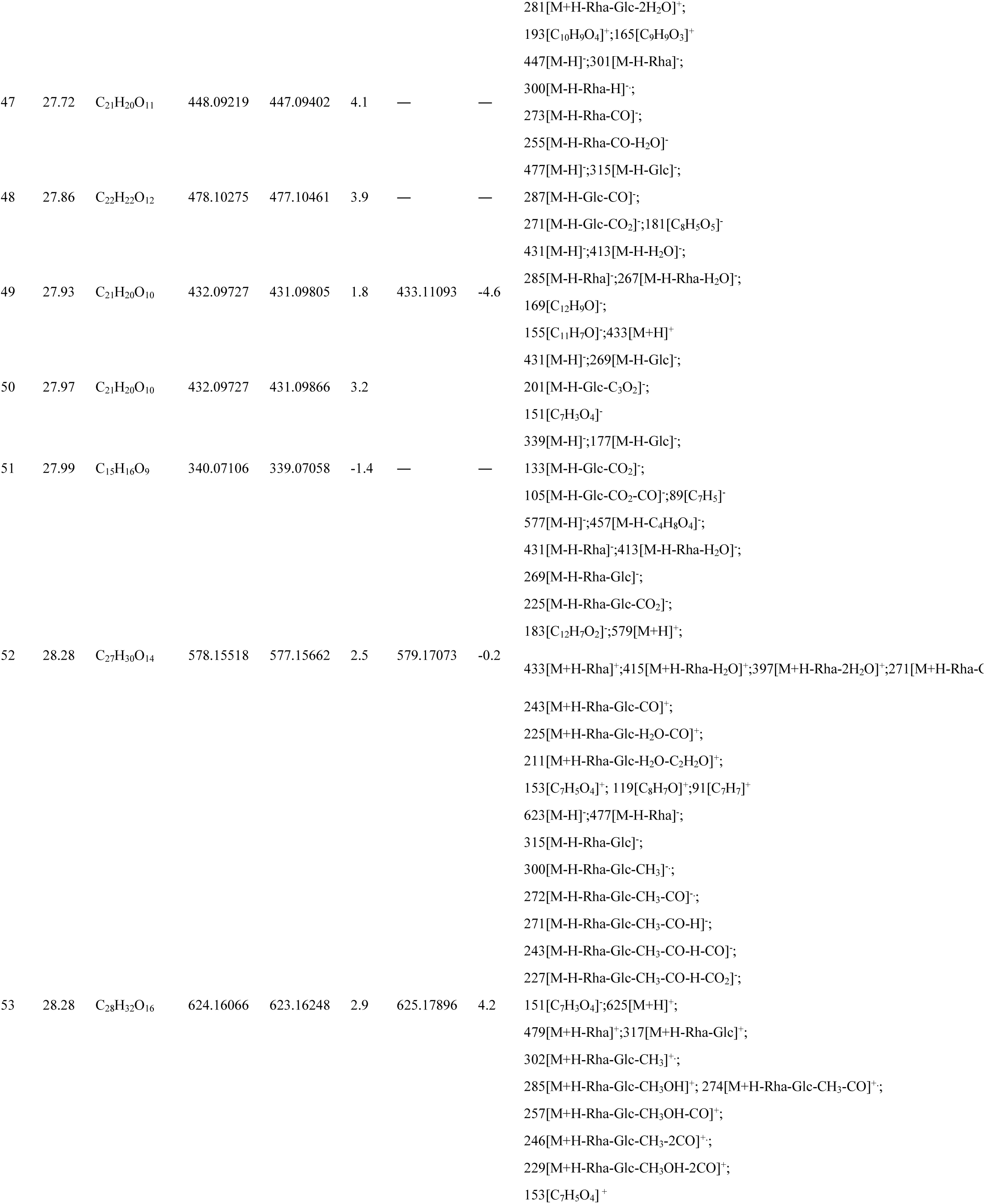
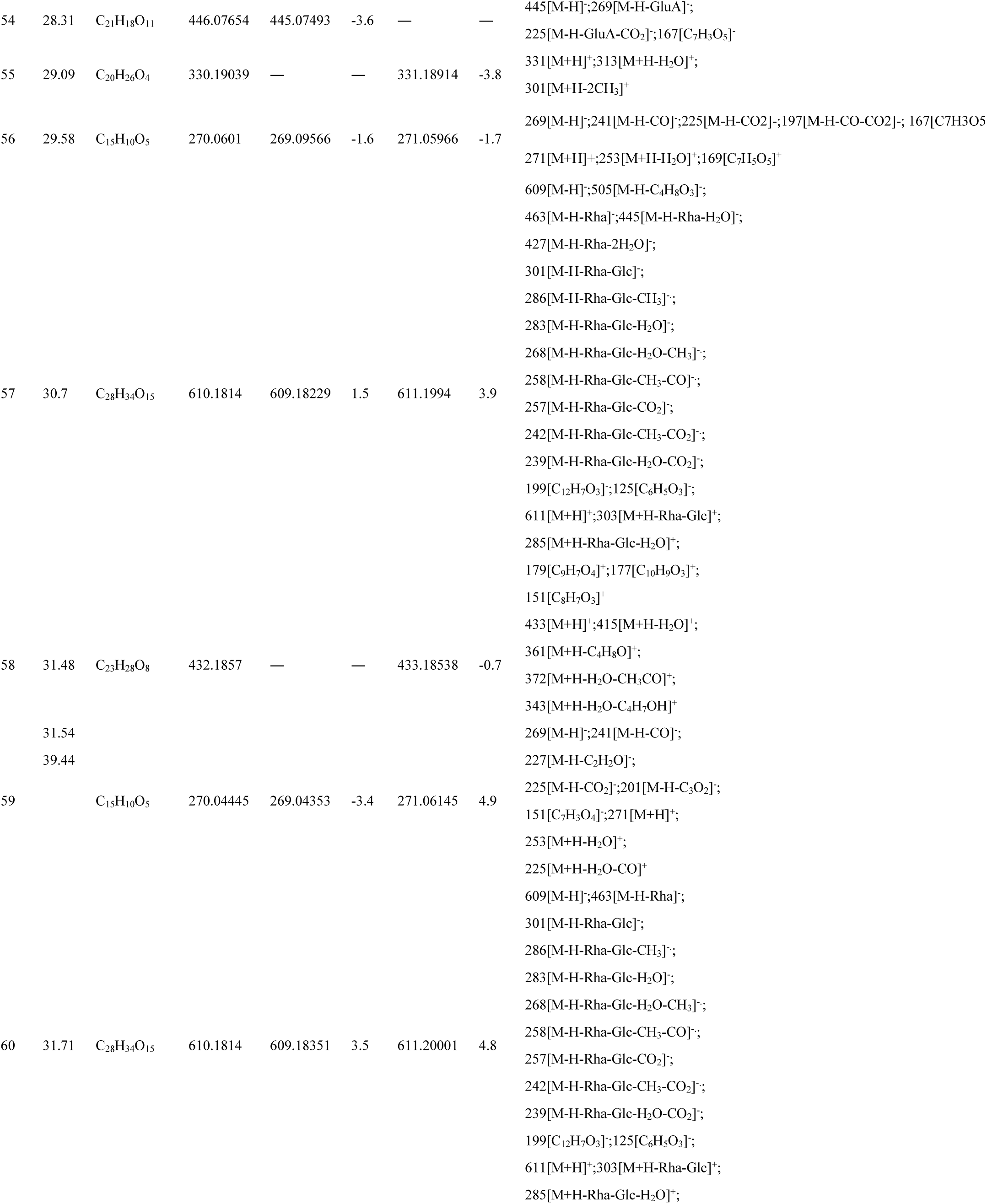
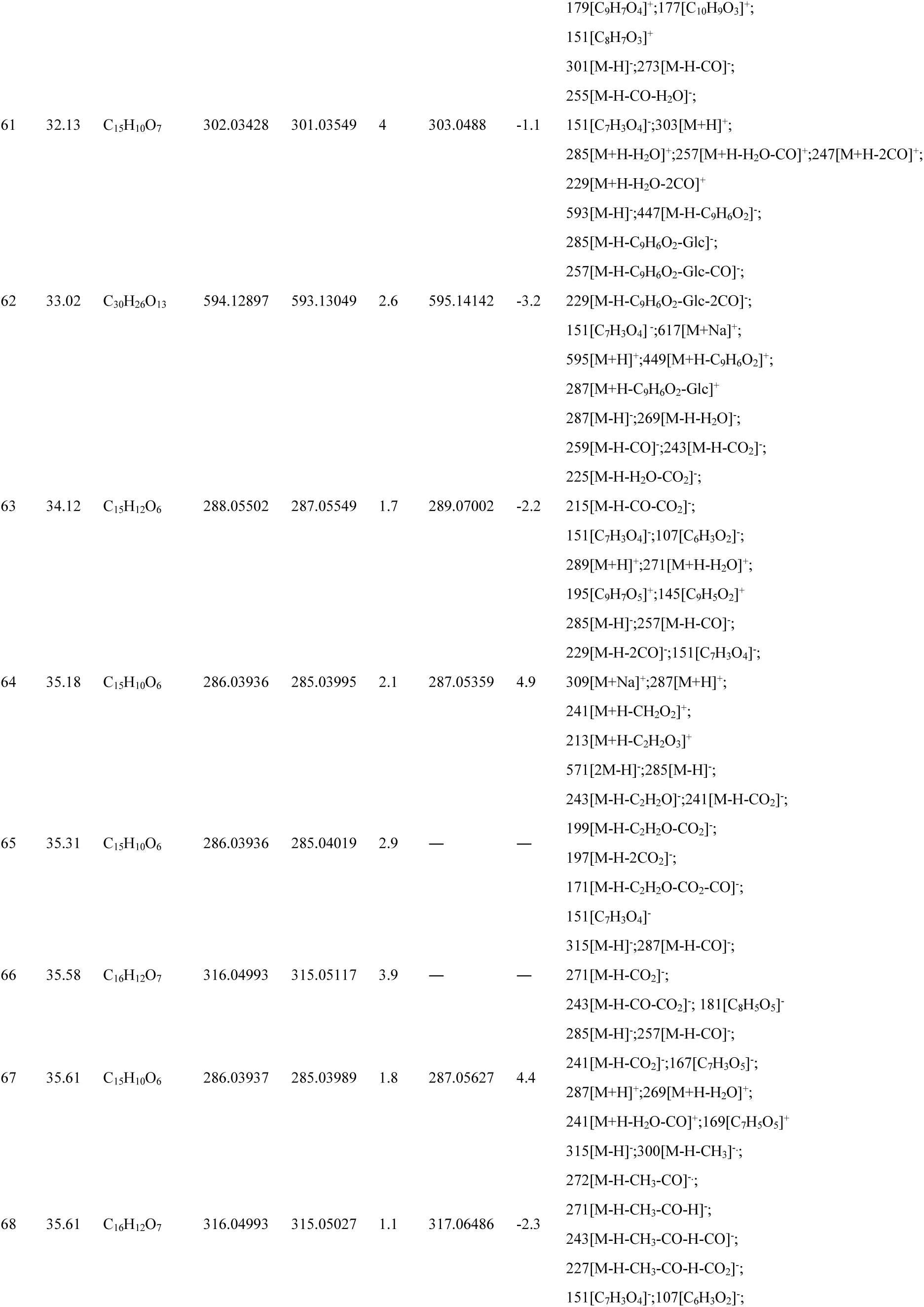
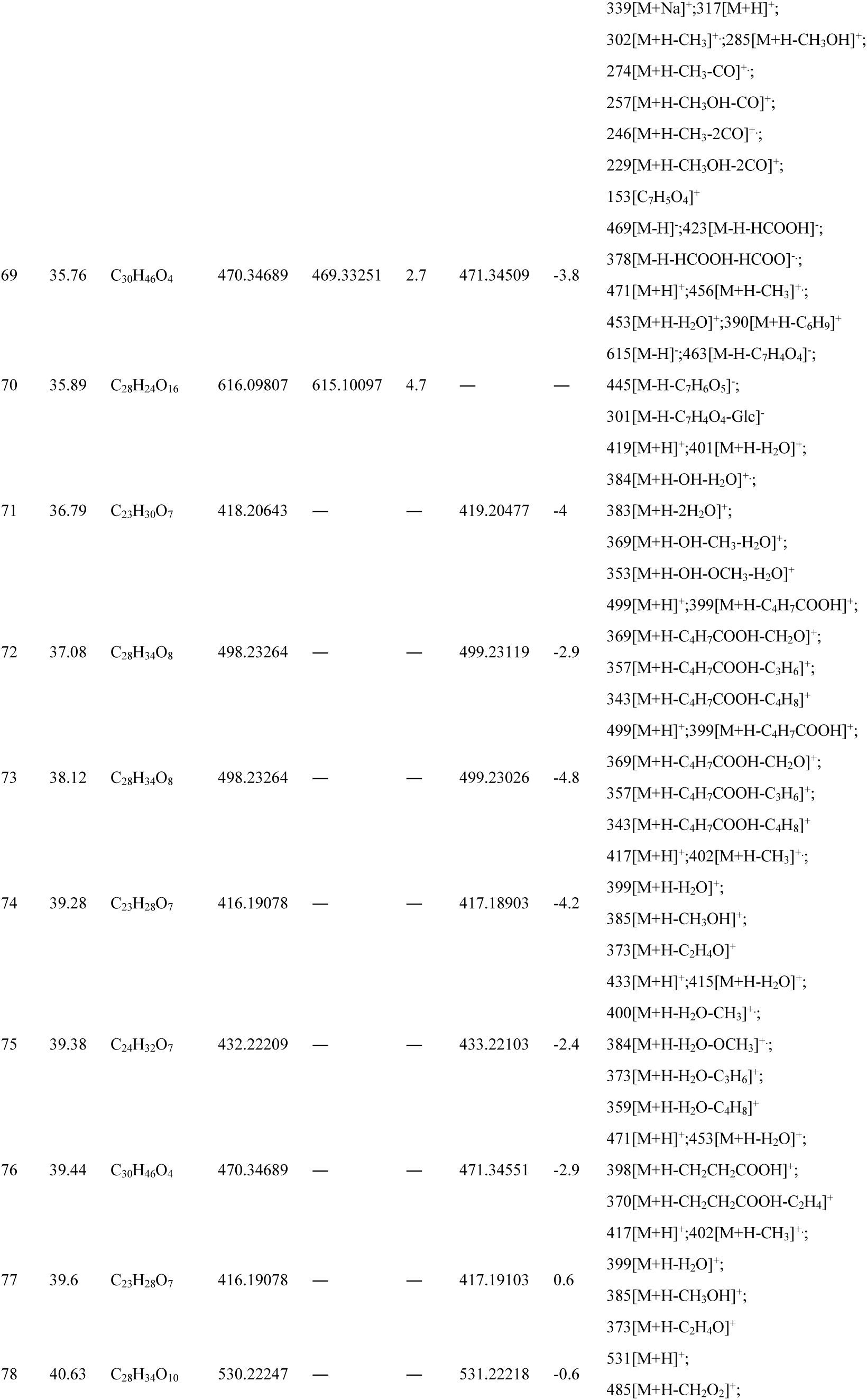
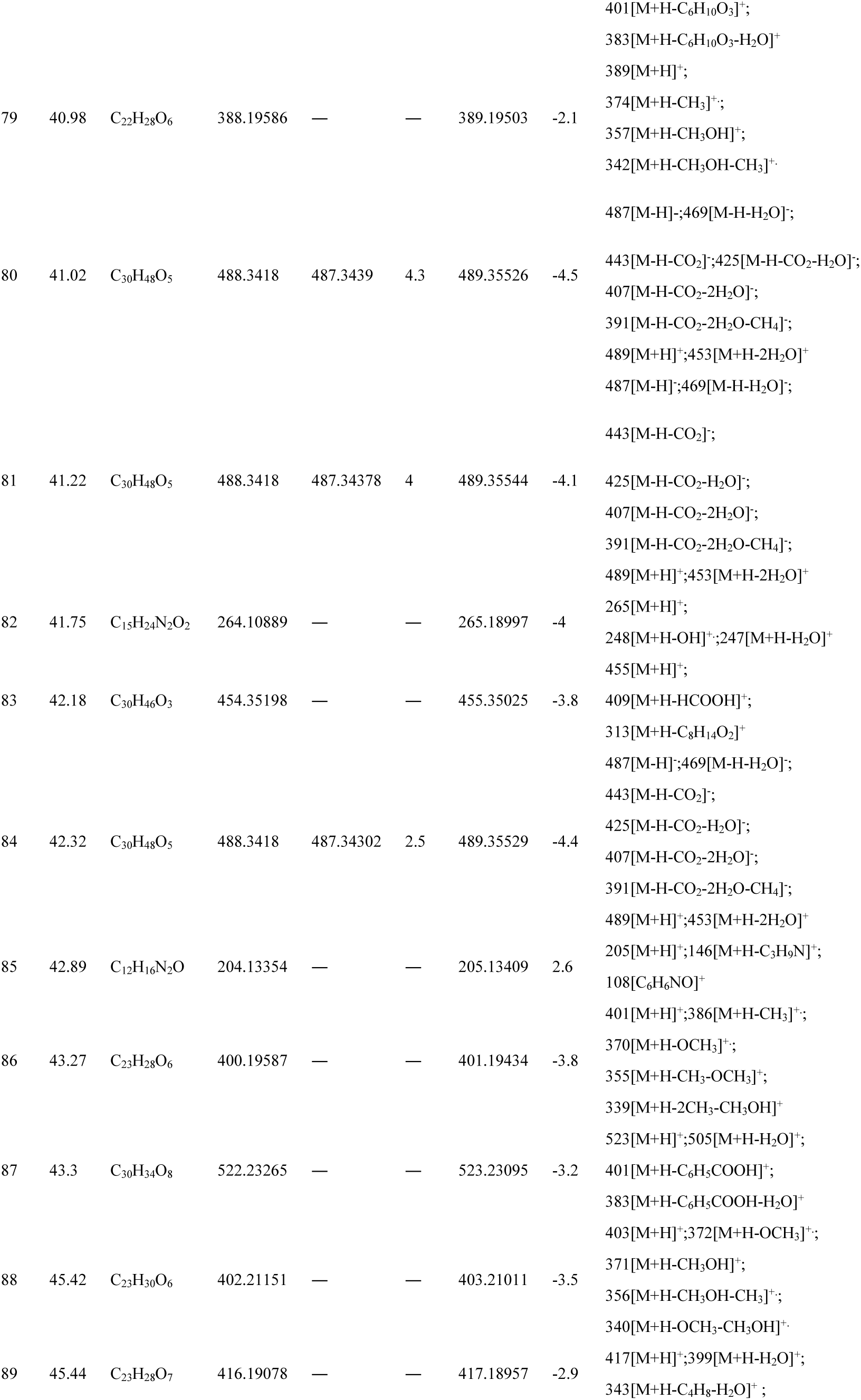
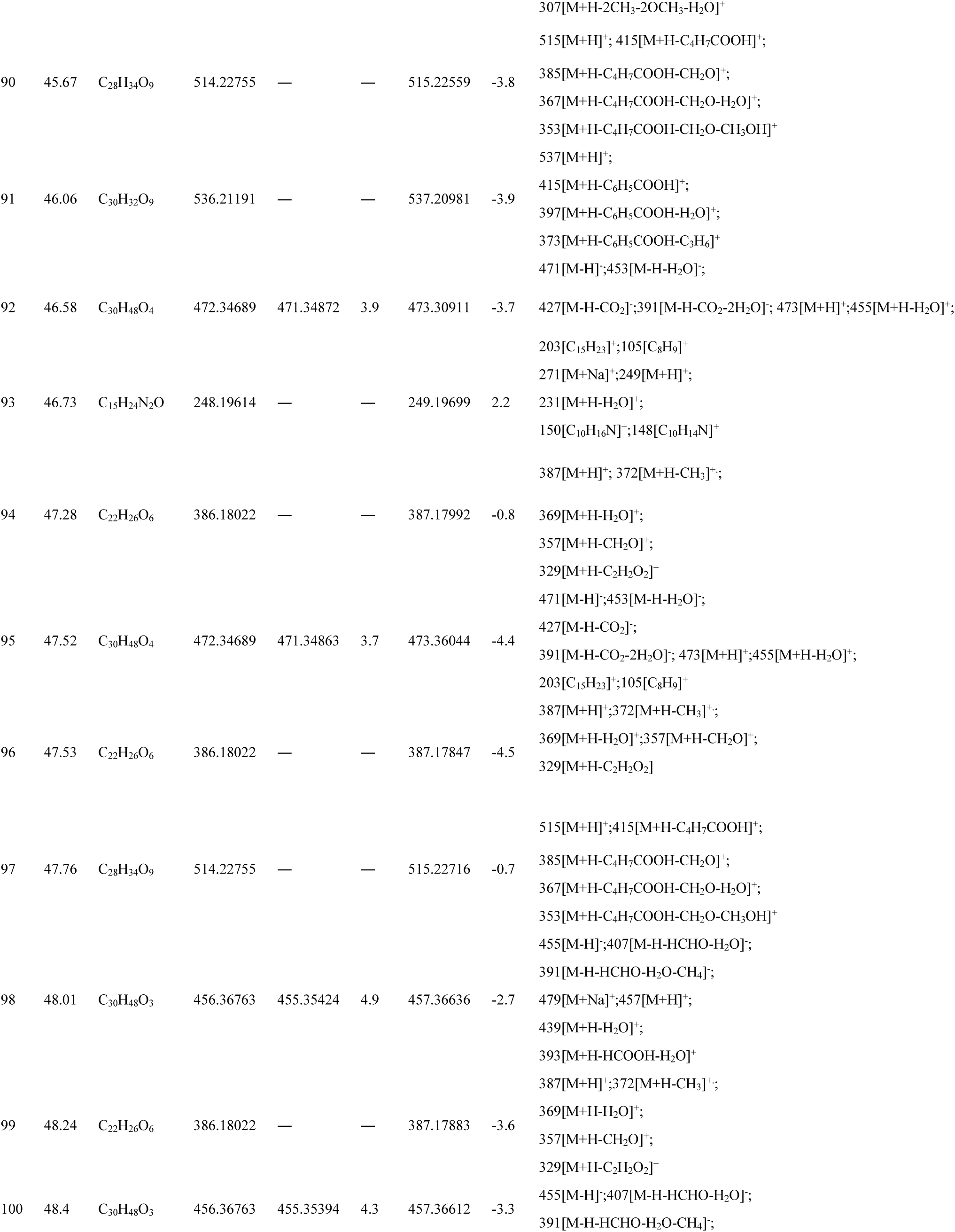
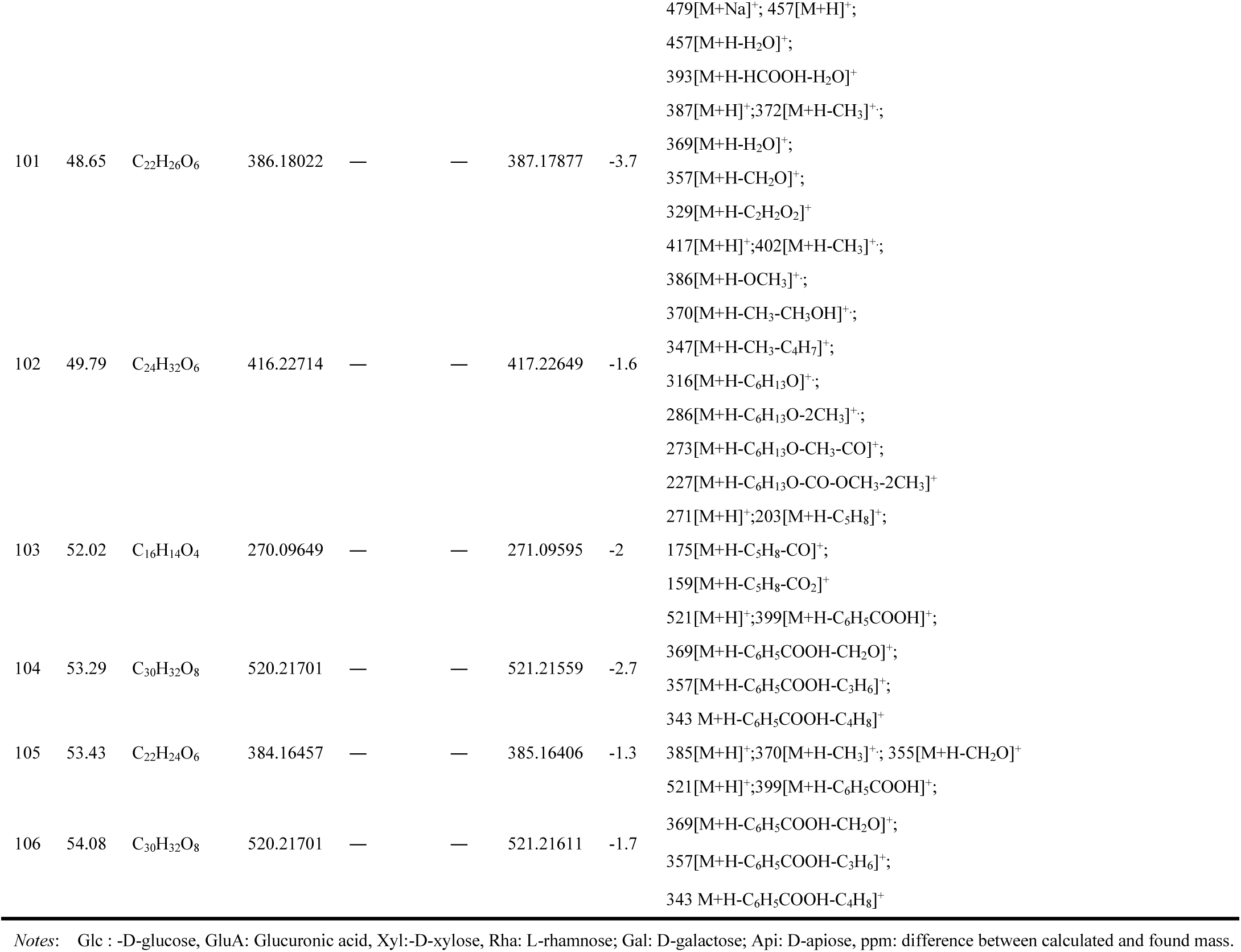

**Supply Dataset S2.**
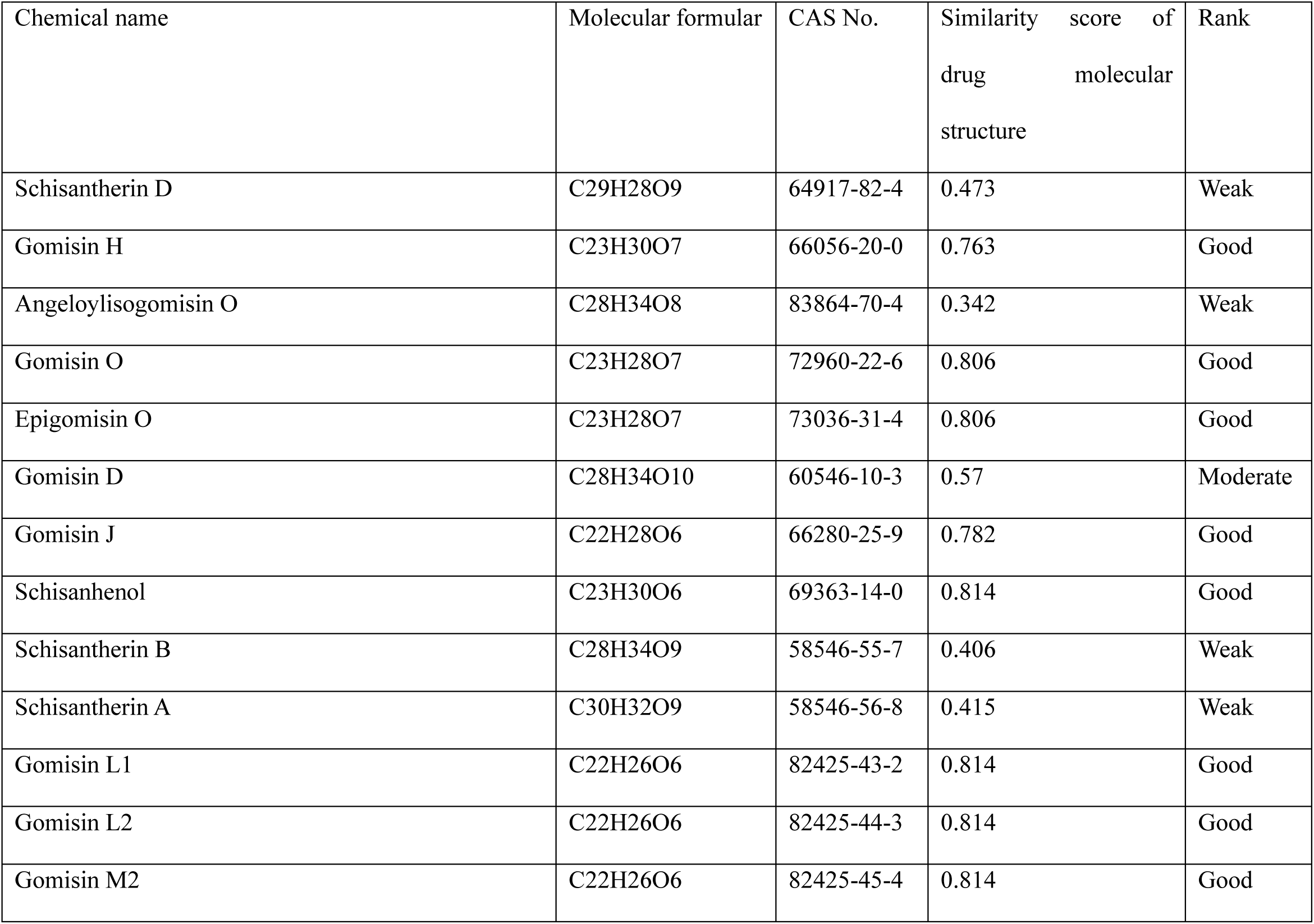
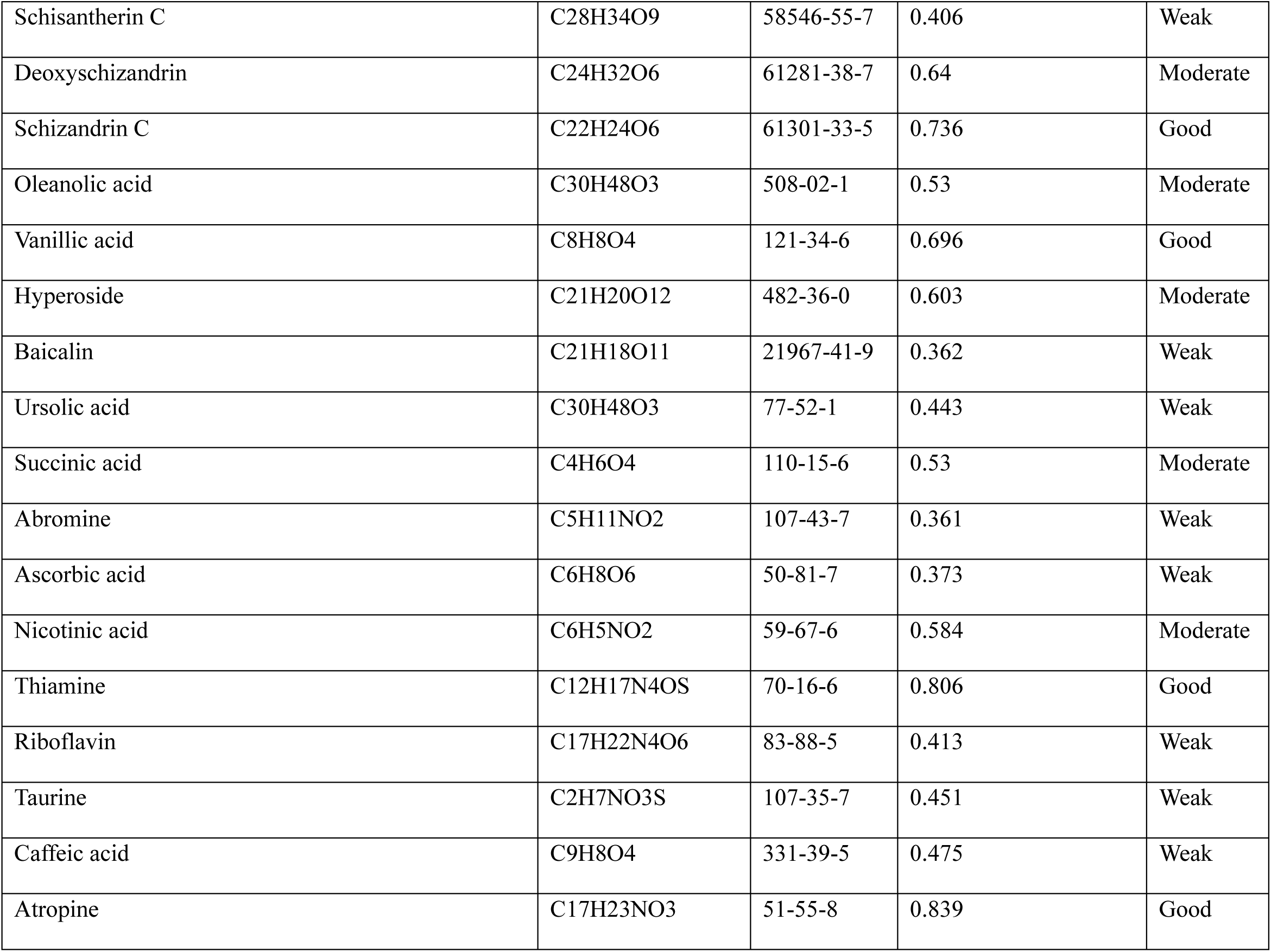
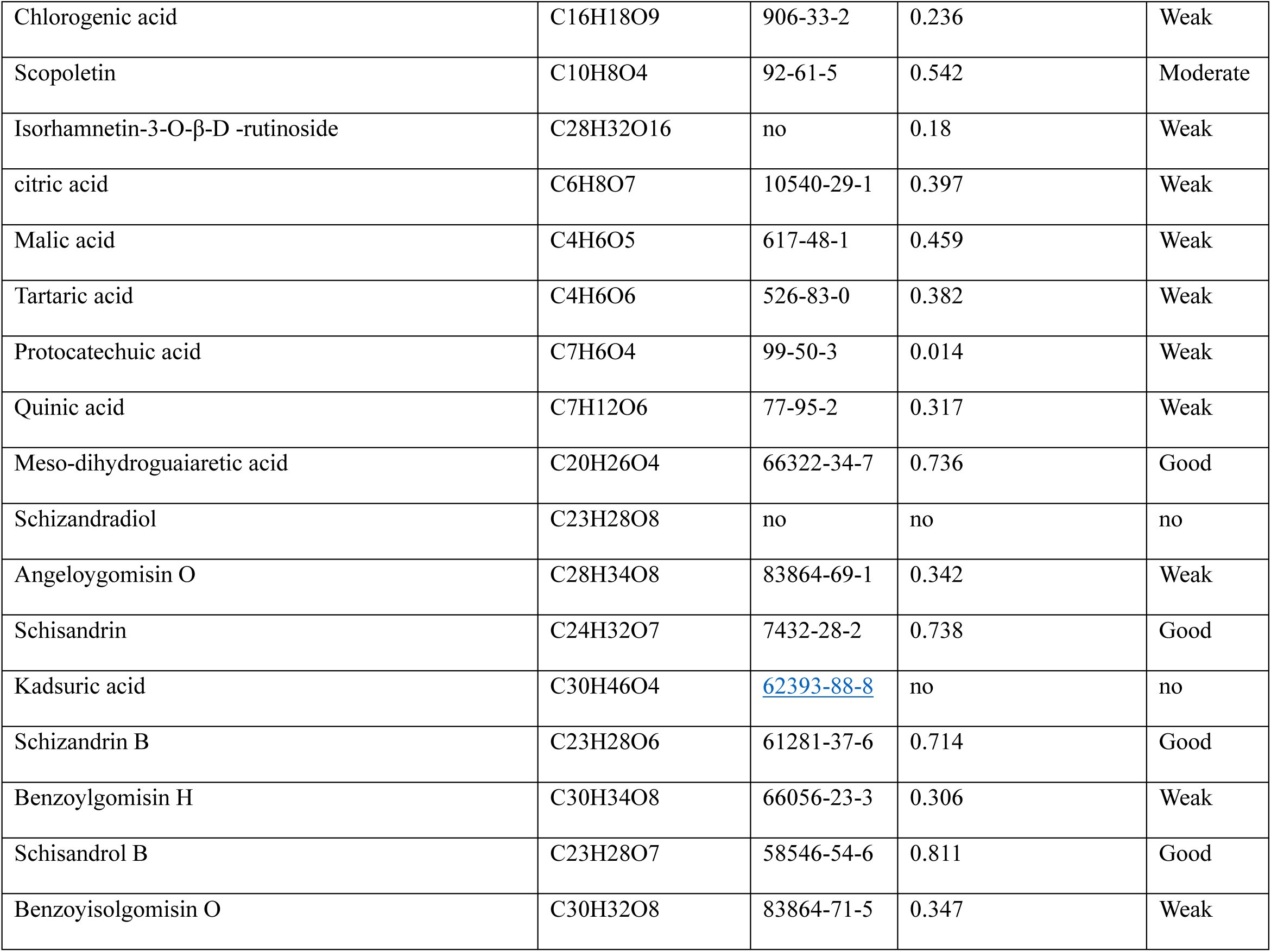
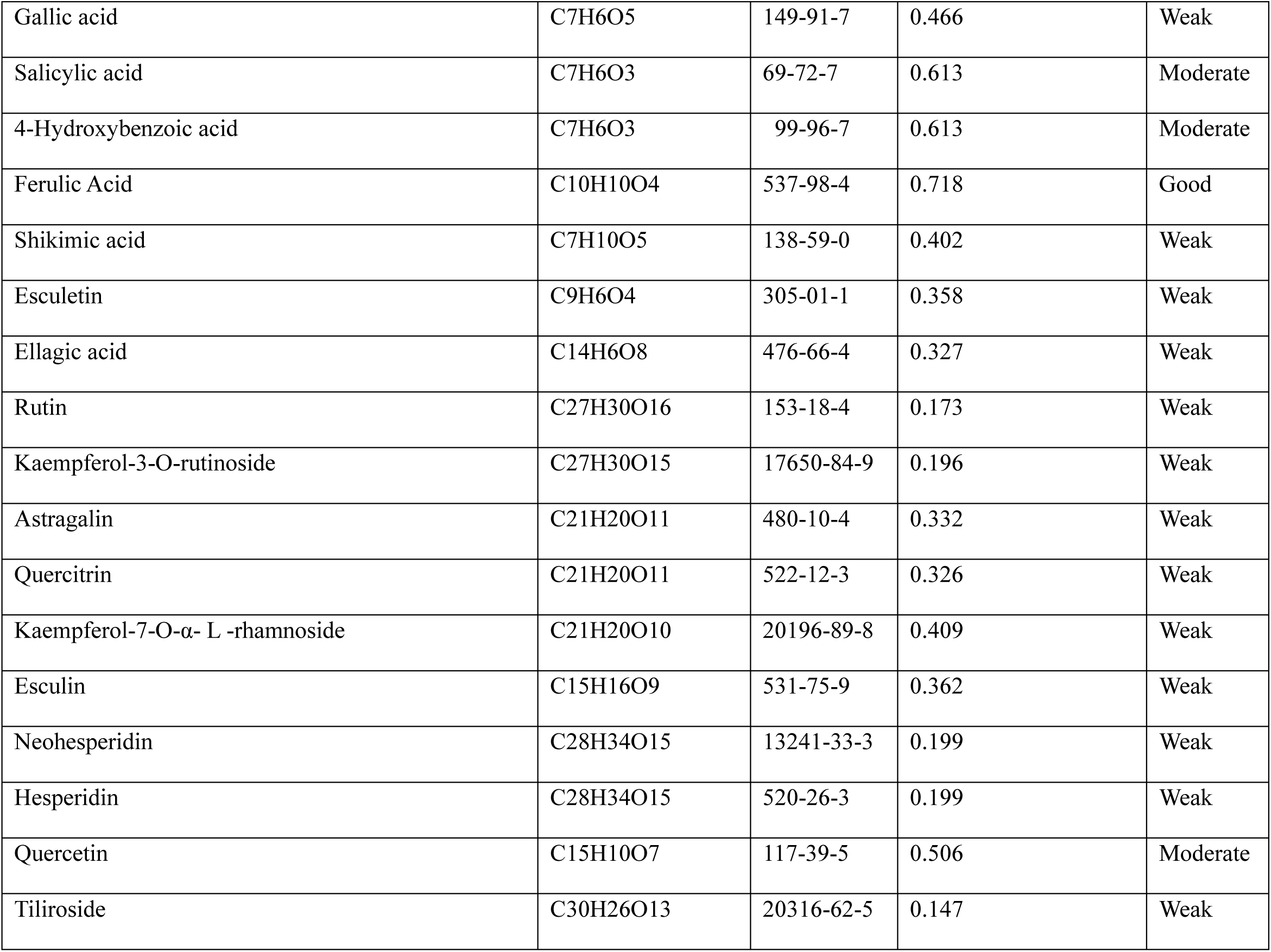
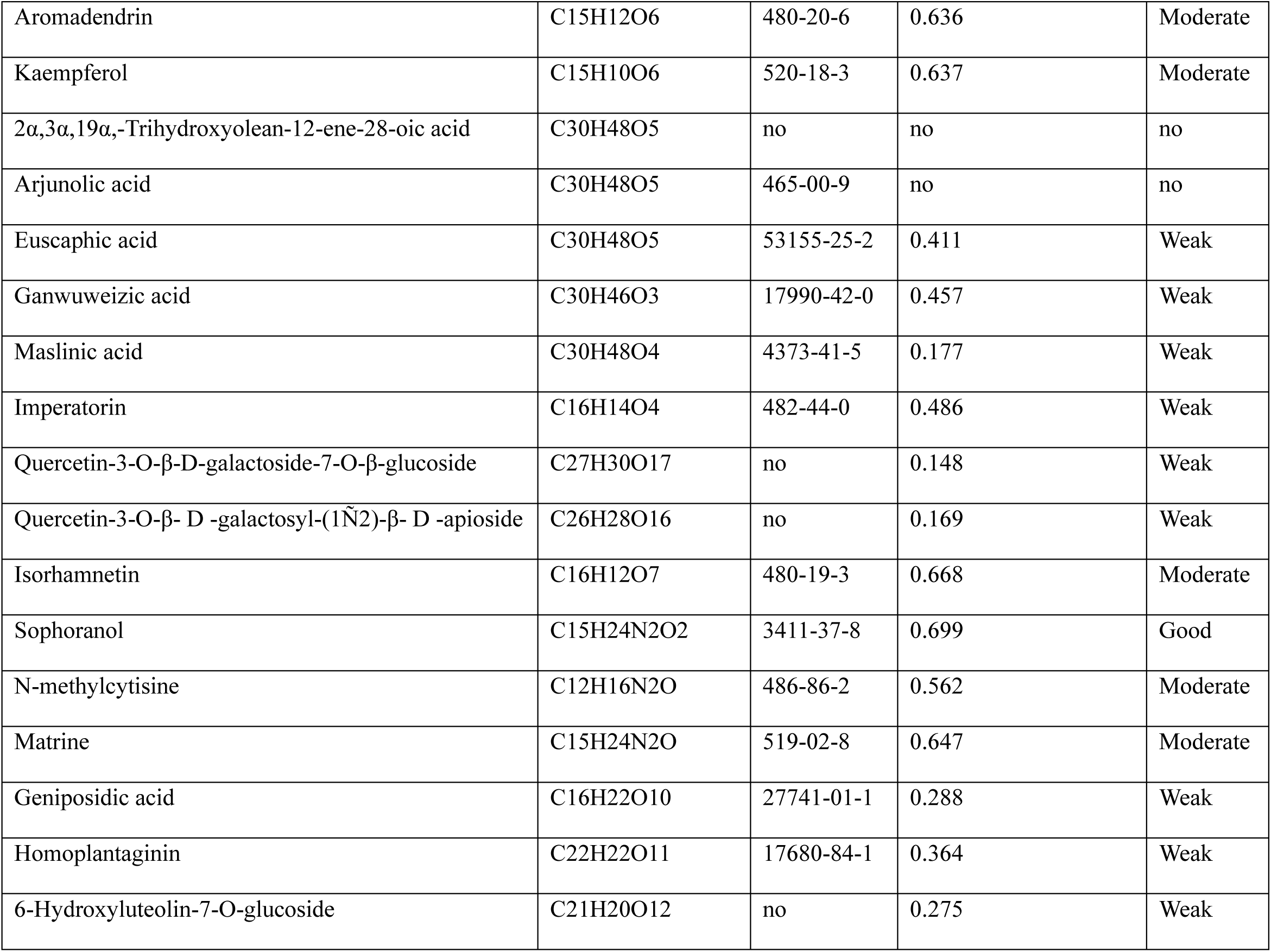
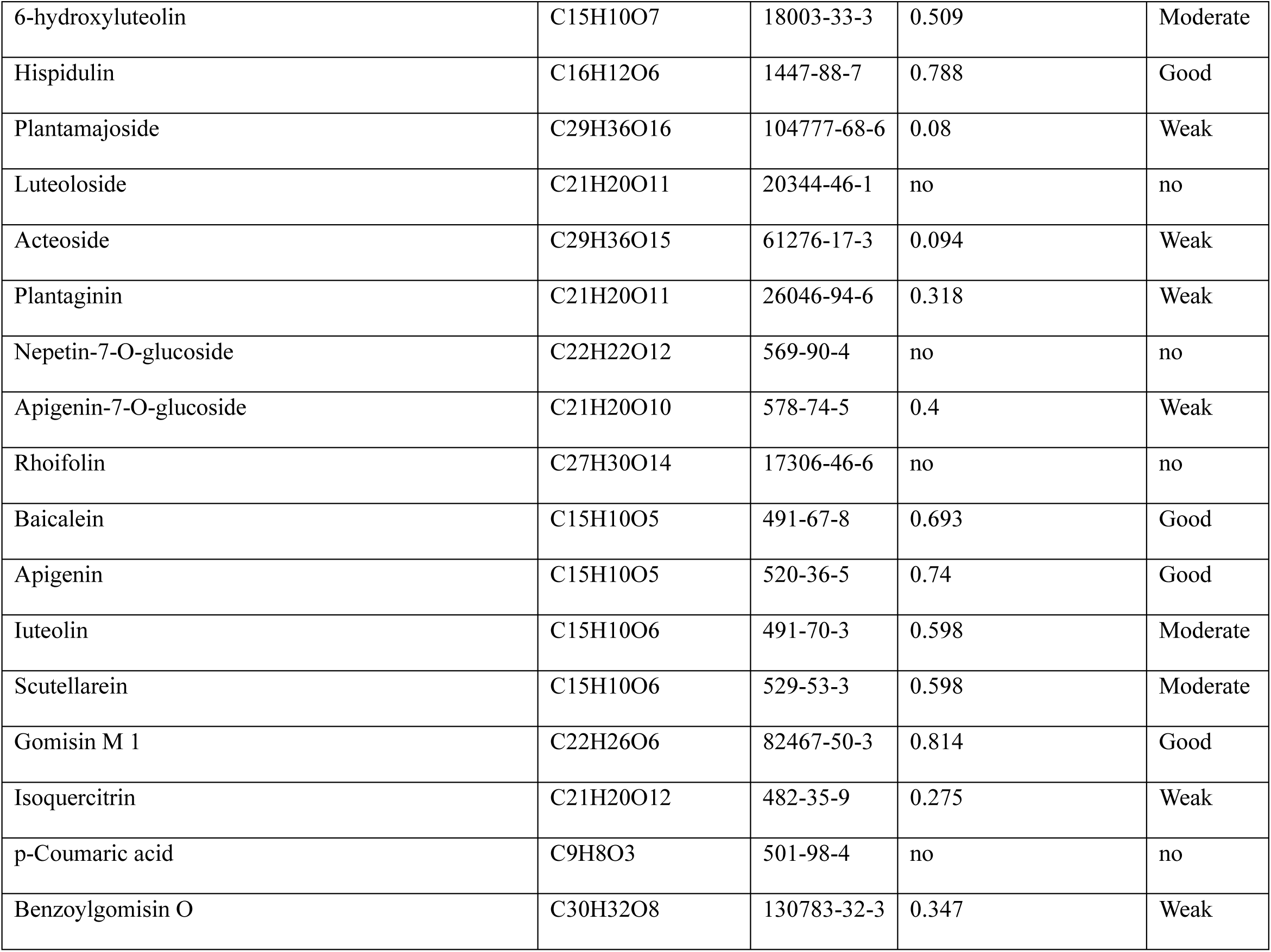
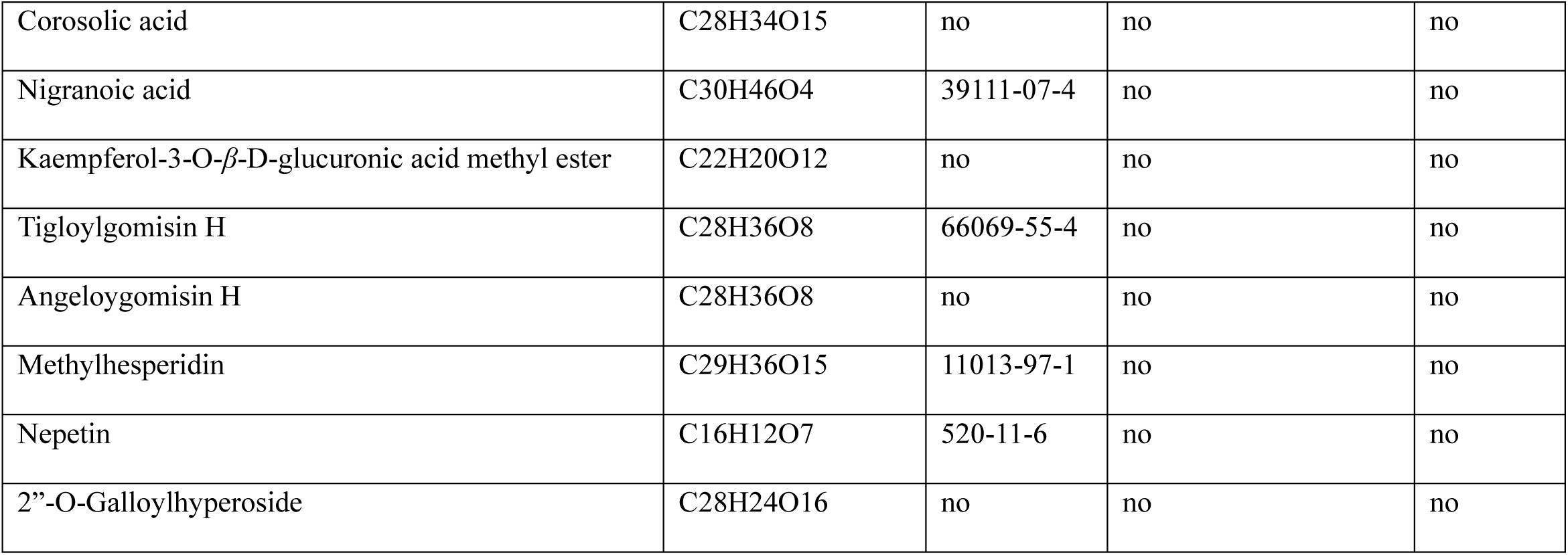

**Supply Dataset S3.**

| Chemical name | Type | Factor<br>1(Similarity<br>score of drug<br>molecular<br>structure) | Factor<br>2<br>(Peak<br>abundance) | FAC1_1 | FAC2_1 | Factor<br>comprehensive<br>score |
| --- | --- | --- | --- | --- | --- | --- |
| Schisandrin B | Phenylpropanoids | 0.714 | 340876 | 3.53928 | -0.85662 | 1.751 |
| Epigomisin O | Phenylpropanoids | 0.806 | 151336 | 1.32027 | 0.88705 | 1.144 |
| Schisandrin | Phenylpropanoids | 0.738 | 183899 | 1.59647 | -0.52995 | 0.731 |
| Schisandrol B | Phenylpropanoids | 0.811 | 82017 | 0.45109 | 0.91145 | 0.638 |
| Schisanhenol | Phenylpropanoids | 0.814 | 76773 | 0.39061 | 0.96943 | 0.626 |
| Atropine | Alkaloids | 0.839 | 13608 | -0.36027 | 1.42935 | 0.368 |
| Thiamine | Alkaloids | 0.806 | 41147 | -0.07737 | 0.75568 | 0.262 |
| Gomisin J | Phenylpropanoids | 0.782 | 53591 | 0.03218 | 0.2567 | 0.124 |
| Hispidulin | Flavonoids | 0.788 | 44121 | -0.07587 | 0.37386 | 0.107 |
| Gomisin M2 | Phenylpropanoids | 0.814 | 9399 | -0.46396 | 0.8891 | 0.087 |
| Gomisin M1 | Phenylpropanoids | 0.814 | 8546 | -0.47478 | 0.88809 | 0.08 |
| Gomisin L1 | Phenylpropanoids | 0.814 | 6791 | -0.49704 | 0.886 | 0.066 |
| GomisinL2 | Phenylpropanoids | 0.814 | 6283 | -0.50349 | 0.88539 | 0.062 |
| Gomisin O | Phenylpropanoids | 0.806 | 10447 | -0.46677 | 0.71908 | 0.016 |
| Gomisin H | Phenylpropanoids | 0.763 | 12396 | -0.52858 | -0.19919 | -0.395 |
| Apigenin | Flavonoids | 0.74 | 10053 | -0.60458 | -0.69439 | -0.641 |
| Meso-dihydroguaiaretic acid | Phenylpropanoids | 0.736 | 11459 | -0.59479 | -0.77835 | -0.669 |
| Schizandrin C | Phenylpropanoids | 0.736 | 10777 | -0.60344 | -0.77916 | -0.675 |
| Ferulic Acid | Organic Acids | 0.718 | 13077 | -0.61049 | -1.16178 | -0.835 |
| Vanillic acid | Organic Acids | 0.696 | 37631 | -0.34332 | -1.60351 | -0.856 |
| Sophoranol | Alkaloids | 0.699 | 30676 | -0.4255 | -1.54758 | -0.882 |
| Baicalein | Flavonoids | 0.693 | 10015 | -0.69964 | -1.70066 | -1.107 |

**Supply Dataset S4.**
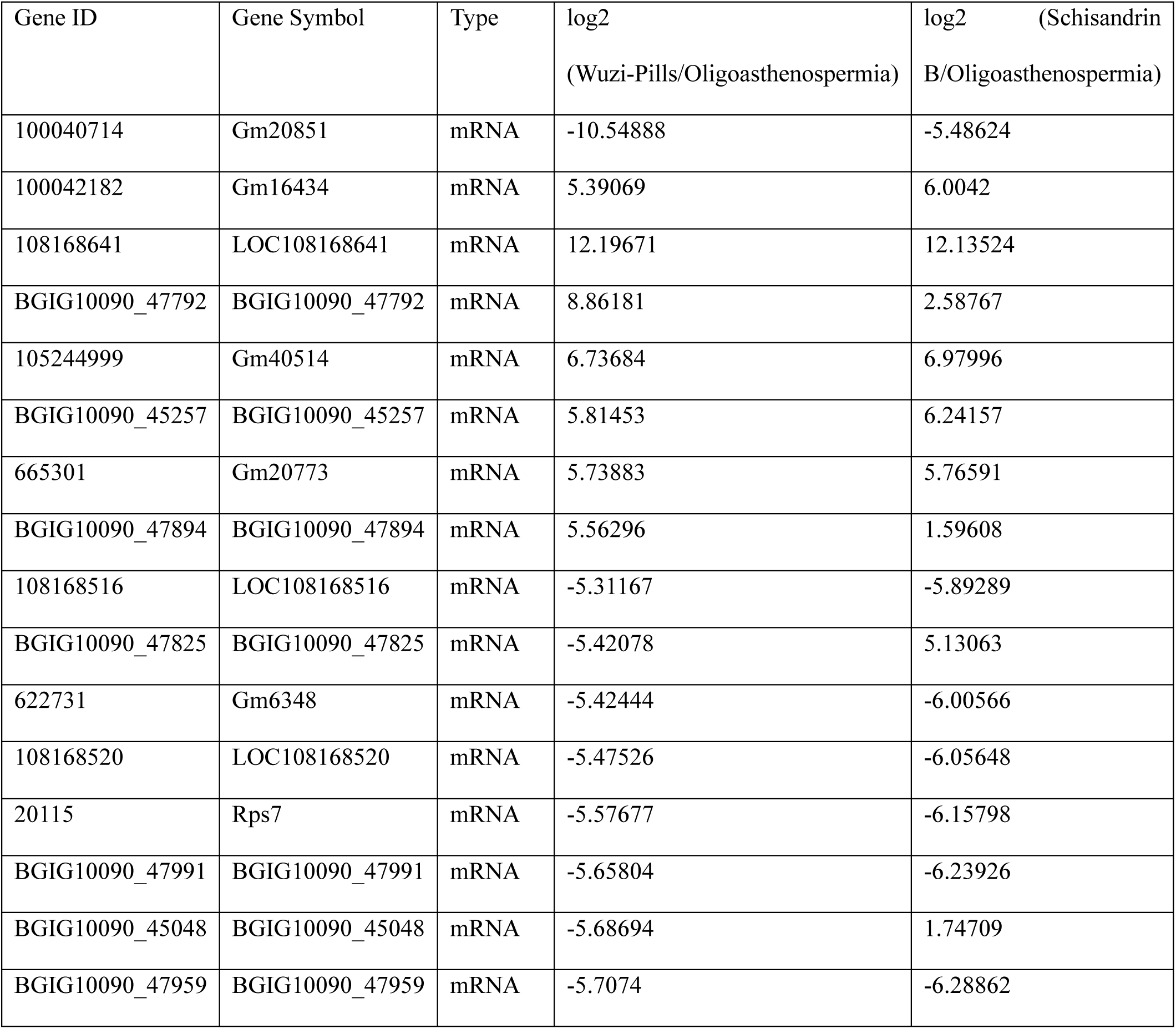
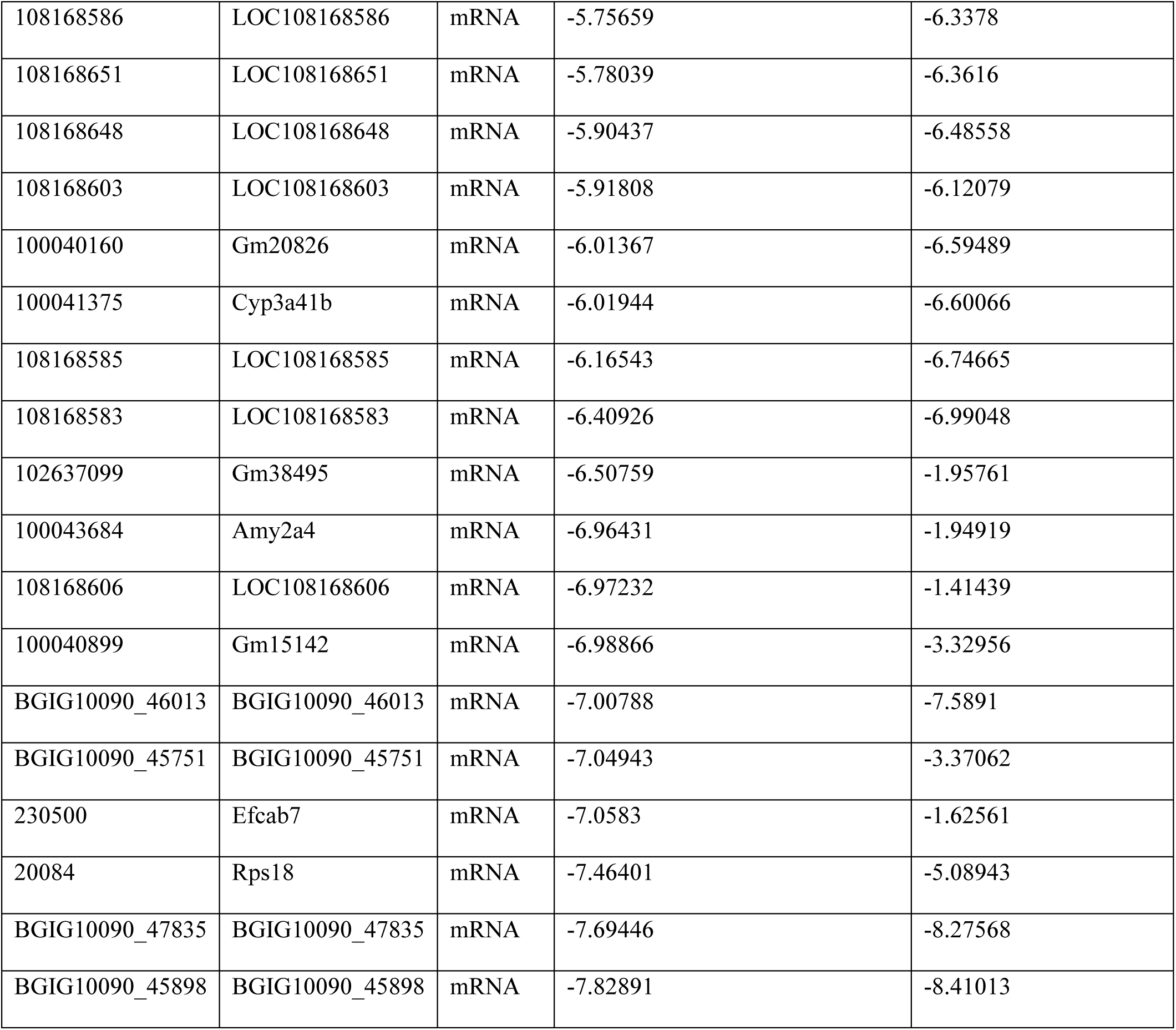
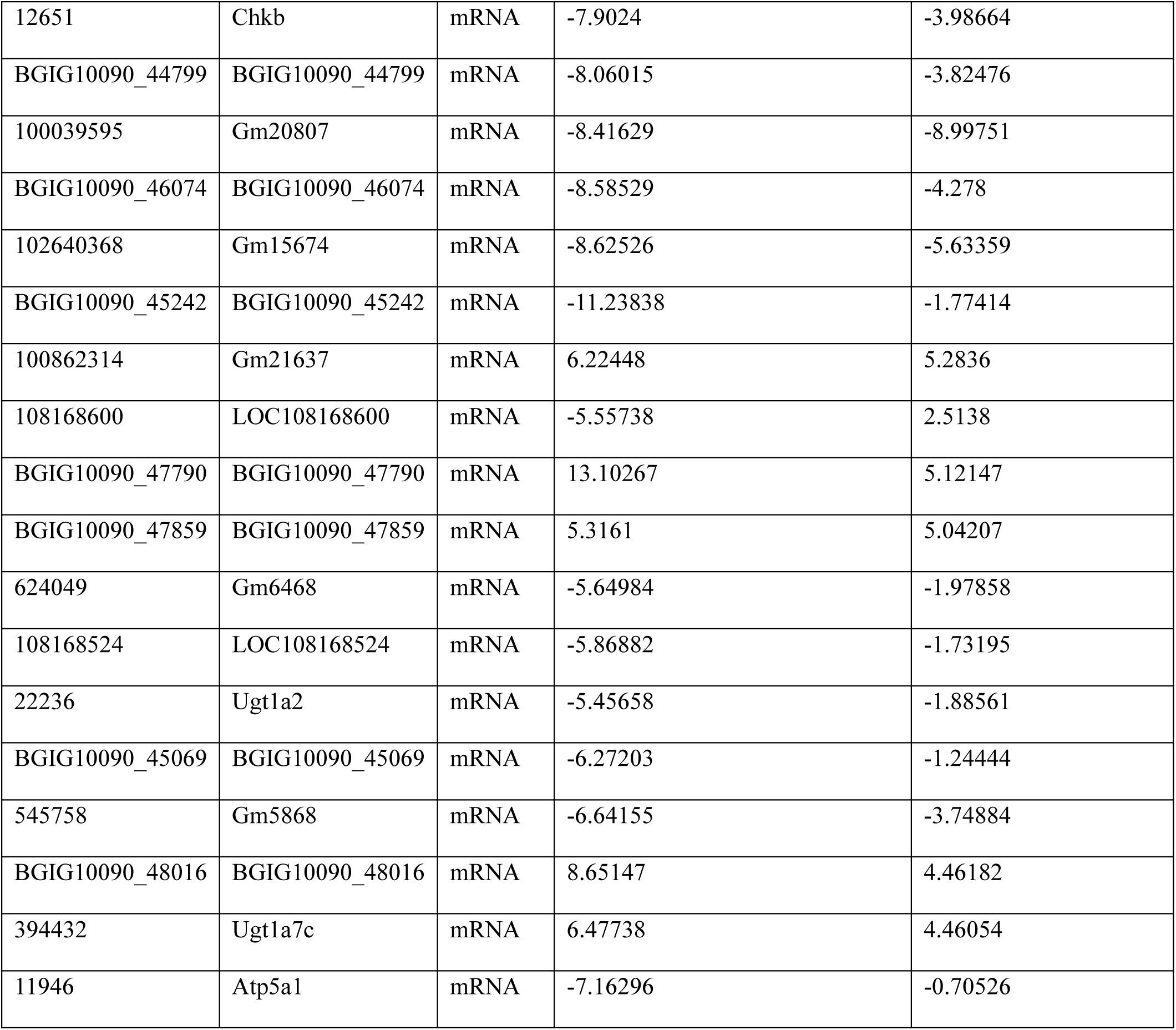
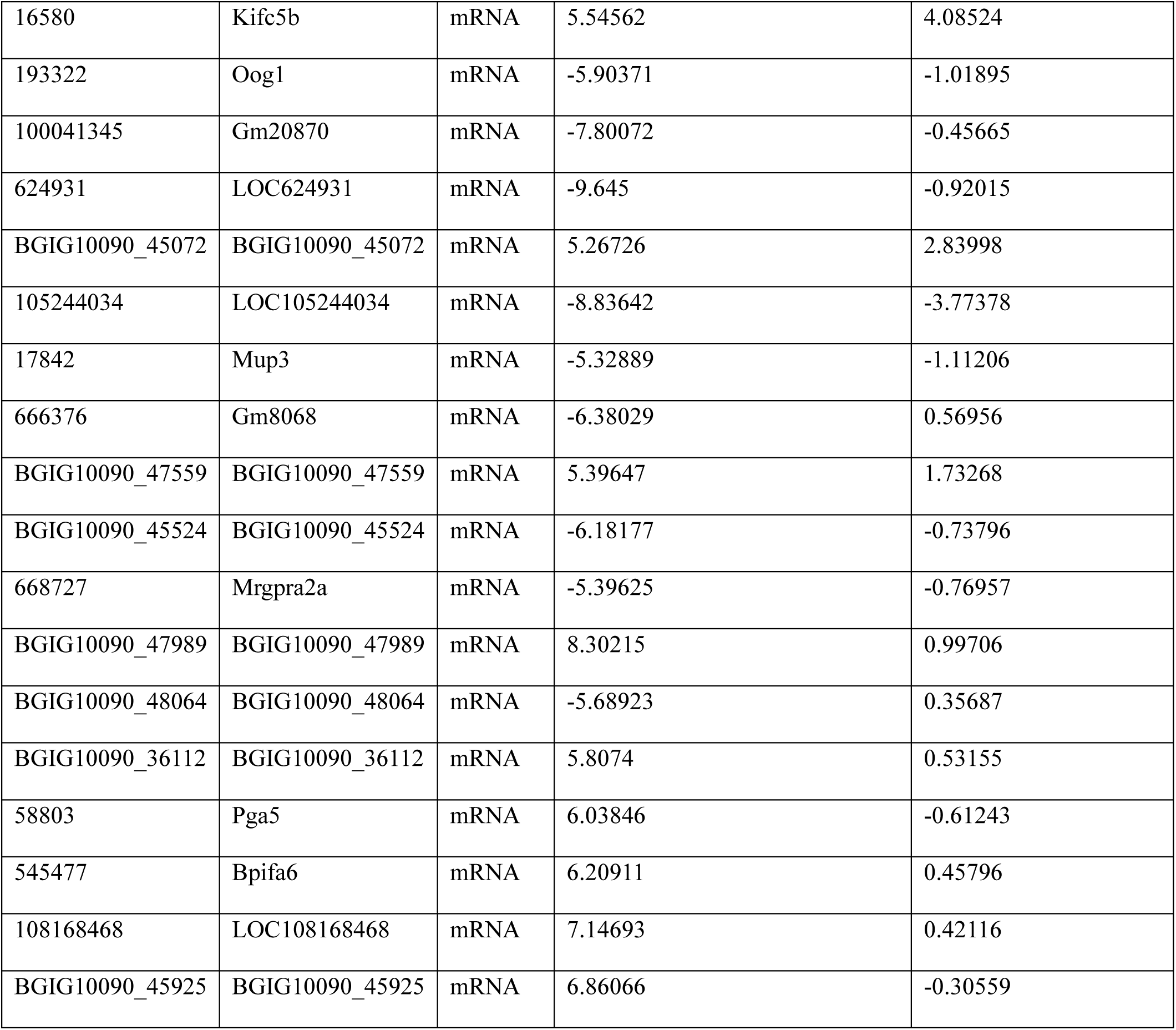
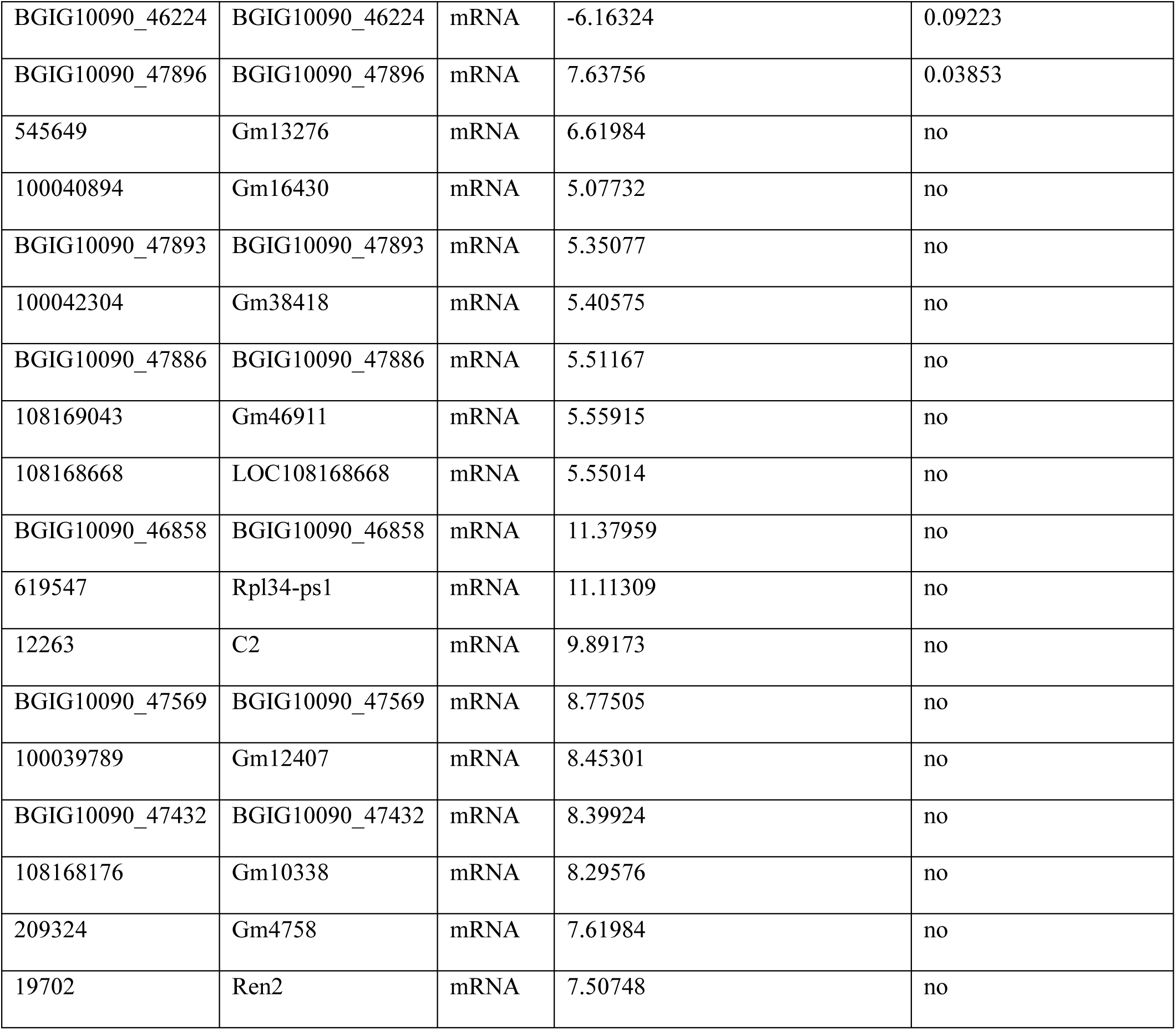
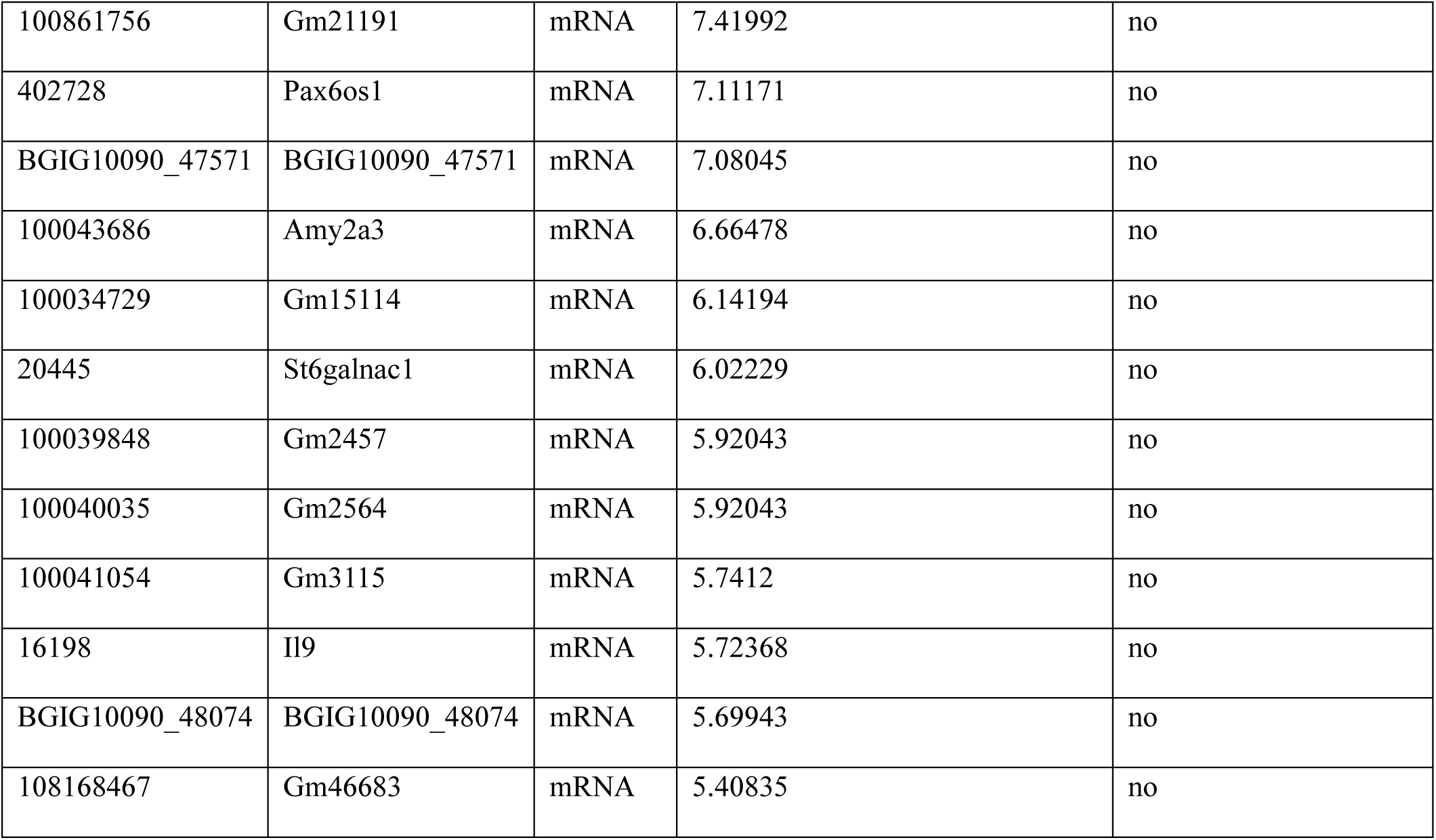

**Supply Dataset S5-1.**

| Group | concentration($10^6$/ML) | Sperm mobility(%) | Progressive mobile sperm(%) |
| --- | --- | --- | --- |
| Normal | 46.67 | 70.484 | 69.776 |
| Normal | 27.395 | 68.584 | 67.994 |
| Normal | 40.83 | 62.113 | 61.293 |
| Normal | 42.86 | 70.125 | 68.125 |
| Normal | 39.5 | 64.531 | 63.087 |
| Normal | 29.095 | 63.312 | 62.683 |
| Oligoasthenospermia | 8.89 | 16.536 | 16.142 |
| Oligoasthenospermia | 9.985 | 32.407 | 29.938 |
| Oligoasthenospermia | 10.145 | 31.959 | 29.897 |
| Oligoasthenospermia | 8.71 | 33.811 | 31.805 |
| Oligoasthenospermia | 10.235 | 31.637 | 30.182 |
| Oligoasthenospermia | 7.73 | 24.813 | 30.488 |
| Schisandrin B | 61.54 | 70.035 | 69.085 |
| Schisandrin B | 27.195 | 50.474 | 50.237 |
| Schisandrin B | 29.735 | 48.831 | 47.792 |
| Schisandrin B | 90.39 | 87.414 | 86.726 |
| Schisandrin B | 31.015 | 59.498 | 58.173 |
| Schisandrin B | 55.92 | 78.547 | 77.832 |
| Wuzi-Pills | 13.905 | 43.956 | 43.956 |
| Wuzi-Pills | 24.17 | 53.56 | 52.204 |
| Wuzi-Pills | 21.285 | 26.751 | 26.212 |
| Wuzi-Pills | 27.7 | 47.887 | 46.197 |
| Wuzi-Pills | 52.14 | 78.838 | 78.285 |
| Wuzi-Pills | 26.63 | 44.637 | 43.153 |
| Testosterone propionate | 13.735 | 27.5 | 26.429 |
| Testosterone propionate | 34.46 | 71.243 | 70.686 |
| Testosterone propionate | 10.375 | 39.759 | 39.157 |
| Testosterone propionate | 14.76 | 23.454 | 22.939 |
| Testosterone propionate | 7.38 | 16.713 | 13.314 |
| Testosterone propionate | 21.285 | 21.122 | 22.066 |

**Supply Dataset S5-2.**
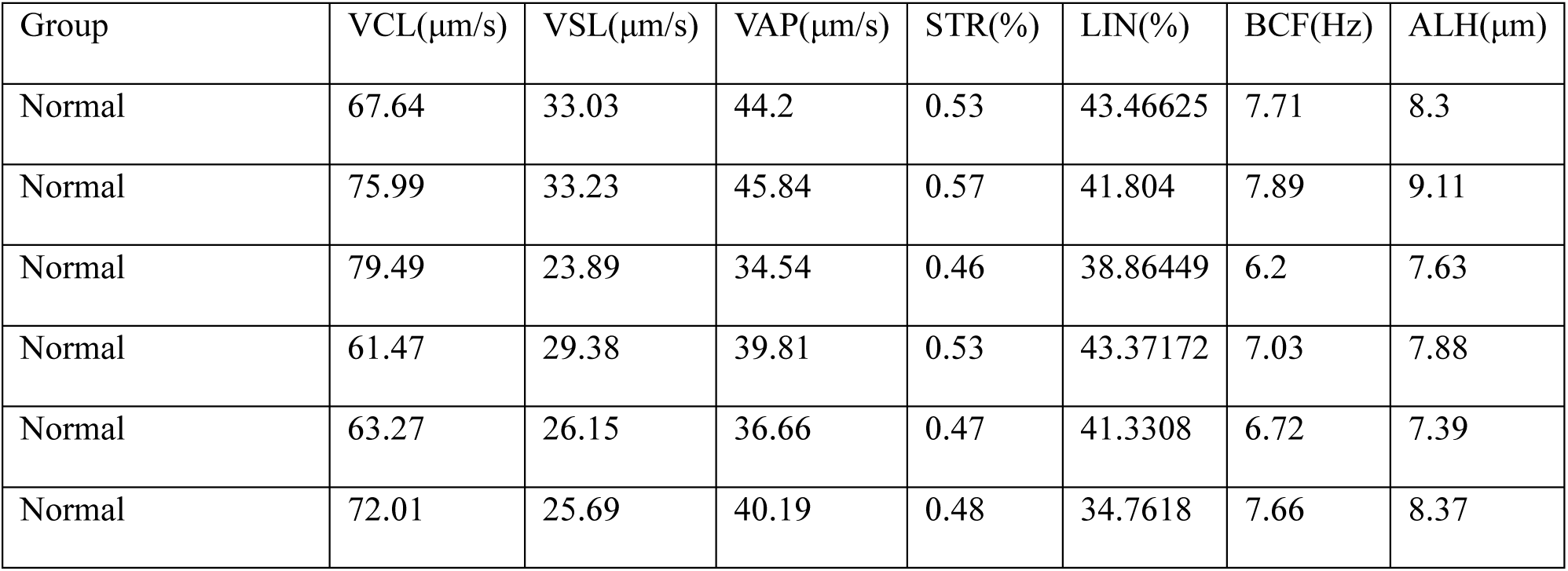
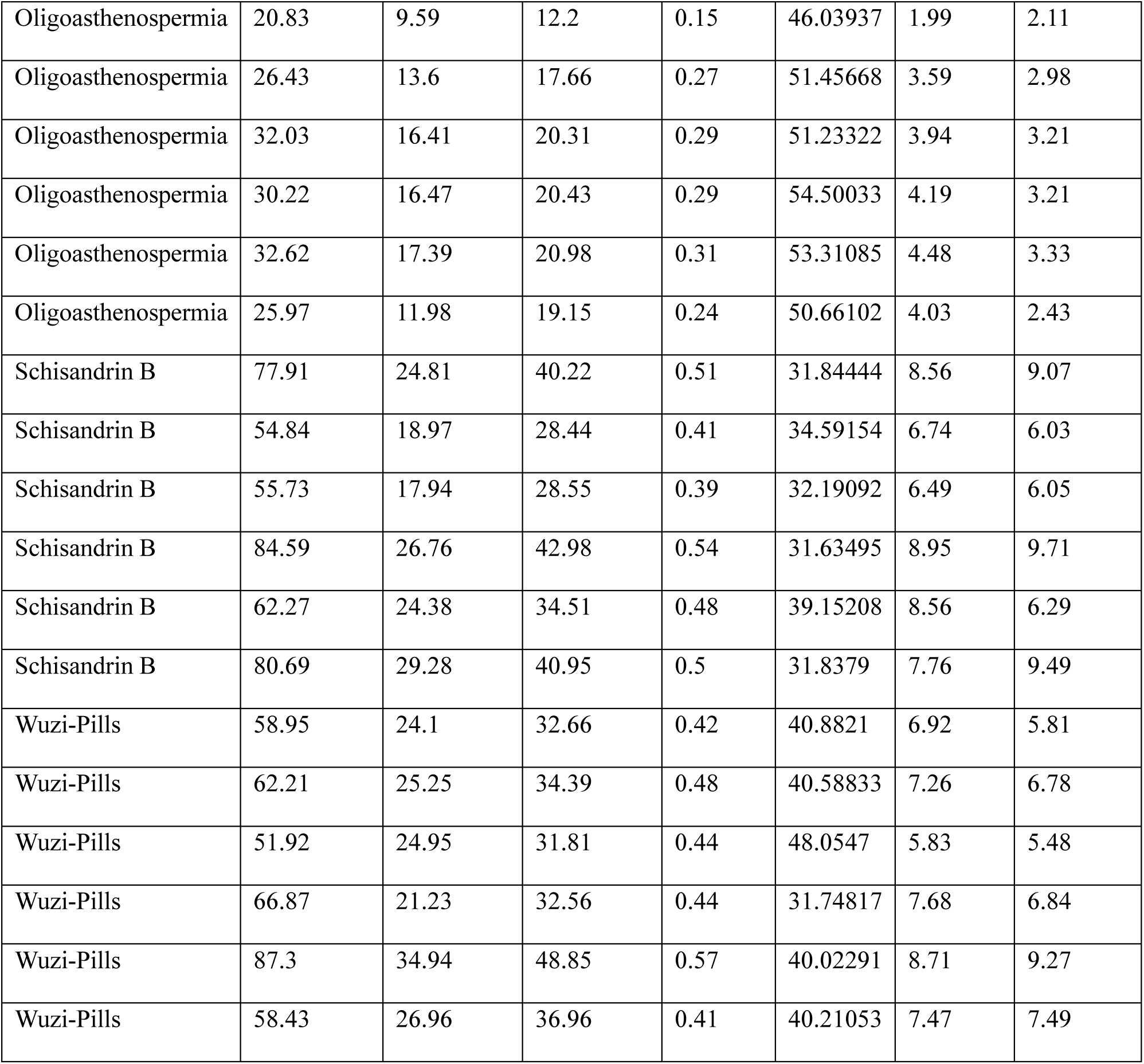
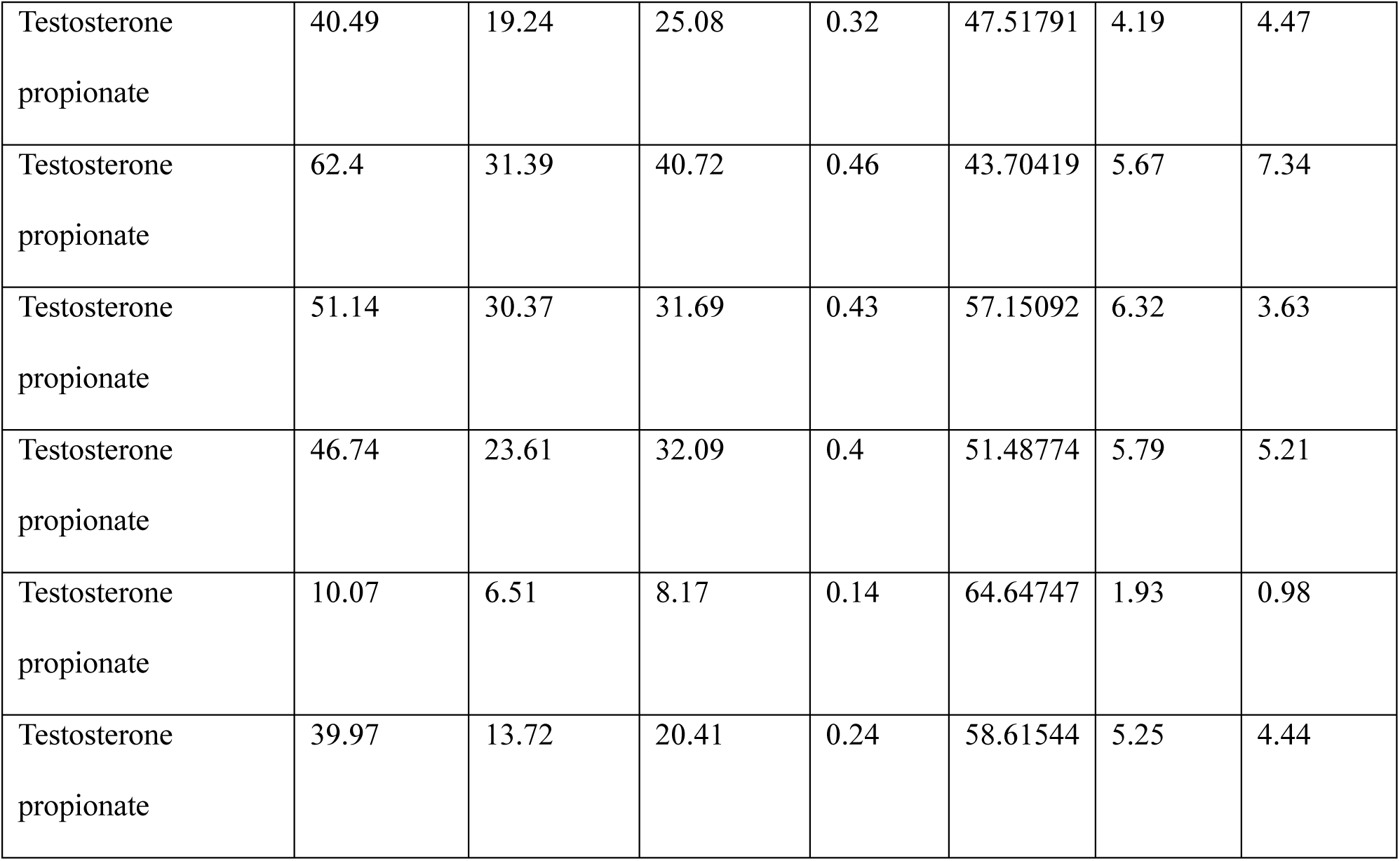

| Group | VCL($\mu\text{m/s}$) | VSL($\mu\text{m/s}$) | VAP($\mu\text{m/s}$) | STR(%) | LIN(%) | BCF(Hz) | ALH($\mu\text{m}$) |
| --- | --- | --- | --- | --- | --- | --- | --- |
| Normal | 67.64 | 33.03 | 44.2 | 0.53 | 43.46625 | 7.71 | 8.3 |
| Normal | 75.99 | 33.23 | 45.84 | 0.57 | 41.804 | 7.89 | 9.11 |
| Normal | 79.49 | 23.89 | 34.54 | 0.46 | 38.86449 | 6.2 | 7.63 |
| Normal | 61.47 | 29.38 | 39.81 | 0.53 | 43.37172 | 7.03 | 7.88 |
| Normal | 63.27 | 26.15 | 36.66 | 0.47 | 41.3308 | 6.72 | 7.39 |
| Normal | 72.01 | 25.69 | 40.19 | 0.48 | 34.7618 | 7.66 | 8.37 |
| Oligoasthenospermia | 20.83 | 9.59 | 12.2 | 0.15 | 46.03937 | 1.99 | 2.11 |
| Oligoasthenospermia | 26.43 | 13.6 | 17.66 | 0.27 | 51.45668 | 3.59 | 2.98 |
| Oligoasthenospermia | 32.03 | 16.41 | 20.31 | 0.29 | 51.23322 | 3.94 | 3.21 |
| Oligoasthenospermia | 30.22 | 16.47 | 20.43 | 0.29 | 54.50033 | 4.19 | 3.21 |
| Oligoasthenospermia | 32.62 | 17.39 | 20.98 | 0.31 | 53.31085 | 4.48 | 3.33 |
| Oligoasthenospermia | 25.97 | 11.98 | 19.15 | 0.24 | 50.66102 | 4.03 | 2.43 |
| Schisandrin B | 77.91 | 24.81 | 40.22 | 0.51 | 31.84444 | 8.56 | 9.07 |
| Schisandrin B | 54.84 | 18.97 | 28.44 | 0.41 | 34.59154 | 6.74 | 6.03 |
| Schisandrin B | 55.73 | 17.94 | 28.55 | 0.39 | 32.19092 | 6.49 | 6.05 |
| Schisandrin B | 84.59 | 26.76 | 42.98 | 0.54 | 31.63495 | 8.95 | 9.71 |
| Schisandrin B | 62.27 | 24.38 | 34.51 | 0.48 | 39.15208 | 8.56 | 6.29 |
| Schisandrin B | 80.69 | 29.28 | 40.95 | 0.5 | 31.8379 | 7.76 | 9.49 |
| Wuzi-Pills | 58.95 | 24.1 | 32.66 | 0.42 | 40.8821 | 6.92 | 5.81 |
| Wuzi-Pills | 62.21 | 25.25 | 34.39 | 0.48 | 40.58833 | 7.26 | 6.78 |
| Wuzi-Pills | 51.92 | 24.95 | 31.81 | 0.44 | 48.0547 | 5.83 | 5.48 |
| Wuzi-Pills | 66.87 | 21.23 | 32.56 | 0.44 | 31.74817 | 7.68 | 6.84 |
| Wuzi-Pills | 87.3 | 34.94 | 48.85 | 0.57 | 40.02291 | 8.71 | 9.27 |
| Wuzi-Pills | 58.43 | 26.96 | 36.96 | 0.41 | 40.21053 | 7.47 | 7.49 |
| Testosterone<br>propionate | 40.49 | 19.24 | 25.08 | 0.32 | 47.51791 | 4.19 | 4.47 |
| Testosterone<br>propionate | 62.4 | 31.39 | 40.72 | 0.46 | 43.70419 | 5.67 | 7.34 |
| Testosterone<br>propionate | 51.14 | 30.37 | 31.69 | 0.43 | 57.15092 | 6.32 | 3.63 |
| Testosterone<br>propionate | 46.74 | 23.61 | 32.09 | 0.4 | 51.48774 | 5.79 | 5.21 |
| Testosterone<br>propionate | 10.07 | 6.51 | 8.17 | 0.14 | 64.64747 | 1.93 | 0.98 |
| Testosterone<br>propionate | 39.97 | 13.72 | 20.41 | 0.24 | 58.61544 | 5.25 | 4.44 |

**Supply Dataset S6.**
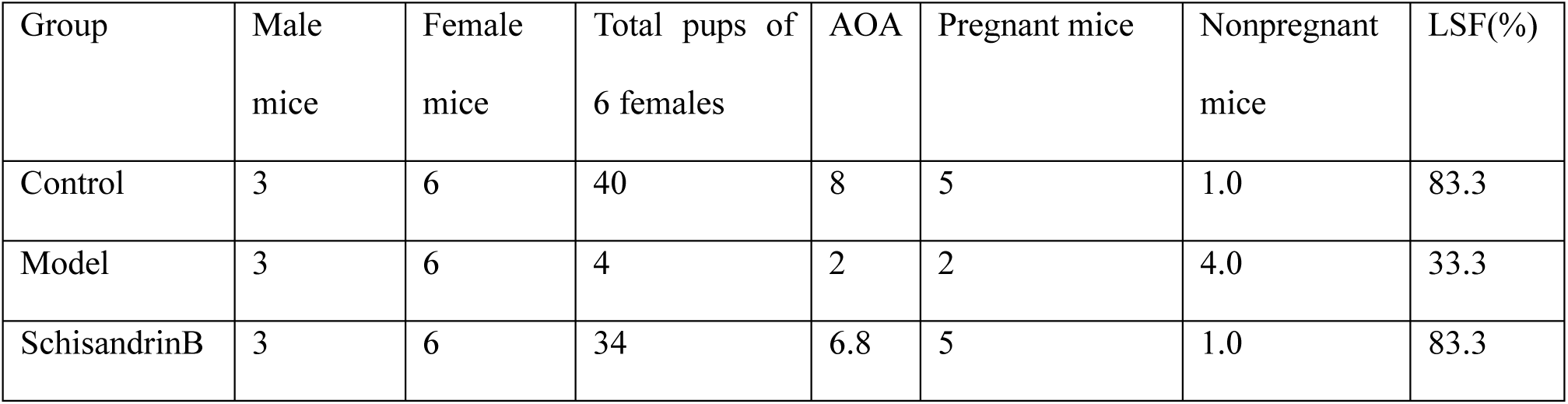
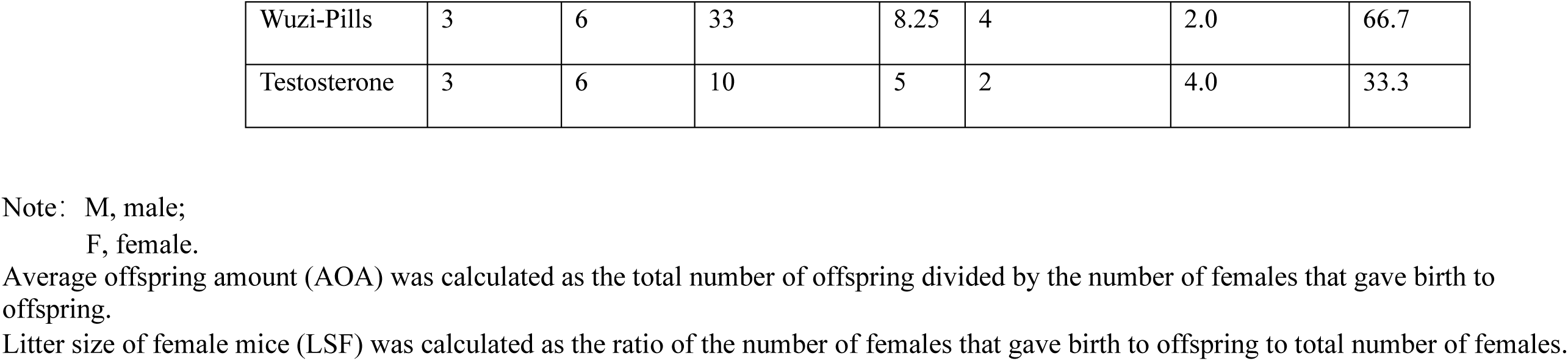

**Supply Dataset S7.**
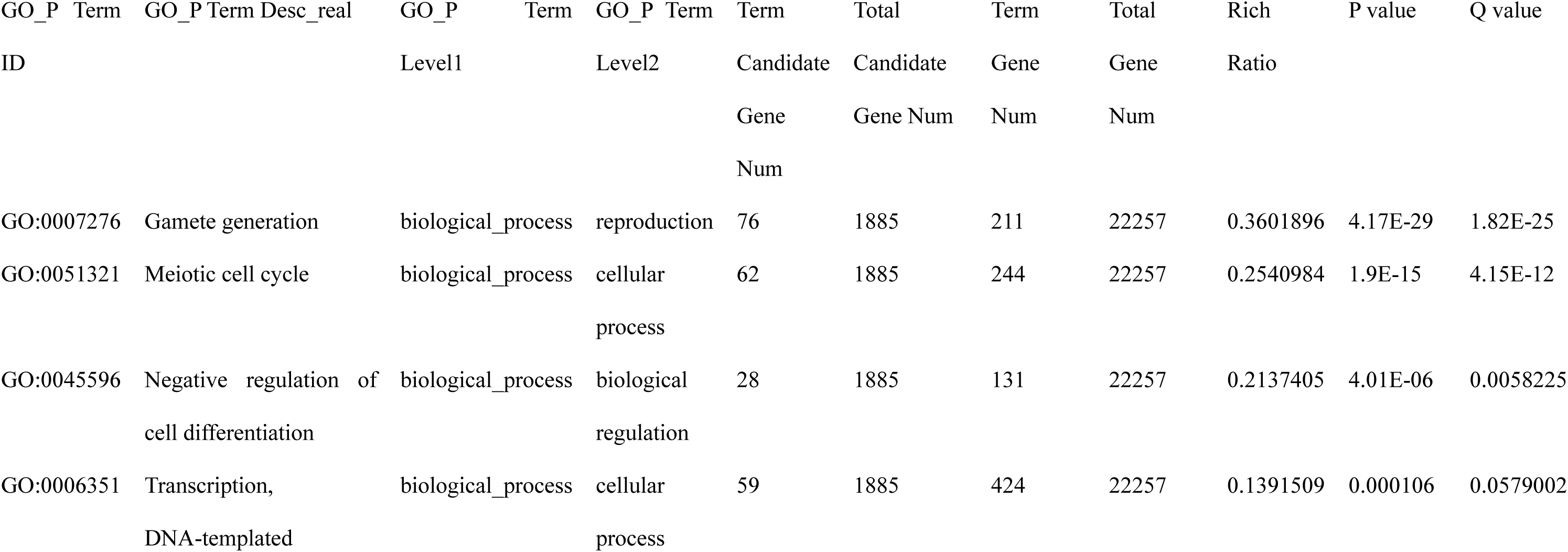
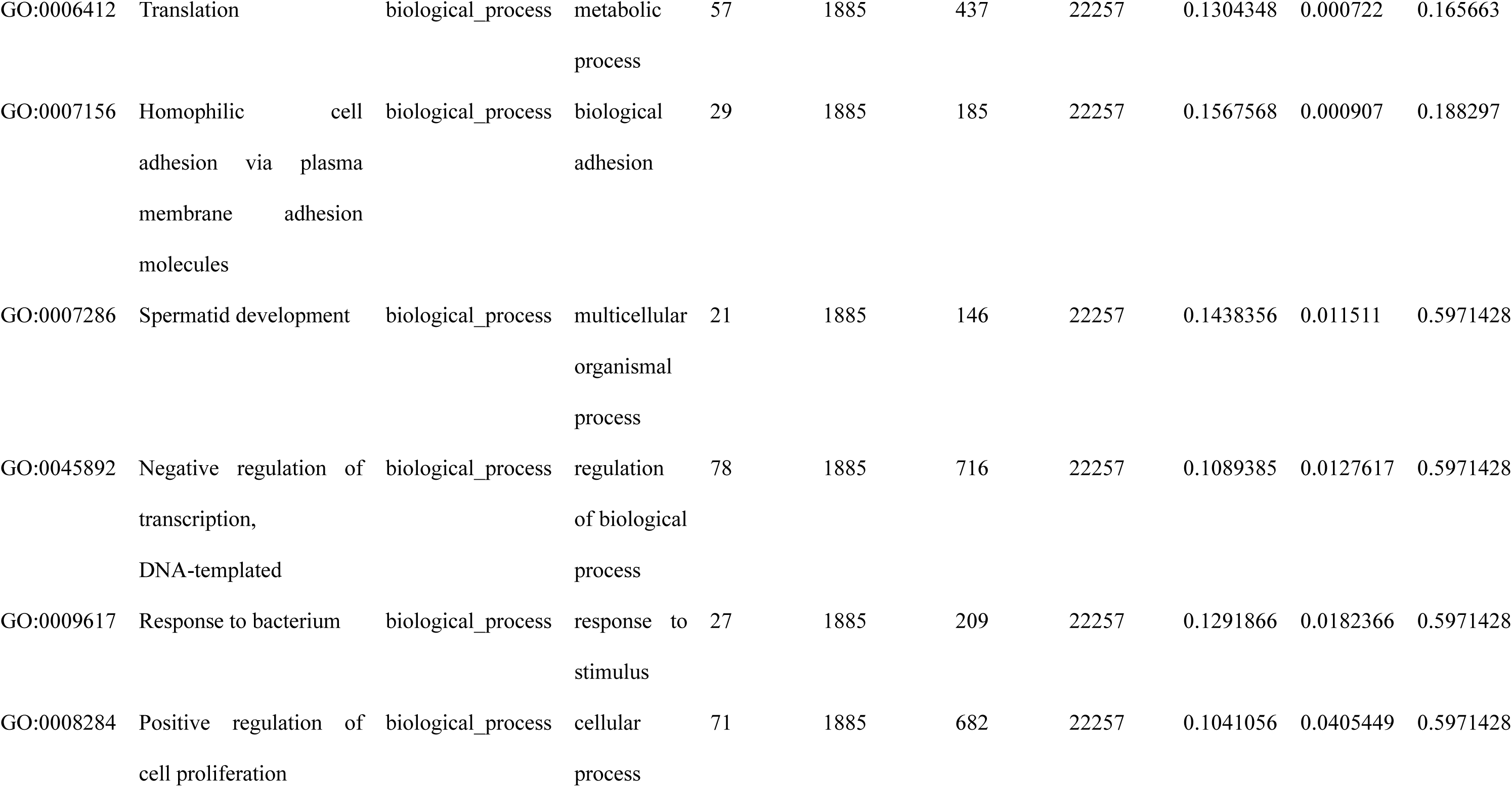

**Supply Dataset S8.**
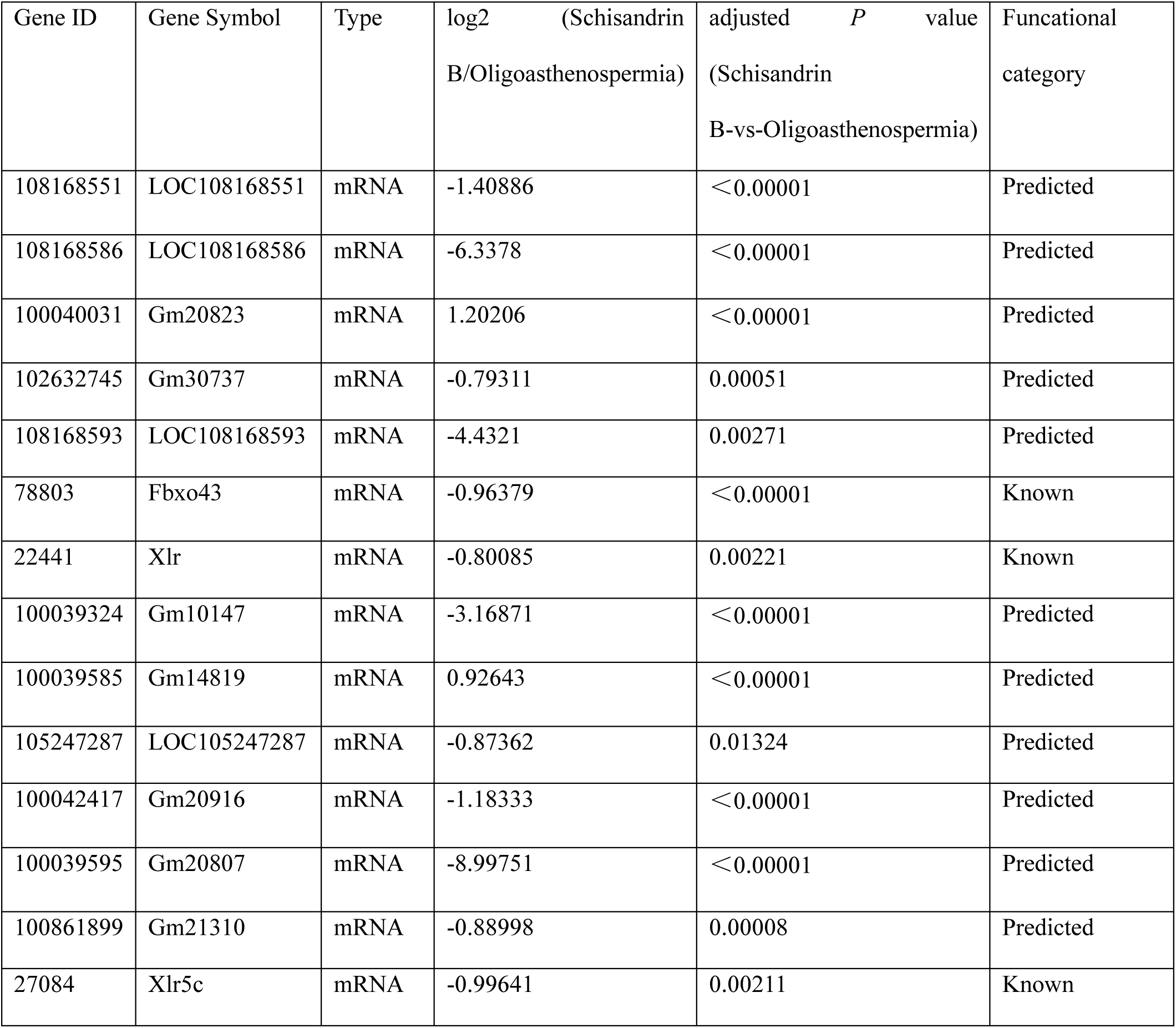
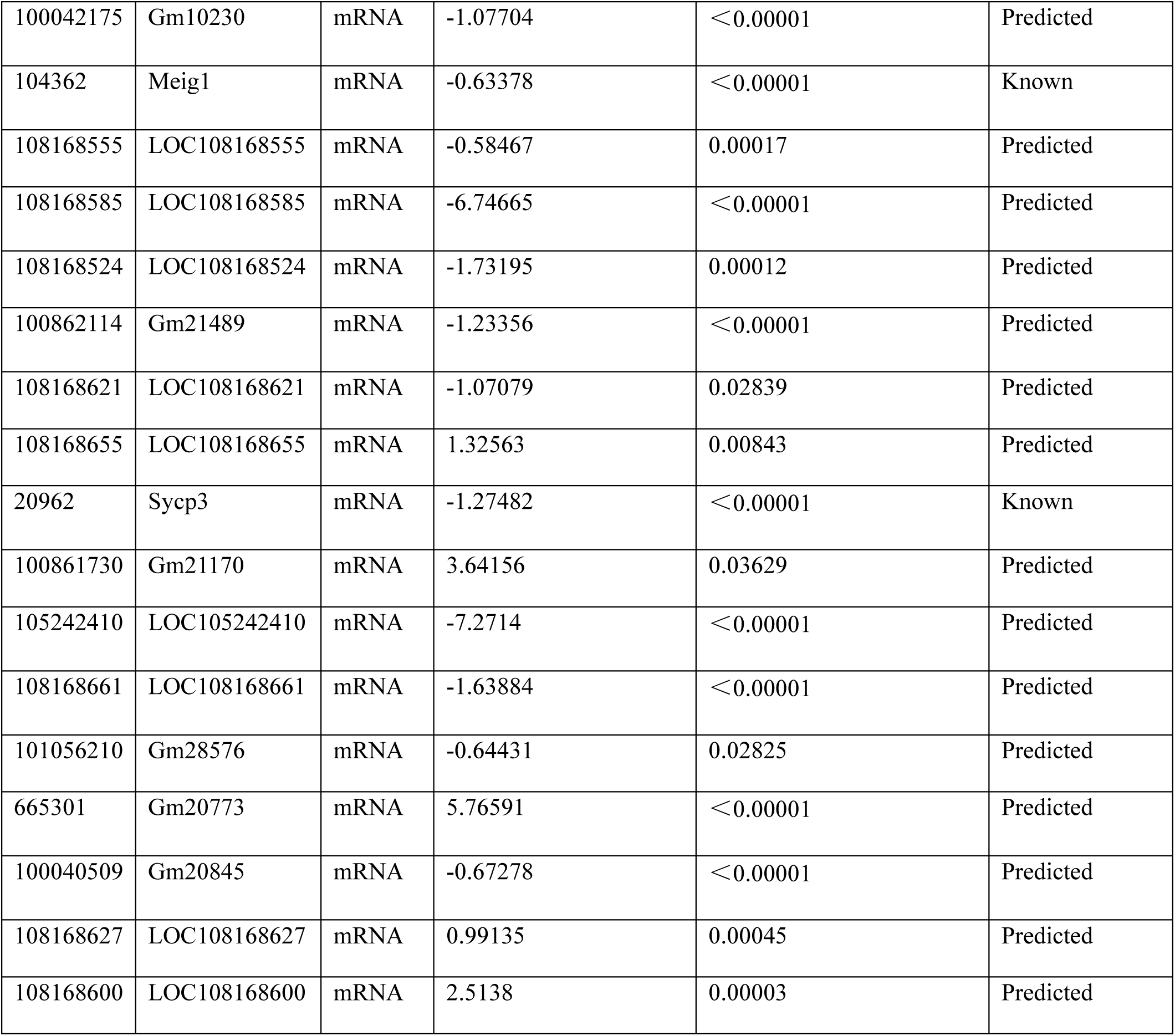
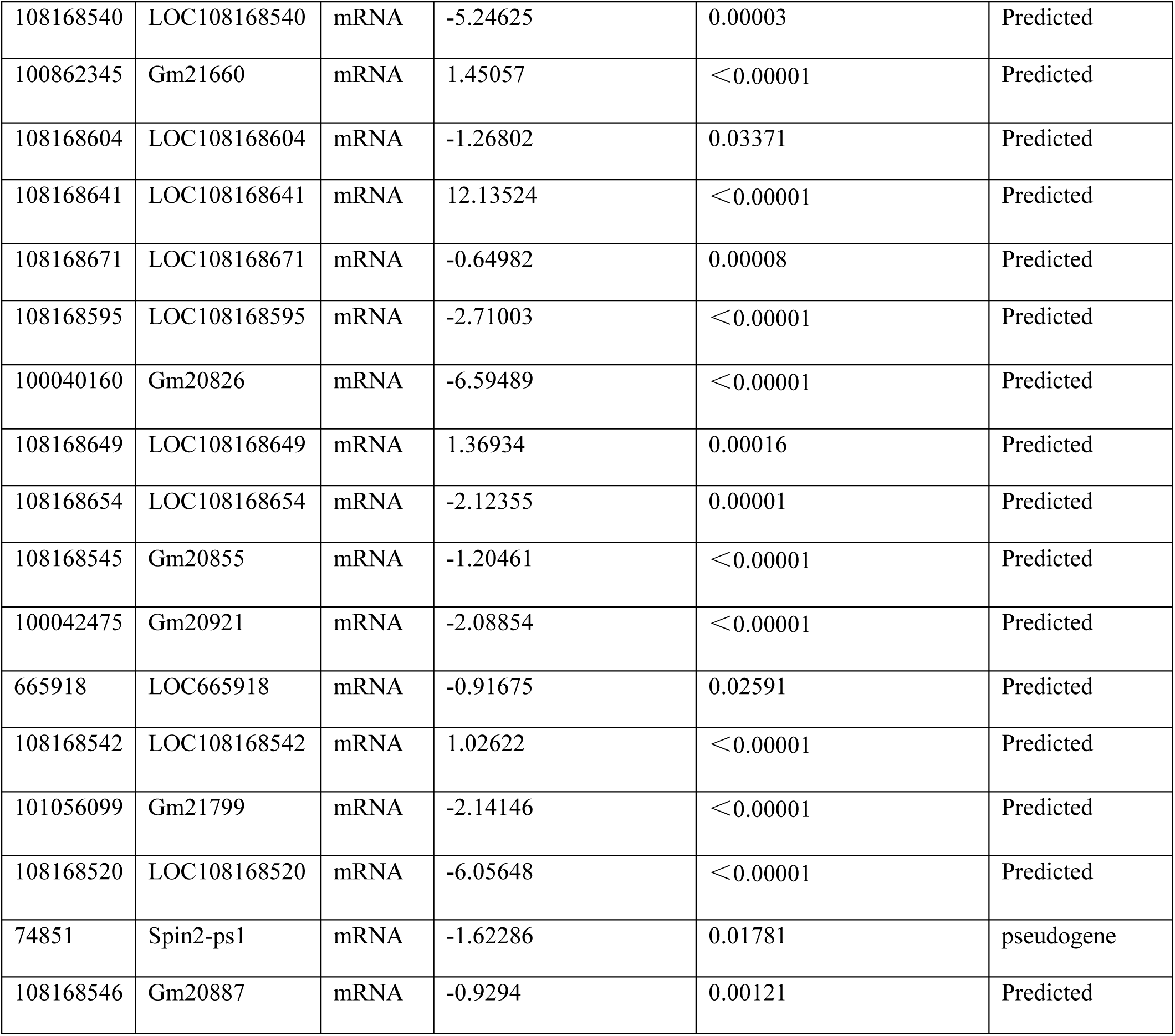
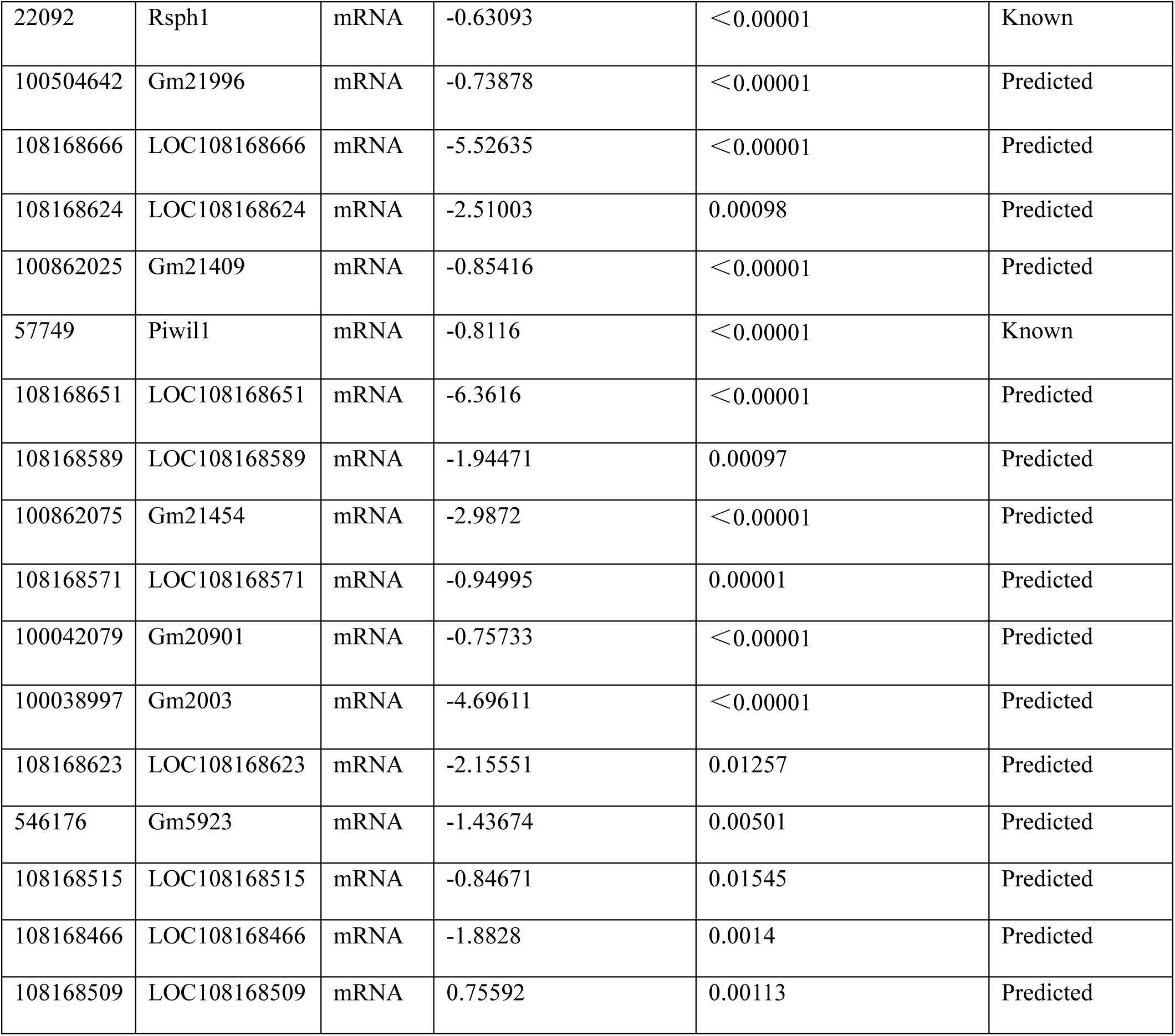
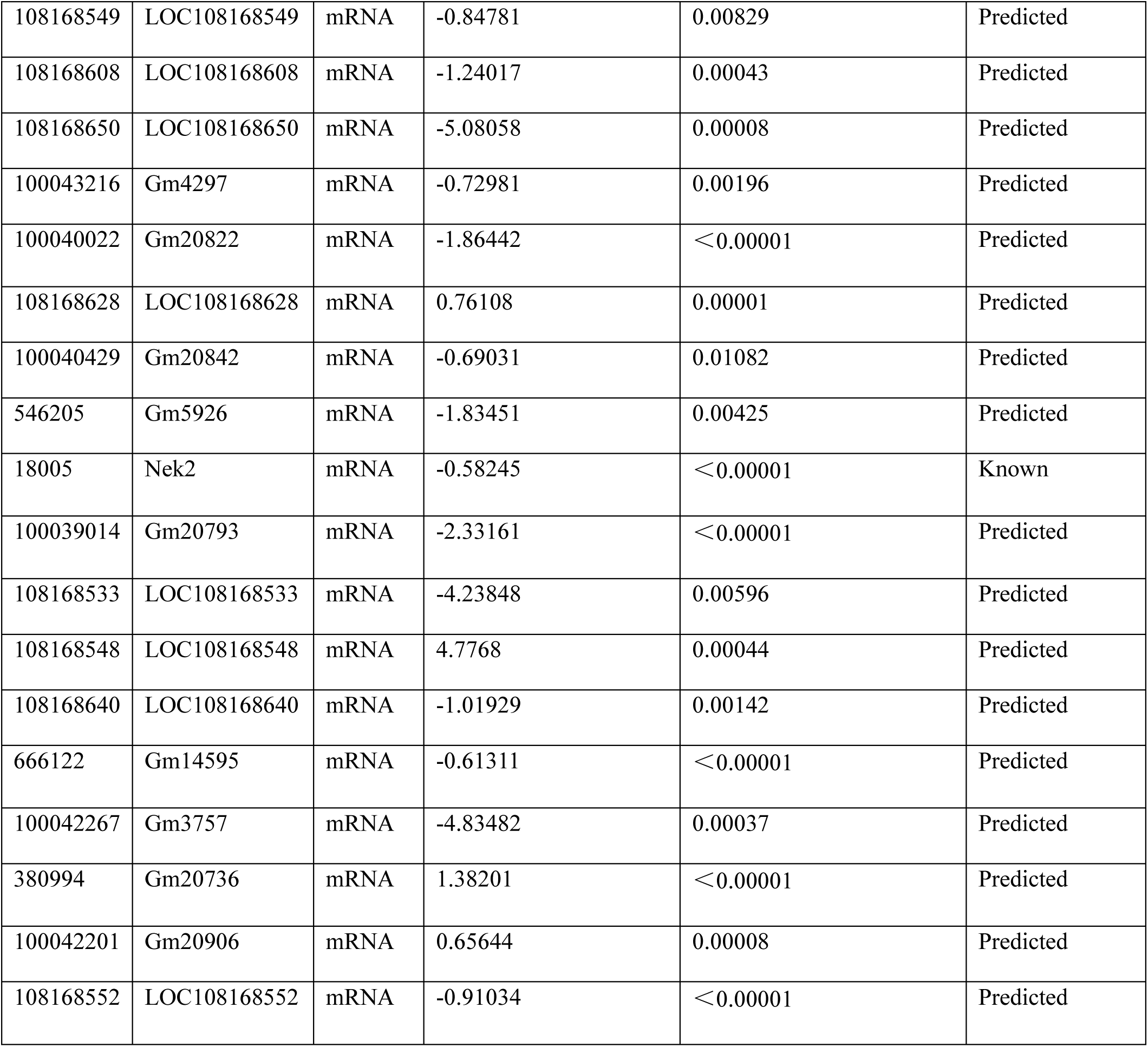
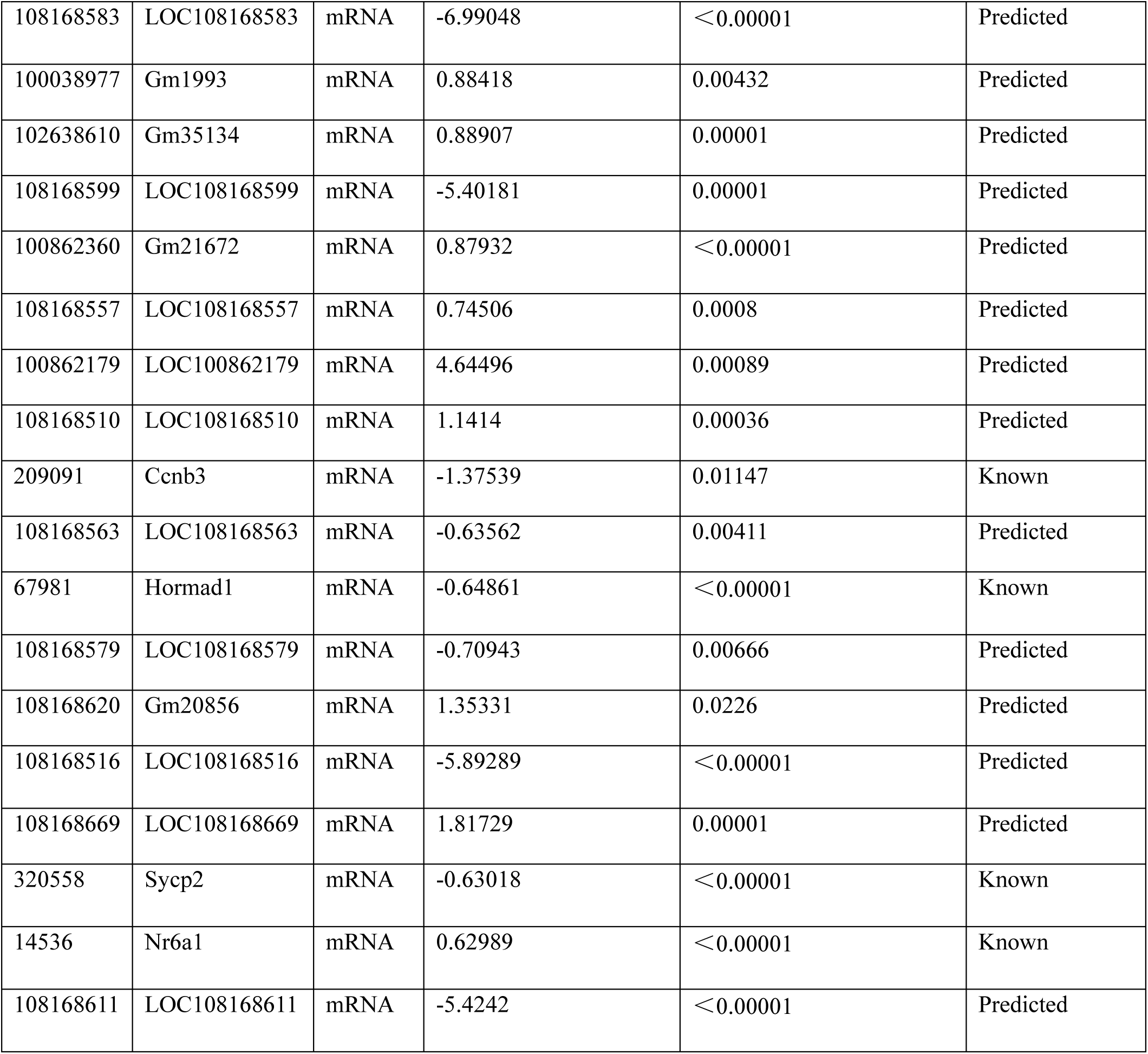
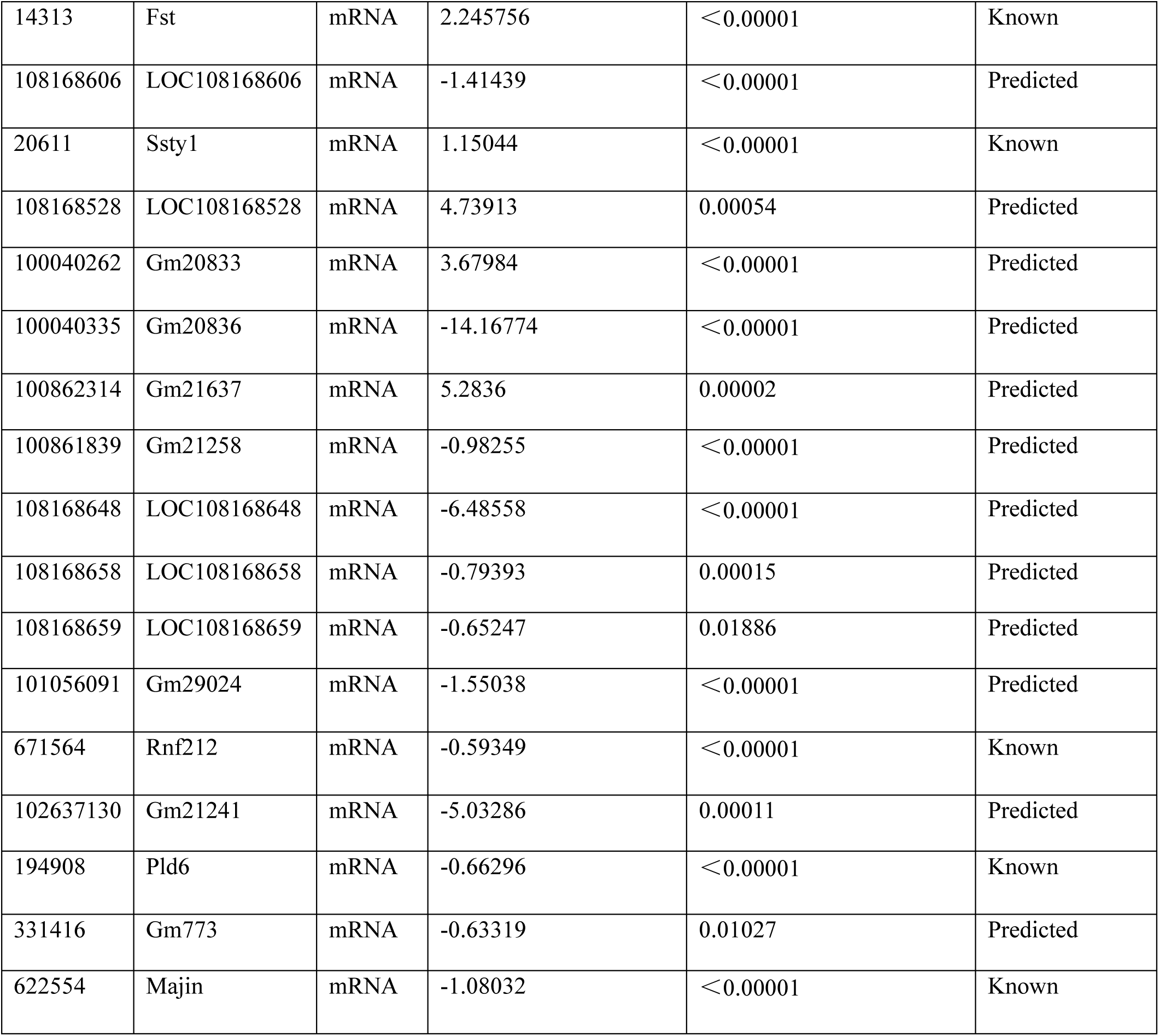
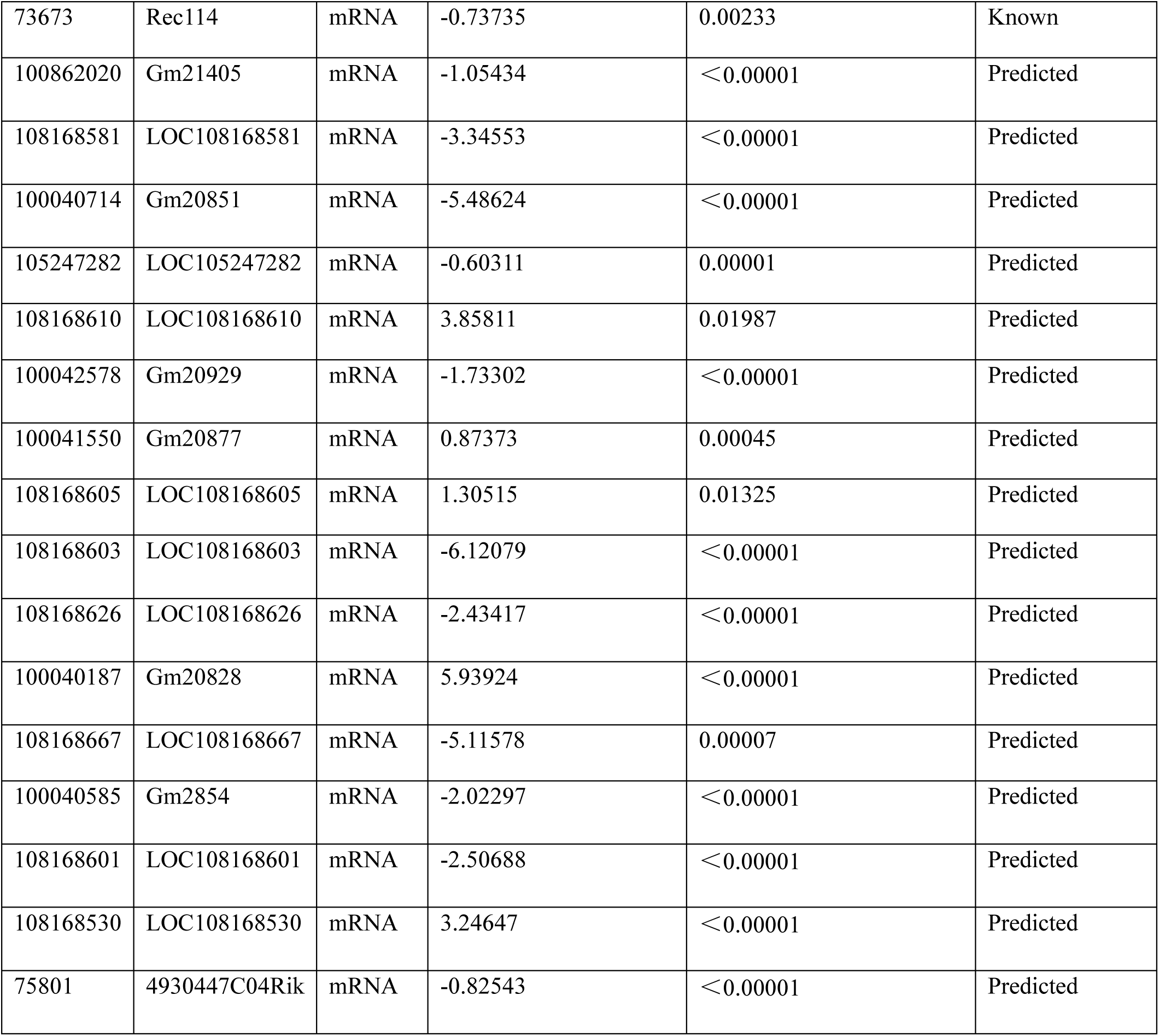
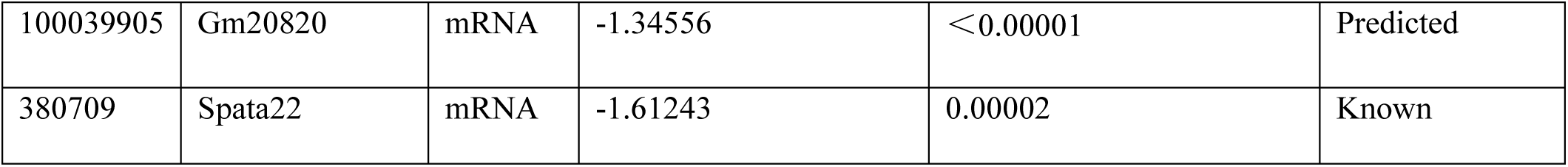

**Supply Dataset S9.**
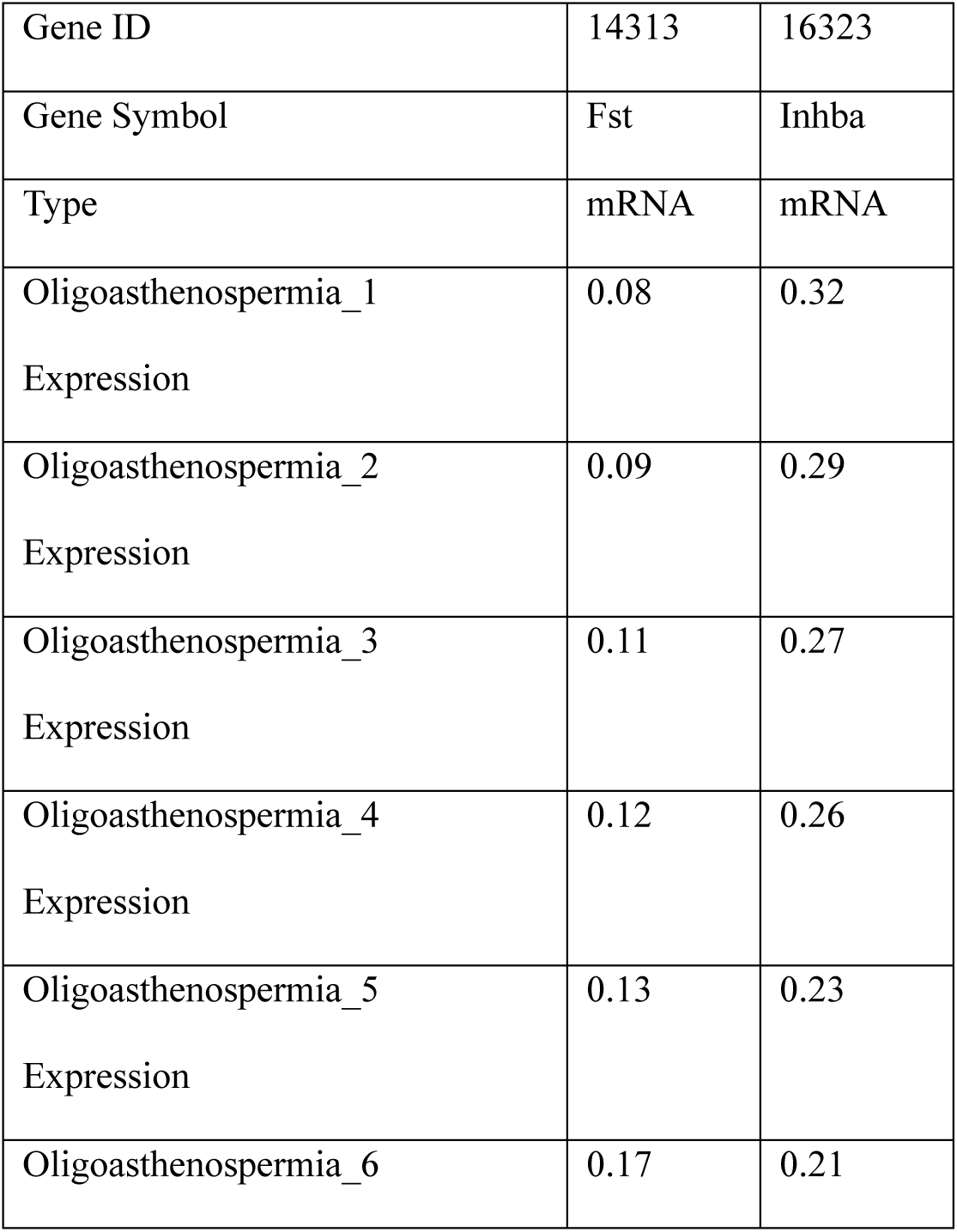
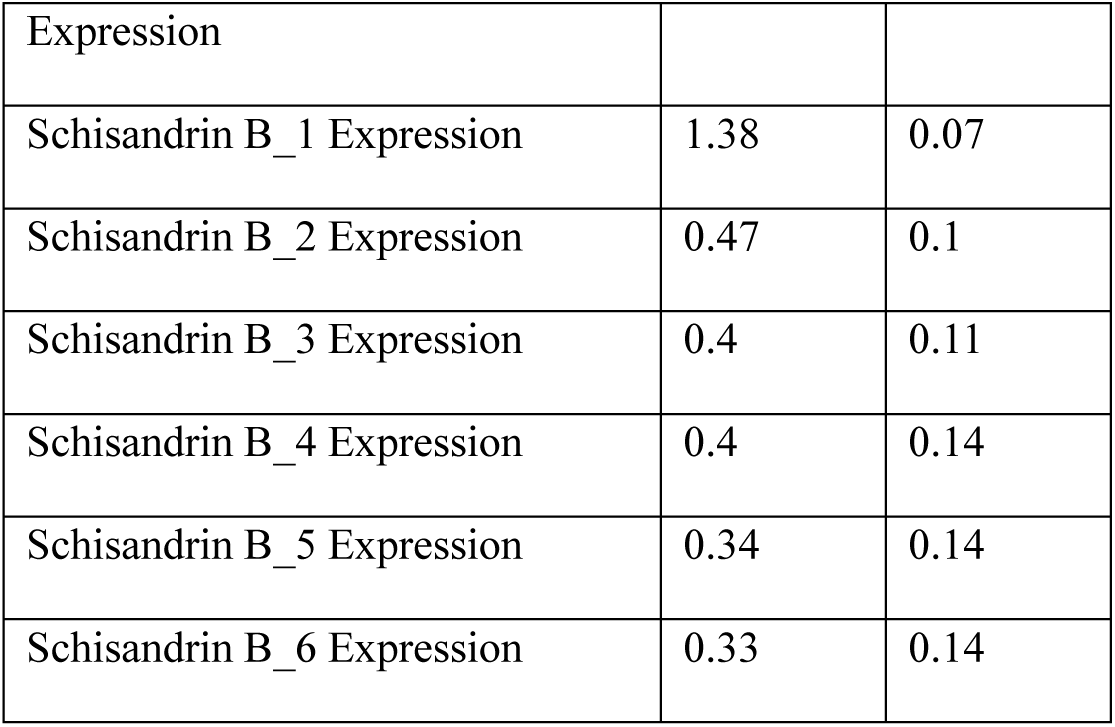

**Supply Dataset S10-1.**
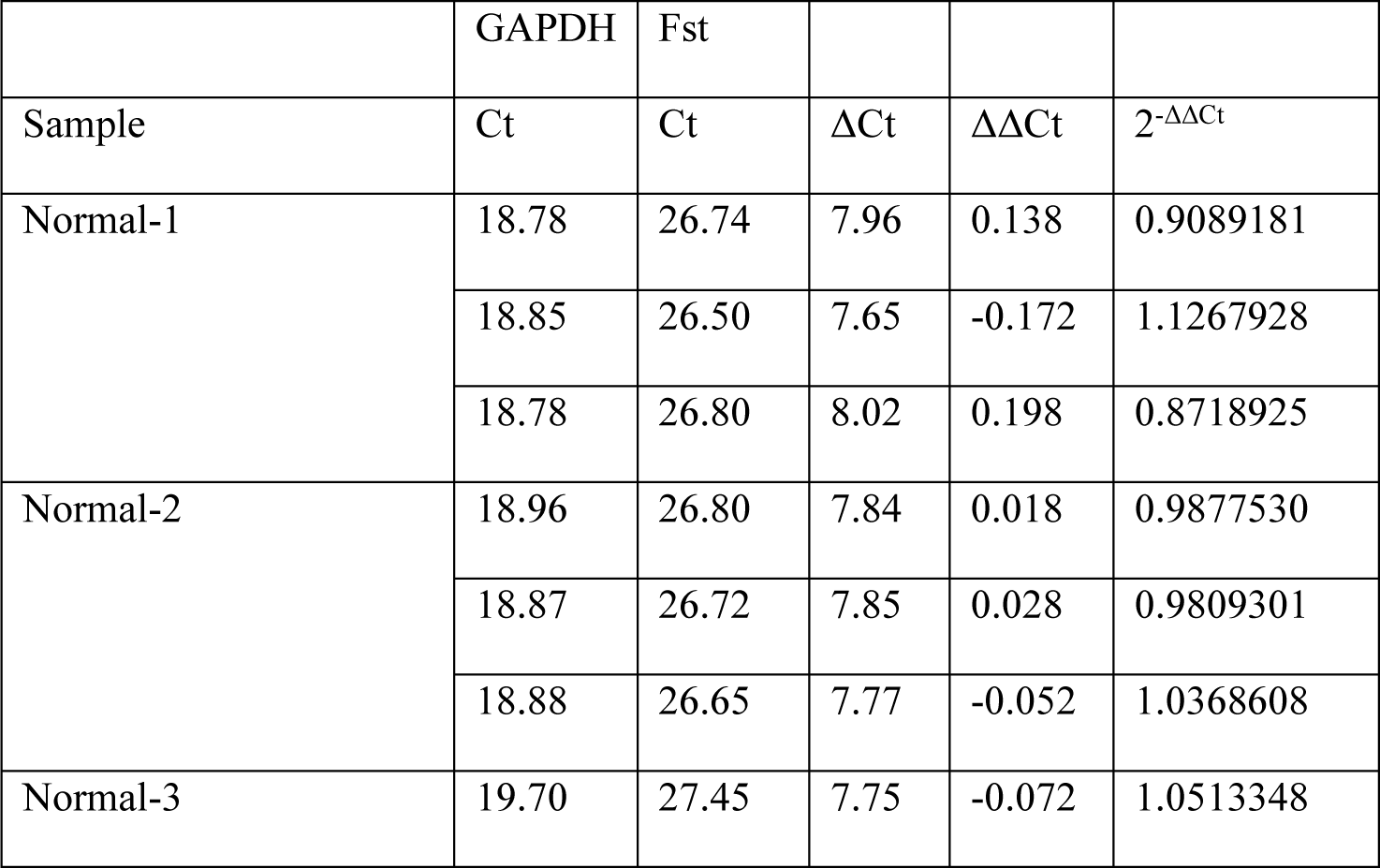
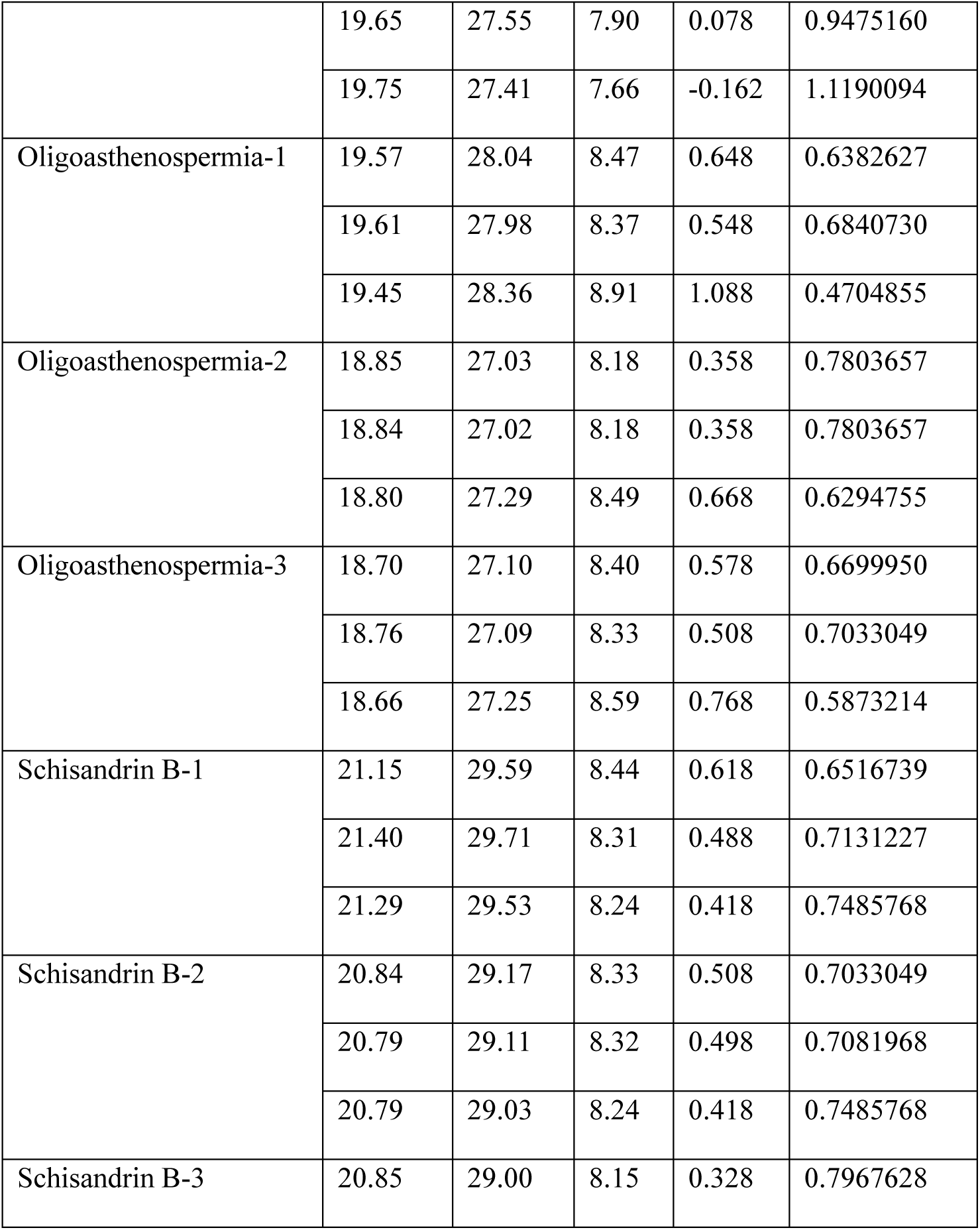
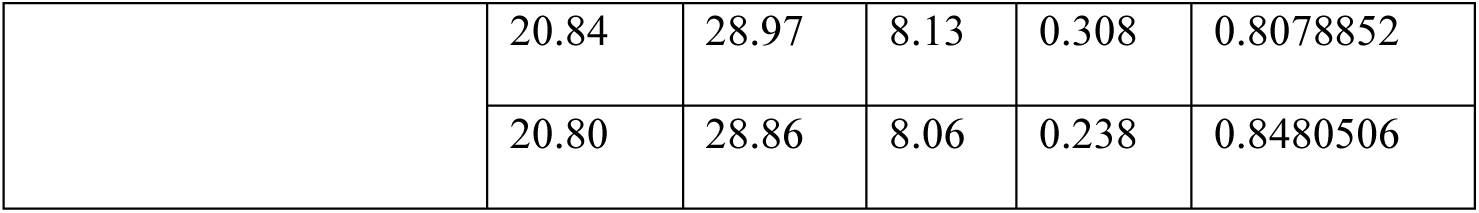

|  | GAPDH | Fst |  |  |  |
| --- | --- | --- | --- | --- | --- |
| Sample | Ct | Ct | $\Delta Ct$ | $\Delta\Delta Ct$ | $2^{-\Delta\Delta Ct}$ |
| Normal-1 | 18.78 | 26.74 | 7.96 | 0.138 | 0.9089181 |
|  | 18.85 | 26.50 | 7.65 | -0.172 | 1.1267928 |
|  | 18.78 | 26.80 | 8.02 | 0.198 | 0.8718925 |
| Normal-2 | 18.96 | 26.80 | 7.84 | 0.018 | 0.9877530 |
|  | 18.87 | 26.72 | 7.85 | 0.028 | 0.9809301 |
|  | 18.88 | 26.65 | 7.77 | -0.052 | 1.0368608 |
| Normal-3 | 19.70 | 27.45 | 7.75 | -0.072 | 1.0513348 |
|  | 19.65 | 27.55 | 7.90 | 0.078 | 0.9475160 |
|  | 19.75 | 27.41 | 7.66 | -0.162 | 1.1190094 |
| Oligoasthenospermia-1 | 19.57 | 28.04 | 8.47 | 0.648 | 0.6382627 |
|  | 19.61 | 27.98 | 8.37 | 0.548 | 0.6840730 |
|  | 19.45 | 28.36 | 8.91 | 1.088 | 0.4704855 |
| Oligoasthenospermia-2 | 18.85 | 27.03 | 8.18 | 0.358 | 0.7803657 |
|  | 18.84 | 27.02 | 8.18 | 0.358 | 0.7803657 |
|  | 18.80 | 27.29 | 8.49 | 0.668 | 0.6294755 |
| Oligoasthenospermia-3 | 18.70 | 27.10 | 8.40 | 0.578 | 0.6699950 |
|  | 18.76 | 27.09 | 8.33 | 0.508 | 0.7033049 |
|  | 18.66 | 27.25 | 8.59 | 0.768 | 0.5873214 |
| Schisandrin B-1 | 21.15 | 29.59 | 8.44 | 0.618 | 0.6516739 |
|  | 21.40 | 29.71 | 8.31 | 0.488 | 0.7131227 |
|  | 21.29 | 29.53 | 8.24 | 0.418 | 0.7485768 |
| Schisandrin B-2 | 20.84 | 29.17 | 8.33 | 0.508 | 0.7033049 |
|  | 20.79 | 29.11 | 8.32 | 0.498 | 0.7081968 |
|  | 20.79 | 29.03 | 8.24 | 0.418 | 0.7485768 |
| Schisandrin B-3 | 20.85 | 29.00 | 8.15 | 0.328 | 0.7967628 |
|  | 20.84 | 28.97 | 8.13 | 0.308 | 0.8078852 |
|  | 20.80 | 28.86 | 8.06 | 0.238 | 0.8480506 |

**Supply Dataset S10-2.**

|  | GAPDH | Inhba |  |  |  |
| --- | --- | --- | --- | --- | --- |
| Sample | Ct | Ct | $\Delta Ct$ | $\Delta\Delta Ct$ | $2^{-\Delta\Delta Ct}$ |
| Normal-1 | 18.78 | 27.95 | 9.17 | 0.349 | 0.7851886 |
|  | 18.85 | 27.90 | 9.05 | 0.229 | 0.8532918 |
|  | 18.78 | 27.90 | 9.12 | 0.299 | 0.8128782 |
| Normal-2 | 18.96 | 27.56 | 8.60 | -0.221 | 1.1656310 |
|  | 18.87 | 27.52 | 8.65 | -0.171 | 1.1259253 |
|  | 18.88 | 27.43 | 8.55 | -0.271 | 1.2067368 |
| Normal-3 | 19.70 | 28.43 | 8.73 | -0.091 | 1.0651902 |
|  | 19.65 | 28.51 | 8.86 | 0.039 | 0.9734043 |
|  | 19.75 | 28.41 | 8.66 | -0.161 | 1.1181480 |
| Oligoasthenospermia-1 | 19.57 | 27.13 | 7.56 | -1.261 | 2.3968026 |
|  | 19.61 | 27.36 | 7.75 | -1.071 | 2.1010509 |
|  | 19.45 | 27.57 | 8.12 | -0.701 | 1.6257564 |
| Oligoasthenospermia-2 | 18.85 | 26.89 | 8.04 | -0.781 | 1.7184538 |
|  | 18.84 | 27.00 | 8.16 | -0.661 | 1.5813000 |
|  | 18.80 | 27.10 | 8.30 | -0.521 | 1.4350600 |
| Oligoasthenospermia-3 | 18.70 | 26.94 | 8.24 | -0.581 | 1.4960010 |
|  | 18.76 | 27.09 | 8.33 | -0.491 | 1.4055269 |
|  | 18.66 | 27.07 | 8.41 | -0.411 | 1.3297095 |
| Schisandrin B-1 | 21.15 | 29.72 | 8.57 | -0.251 | 1.1901233 |
|  | 21.40 | 29.62 | 8.22 | -0.601 | 1.5168844 |
|  | 21.37 | 29.67 | 8.30 | -0.521 | 1.4350600 |
| Schisandrin B-2 | 20.84 | 29.49 | 8.65 | -0.171 | 1.1259253 |
|  | 20.73 | 29.75 | 9.02 | 0.199 | 0.8712213 |
|  | 20.79 | 28.92 | 8.13 | -0.691 | 1.6145265 |
| Schisandrin B-3 | 20.85 | 29.14 | 8.29 | -0.531 | 1.4450417 |
|  | 20.84 | 28.97 | 8.13 | -0.691 | 1.6145265 |
|  | 20.80 | 28.94 | 8.14 | -0.681 | 1.6033741 |

**Supply Dataset S11.**

| Sample | Analyte | Standard curve | r | Range (ng/mL) | LLoQ (ng/mL) |
| --- | --- | --- | --- | --- | --- |
| Plasma | Schisandrin B | $y = 8.03454x - 0.35313$ | 0.9997 | 0.065–26.087 | 0.065 |
| Testis | Schisandrin B | $y = 1.53705x + 0.177647$ | 0.9987 | 0.1–30.00 | 0.1 |

**Supply Dataset S12.**
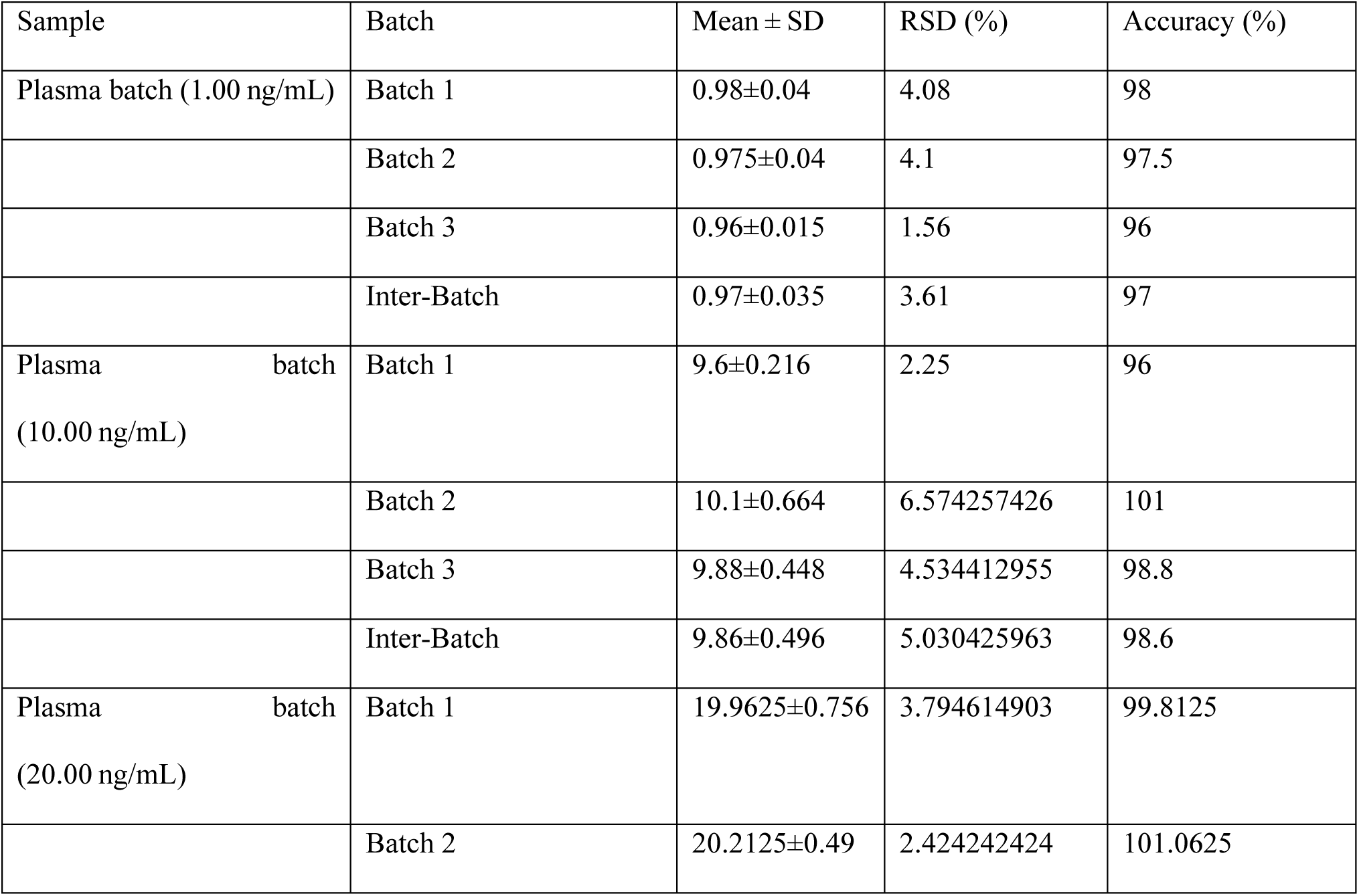
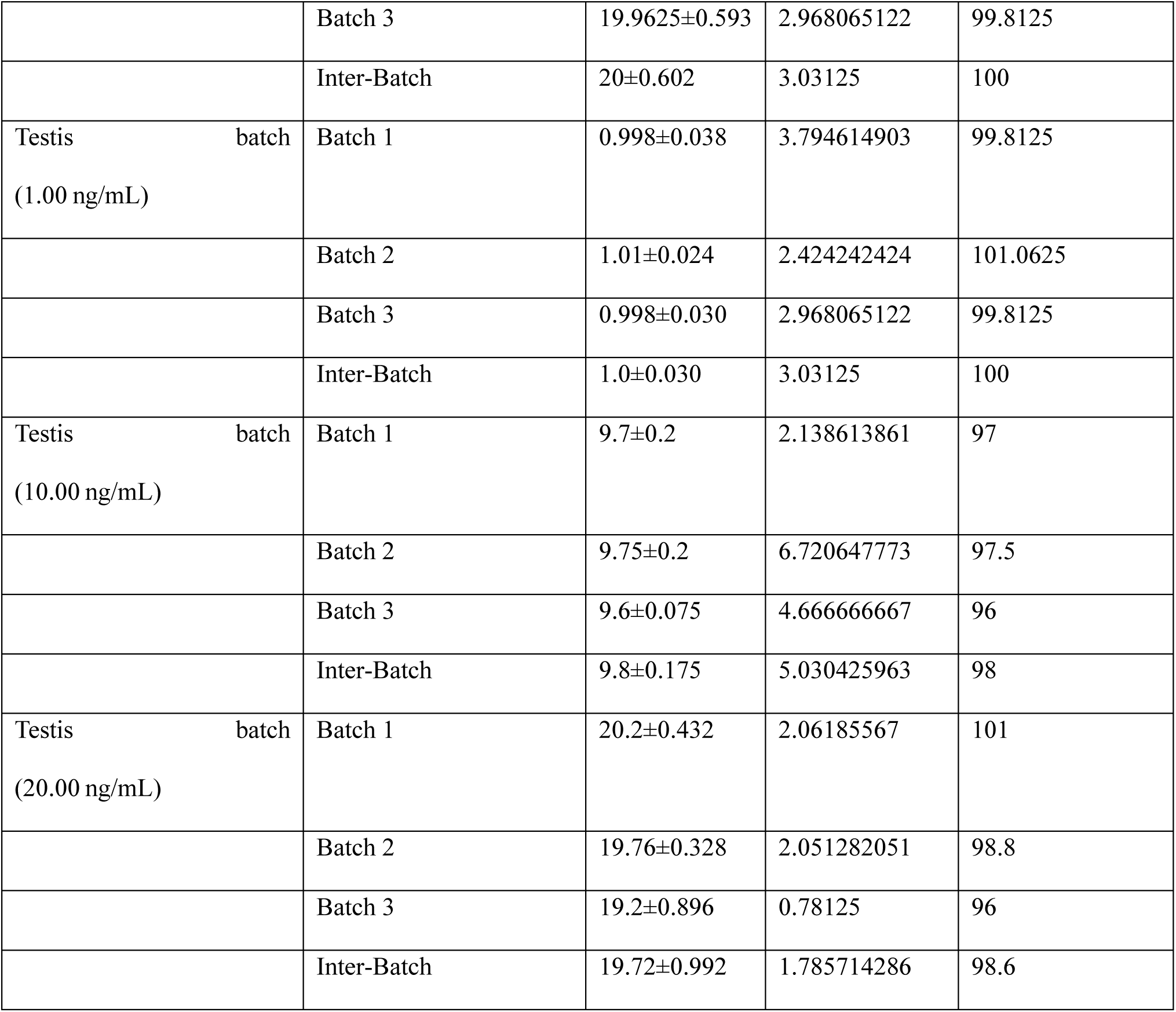

| Sample | Batch | Mean ± SD | RSD (%) | Accuracy (%) |
| --- | --- | --- | --- | --- |
| Plasma batch (1.00 ng/mL) | Batch 1 | 0.98±0.04 | 4.08 | 98 |
|  | Batch 2 | 0.975±0.04 | 4.1 | 97.5 |
|  | Batch 3 | 0.96±0.015 | 1.56 | 96 |
|  | Inter-Batch | 0.97±0.035 | 3.61 | 97 |
| Plasma batch (10.00 ng/mL) | Batch 1 | 9.6±0.216 | 2.25 | 96 |
|  | Batch 2 | 10.1±0.664 | 6.574257426 | 101 |
|  | Batch 3 | 9.88±0.448 | 4.534412955 | 98.8 |
|  | Inter-Batch | 9.86±0.496 | 5.030425963 | 98.6 |
| Plasma batch (20.00 ng/mL) | Batch 1 | 19.9625±0.756 | 3.794614903 | 99.8125 |
|  | Batch 2 | 20.2125±0.49 | 2.424242424 | 101.0625 |
|  | Batch 3 | 19.9625±0.593 | 2.968065122 | 99.8125 |
|  | Inter-Batch | 20±0.602 | 3.03125 | 100 |
| Testis batch<br>(1.00 ng/mL) | Batch 1 | 0.998±0.038 | 3.794614903 | 99.8125 |
|  | Batch 2 | 1.01±0.024 | 2.424242424 | 101.0625 |
|  | Batch 3 | 0.998±0.030 | 2.968065122 | 99.8125 |
|  | Inter-Batch | 1.0±0.030 | 3.03125 | 100 |
| Testis batch<br>(10.00 ng/mL) | Batch 1 | 9.7±0.2 | 2.138613861 | 97 |
|  | Batch 2 | 9.75±0.2 | 6.720647773 | 97.5 |
|  | Batch 3 | 9.6±0.075 | 4.666666667 | 96 |
|  | Inter-Batch | 9.8±0.175 | 5.030425963 | 98 |
| Testis batch<br>(20.00 ng/mL) | Batch 1 | 20.2±0.432 | 2.06185567 | 101 |
|  | Batch 2 | 19.76±0.328 | 2.051282051 | 98.8 |
|  | Batch 3 | 19.2±0.896 | 0.78125 | 96 |
|  | Inter-Batch | 19.72±0.992 | 1.785714286 | 98.6 |

**Supply Dataset S13.**

| Sample | Stability | Mean ± SD | RSD (%) | Accuracy (%) |
| --- | --- | --- | --- | --- |
| Plasma sample (1.00 ng/mL) | Three freeze–thaw cycles(-20 to 25 °C) | 0.98±0.04 | 7.853403141 | 95.5 |
| | Room temperature (25 °C) for 24 h | 0.975±0.04 | 3.092783505 | 97 |
| | Storage at –80 °C for 1 month | 0.96±0.015 | 2.162162162 | 92.5 |
| Plasma sample 10.00 ng/mL) | Three freeze–thaw cycles(-20 to 25 °C) | 9.6±0.216 | 3.373015873 | 100.8 |
| | Room temperature (25 °C) for 24 h | 10.1±0.664 | 1.206543967 | 97.8 |
| | Storage at –80 °C for 1 month | 9.88±0.448 | 3.747534517 | 101.4 |
| Plasma sample (20.00 ng/mL) | Three freeze–thaw cycles(-20 to 25 °C) | 19.9625±0.756 | 1.245880026 | 94.8125 |
| | Room temperature (25 °C) for 24 h | 20.2125±0.49 | 0.718484651 | 95.6875 |
| | Storage at –80 °C for 1 month | 19.9625±0.593 | 3.364369957 | 100.6875 |
| Testis sample (1.00 ng/mL) | Three freeze–thaw cycles(-20 to 25 °C) | 0.998±0.038 | 1.245880026 | 94.8125 |
| | Room temperature (25 °C) for 24 h | 1.01±0.024 | 0.718484651 | 95.6875 |
| | Storage at –80 °C for 1 month | 0.998±0.030 | 3.364369957 | 100.6875 |
| Testis sample (10.00 ng/mL) | Three freeze–thaw cycles(-20 to 25 °C) | 9.7±0.2 | 89.0052356 | 95.5 |
| | Room temperature (25 °C) for 24 h | 9.75±0.2 | 30.41237113 | 97 |
| | Storage at –80 °C for 1 month | 9.6±0.075 | 102.7027027 | 92.5 |
| Testis sample (20.00 ng/mL) | Three freeze–thaw cycles(-20 to 25 °C) | 20.2±0.432 | 3.373015873 | 100.8 |
|  | Room temperature (25 °C) for 24 h | 19.76±0.328 | 1.206543967 | 97.8 |
|  | Storage at –80 °C for 1 month | 19.2±0.896 | 3.747534517 | 101.4 |

**Supply Dataset S14.**

| Sample | Thepretical<br>concentration<br>(ng/mL) | Recovery (%) | RSD (%) |
| --- | --- | --- | --- |
| Plasma<br>sample | 1 | 85.5±4.87 | 5.13 |
|  | 10 | 88.9±3.68 | 3.29 |
|  | 20 | 86.7±3.33 | 2.37 |
| Testis<br>sample | 1 | 89.9±3.13 | 3.26 |
|  | 10 | 93.1±3.87 | 4.06 |
|  | 20 | 90.33±1.79 | 1.91 |

**Supply Dataset S15.**
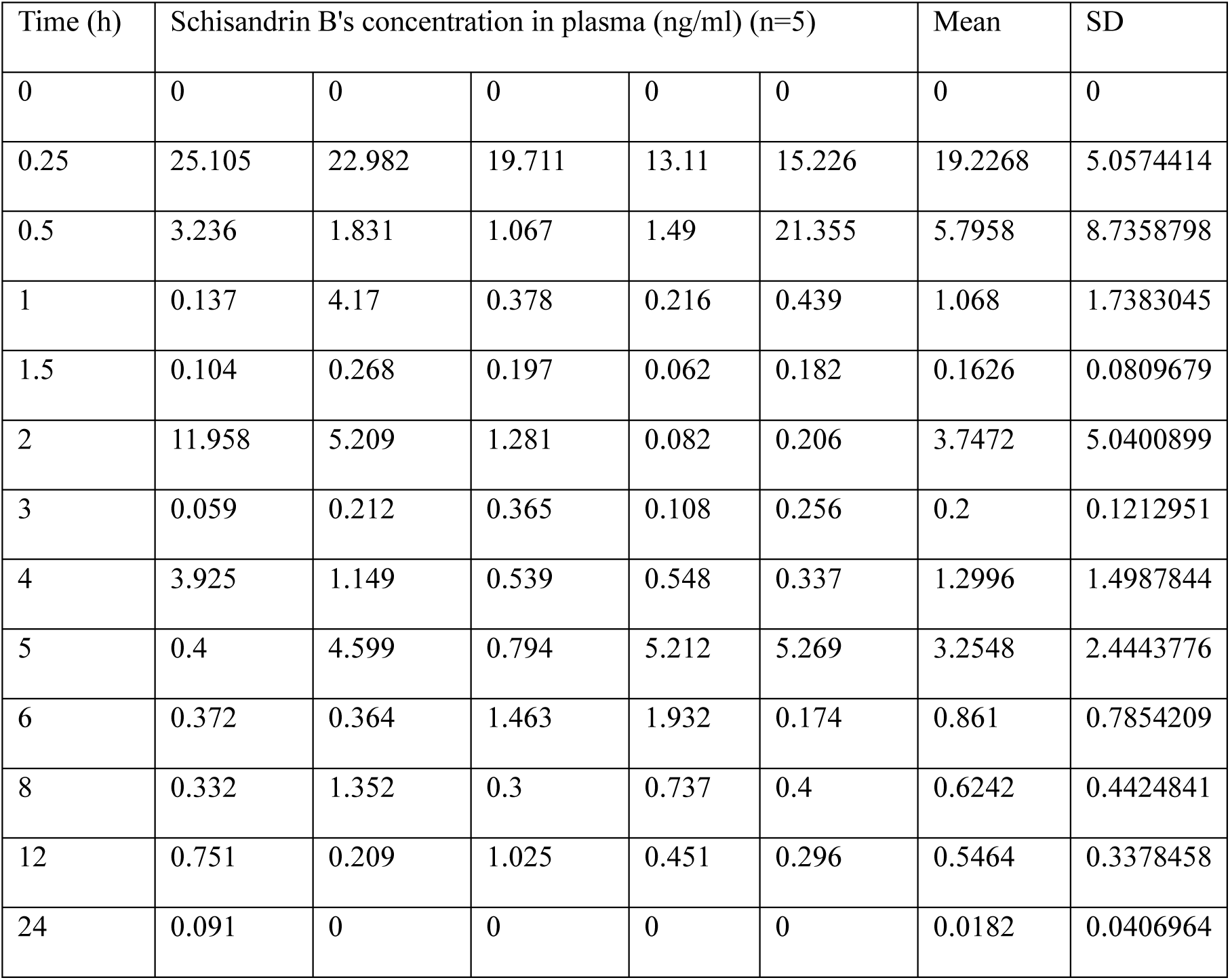

**Supply Dataset S16.**

| Time (h) | Schisandrin B's concentration in testis (ng/ml) (n=5) |  |  |  |  | Mean | SD |
| --- | --- | --- | --- | --- | --- | --- | --- |
| 0 | 0 | 0 | 0 | 0 | 0 | 0 | 0 |
| 0.25 | 4.258 | 4.97 | 3.535 | 8.055 | 5.127 | 5.189 | 1.7225155 |
| 0.5 | 2.701 | 6.043 | 2.862 | 1.791 | 4.673 | 3.614 | 1.7134705 |
| 1 | 2.498 | 2.438 | 2.453 | 3.729 | 3.509 | 2.9254 | 0.6383089 |
| 1.5 | 3.108 | 1.763 | 2.455 | 2.281 | 2.859 | 2.4932 | 0.5223669 |
| 2 | 5.192 | 1.722 | 4.218 | 13.457 | 1.679 | 5.2536 | 4.837923 |
| 3 | 6.94 | 1.505 | 2.304 | 2.925 | 1.345 | 3.0038 | 2.2906123 |
| 4 | 5.188 | 0.986 | 2.293 | 0.785 | 1.241 | 2.0986 | 1.8221612 |
| 5 | 1.309 | 1.269 | 0.922 | 0.759 | 0.172 | 0.8862 | 0.4617464 |
| 6 | 0.495 | 0.573 | 0.289 | 0.175 | 0 | 0.3064 | 0.2335033 |
| 8 | 0.117 | 0.647 | 0.193 | 0 | 0 | 0.1914 | 0.2675711 |
| 12 | 0 | 0.353 | 0.108 | 0 | 0 | 0.0922 | 0.1531085 |
| 24 | 0 | 0 | 0 | 0 | 0 | 0 | 0 |

**Supply Dataset S17.**
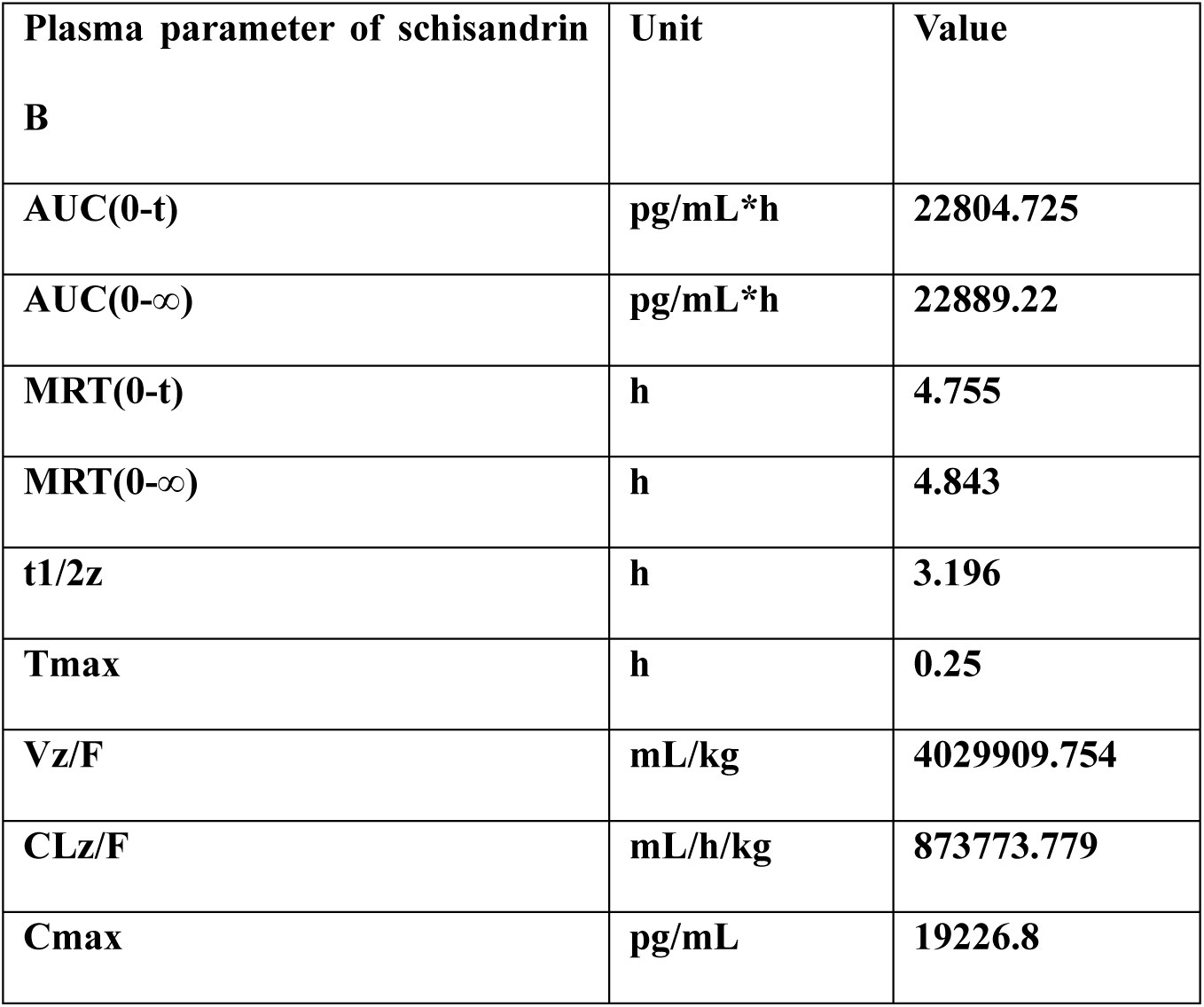

**Supply Dataset S18.**

| Testicular tissues parameter of schisandrin B | Unit | Value |
| --- | --- | --- |
| AUC(0-t) | pg/mL*h | 17062 |
| AUC(0-∞) | pg/mL*h | 17104.88 |
| MRT(0-t) | h | 2.945 |
| MRT(0-∞) | h | 3.011 |
| t1/2z | h | 3.506 |
| Tmax | h | 2 |
| Vz/F | mL/kg | 5915869.487 |
| CLz/F | mL/h/kg | 1169256.937 |
| Cmax | pg/mL | 5253.6 |

## References

1. World Health Organization. Towards more objectivity in diagnosis and management of male fertility. Int J Androl. 1987;(7): 1–53.

2. Dohle GR, Colpi GM, Hargreave TB, Papp GK, Jungwirth A, and Weidner W. EAU guidelines on male infertility. Eur Urol. 2005;48(5):703–11.

3. Boivin J, Bunting L, Collins JA, and Nygren KG. International estimates of infertility prevalence and treatment-seeking: potential need and demand for infertility medical care. *Hum Reprod (Oxford*, England). 2007;22(6):1506–12.

4. Matzuk MM, and Lamb DJ. The biology of infertility: research advances and clinical challenges. Nat Med. 2008;14(11):1197–213.

5. Hayashi K, de Sousa Lopes SM, and Surani MA. Germ cell specification in mice. Science. 2007;316(5823):394–6.

6. Soini S, Ibarreta D, Anastasiadou V, Ayme S, Braga S, Cornel M, et al. The interface between assisted reproductive technologies and genetics: technical, social, ethical and legal issues. Eur J Hum Genet. 2006;14(5):588–645.

7. Zou D, Wang J, Zhang B, Xie S, Wang Q, Xu K, et al. Analysis of Chemical Constituents in Wuzi-Yanzong-Wan by UPLC-ESI-LTQ-Orbitrap-MS. Molecules. 2015;20(12):21373–404.

8. Lamb J, Crawford ED, Peck D, Modell JW, Blat IC, Wrobel MJ, et al. The Connectivity Map: using gene-expression signatures to connect small molecules, genes, and disease. Science. 2006;313(5795):1929–35.

9. Hughes TR, Marton MJ, Jones AR, Roberts CJ, Stoughton R, Armour CD, et al. Functional discovery via a compendium of expression profiles. Cell. 2000;102(1):109–26.

10. Dyle MC, Ebert SM, Cook DP, Kunkel SD, Fox DK, Bongers KS, et al. Systems-based discovery of tomatidine as a natural small molecule inhibitor of skeletal muscle atrophy. J Biol Chem. 2014;289(21):14913–24.

11. Corson TW, and Crews CM. Molecular understanding and modern application of traditional medicines: triumphs and trials. Cell. 2007;130(5):769–74.

12. Brinster RL, and Avarbock MR. Germline transmission of donor haplotype following spermatogonial transplantation. Proc Natl Acad Sci. USA. 1994;91(24):11303–7.

13. Wang X, Ding Q, Zhang Y, Wang H, Ma L, and Xie X. Two allogeneic descendents derived from the high-dose busulfan-treated infertile mouse model after freeze-thawed spermatogonial stem cell transplantation. Fertil Steril. 2008;90(4 Suppl):1538–49.

14. World Health Organization. WHO laboratory manual for the Examination and processing of human semen. Fifth Edition. Geneva: WHO Press;2010.

15. Datta-Mannan A, Yaden B, Krishnan V, Jones BE, and Croy JE. An engineered human follistatin variant: insights into the pharmacokinetic and pharmocodynamic relationships of a novel molecule with broad therapeutic potential. J Pharmaco Exp Ther. 2013;344(3):616–23.

16. Mather JP, Roberts PE, and Krummen LA. Follistatin modulates activin activity in a cell- and tissue-specific manner. Endocrinology. 1993;132(6):2732–4.

17. Jones KL, Mansell A, Patella S, Scott BJ, Hedger MP, de Kretser DM, et al. Activin A is a critical component of the inflammatory response, and its binding protein, follistatin, reduces mortality in endotoxemia.Proc Natl Acad Sci. USA. 2007;104(41):16239–44.

18. Chen W, and Ten Dijke P. Immunoregulation by members of the TGFbeta superfamily. Nat Rev Immunol. 2016;16(12):723–40.

19. Nakamura T, Takio K, Eto Y, Shibai H, Titani K, and Sugino H. Activin-binding protein from rat ovary is follistatin. Science. 1990;247(4944):836–8.

20. Okuma Y, O’Connor AE, Hayashi T, Loveland KL, de Kretser DM, and Hedger MP. Regulated production of activin A and inhibin B throughout the cycle of the seminiferous epithelium in the rat. J Endocrinol. 2006;190(2):331–40.

21. Mendis SH, Meachem SJ, Sarraj MA, and Loveland KL. Activin A balances Sertoli and germ cell proliferation in the fetal mouse testis. Biol Reprod. 2011;84(2):379–91.

22. Tanimoto Y, Tanimoto K, Sugiyama F, Horiguchi H, Murakami K, Yagami K, et al. Male sterility in transgenic mice expressing activin betaA subunit gene in testis. Biochem Biophys Res Commun. 1999;259(3):699–705.

23. Winnall WR, Wu H, Sarraj MA, Rogers PA, de Kretser DM, Girling JE, et al. Expression patterns of activin, inhibin and follistatin variants in the adult male mouse reproductive tract suggest important roles in the epididymis and vas deferens. Reprod Fertil Dev. 2013;25(3):570–80.

24. Wijayarathna R, and de Kretser DM. Activins in reproductive biology and beyond. Hum Reprod Update. 2016;22(3):342–57.

25. Nicolas N, Muir JA, Hayward S, Chen JL, Stanton PG, Gregorevic P, et al. Induction of experimental autoimmune orchitis in mice: responses to elevated circulating levels of the activin-binding protein, follistatin. Reproduction. 2017;154(3):293–305.

26. Singh SS. Preclinical pharmacokinetics: an approach towards safer and efficacious drugs. Curr Drug Metab. 2006;7(2):165–82.

27. Iga K. [Verification of Pharmacokinetic Approaches in Prior Drug Development]. Yakugaku Zasshi. 2019;139(3):437–60.

28. de Campos ML, Padilha EC, and Peccinini RG. A review of pharmacokinetic parameters of metabolites and prodrugs. Drug Metab Lett. 2014;7(2):105–16.

29. US Food and Drug Administration. Guidance for Industry, Bioequivalence: Blood Level Bioequivalence Study. Silver Spring, MD: FDA; 2014.

30. European Medicines Agency. Guideline on Bioanalytical Method Validation, EMEA/CHMP/ EWP/192217/2009 Amsterdam: Committee for Medicinal Products for Human Use, European Medicines Agency; 2011.

